# Activation of SPARDA defense system by filament assembly reveals beta-relay signaling mechanism widespread in prokaryotic Argonautes

**DOI:** 10.1101/2025.04.08.647717

**Authors:** Edvinas Jurgelaitis, Evelina Zagorskaitė, Aurimas Kopūstas, Simonas Asmontas, Elena Manakova, Indrė Dalgėdienė, Ugnė Tylenytė, Arunas Silanskas, Paulius Toliusis, Algirdas Grybauskas, Marijonas Tutkus, Česlovas Venclovas, Mindaugas Zaremba

**Affiliations:** Institute of Biotechnology, Life Sciences Center, Vilnius University, Saulėtekio av. 7, LT-10257, Vilnius, Lithuania; Department of Molecular Compound Physics, Center for Physical Sciences and Technology, Saulėtekio av. 3, LT-10257, Vilnius, Lithuania

**Keywords:** prokaryotic Argonaute, DREN domain, SPARDA, X-ray, cryo-EM, single-molecule, filament

## Abstract

Present in all three domains of life, Argonaute (Ago) proteins use short oligonucleotides as guides to recognize complementary nucleic acid targets and are involved in RNA silencing (eukaryotes) or host defense against invading DNA (prokaryotes). Here, we show that the SPARDA (short prokaryotic Argonaute, DNase associated) system functions in anti-plasmid defense. Upon detection of invading plasmid DNA, it degrades both the invader and host DNA, inducing cell death and preventing further spread of the invader. Upon activation, the recognition signal of the bound guide/target duplex is relayed to other functional SPARDA sites through a structural region, which we termed ‘beta-relay’. The associated dramatic conformational changes trigger the formation of a filament, in which the DREN nuclease domains form tetramers poised to cleave dsDNA. Furthermore, we identified the presence of beta-relay in all pAgo clades, providing new insights into the structural mechanisms of pAgo proteins.

## INTRODUCTION

Argonaute (Ago) proteins are present in all three domains of life (bacteria, archaea, and eukaryotes) and use short (15-35 nt) oligonucleotides as guides to bind complementary nucleic acid targets.^1–10^ Eukaryotic Argonautes (eAgos) form the functional core of the RNA silencing machinery responsible for regulating gene expression, silencing mobile genome elements, and defense against viruses.^11,12^ Prokaryotic Argonautes (pAgos), being genetically and structurally significantly more diverse than their eukaryotic homologs,^13^ play a role in host defense against invading DNA such as plasmids and viruses.^3,14–16^ pAgos are also implicated in genome decatenation^17^ and stimulation of homologous recombination.^18,19^ Traditionally, based on their domain organization, Agos are divided into long and short Agos.^13^ Long Agos (including eAgos) are composed of 4 major functional domains (N – strand separation, PAZ – guide strand 3′-end binding, MID – guide strand 5′-end binding, and PIWI – nucleic acid cleavage) and two structural linker domains L1 and L2.^13^ Phylogenetically long pAgos can be further subdivided into long-A and long-B groups, where all long-B pAgos lack the catalytic DEDX tetrad in the PIWI domain.^13^ To perform their biological functions, catalytically inactive long-B pAgos rely on various associated effector proteins encoded in the vicinity.^20^ Compared to catalytically active long-A pAgos bearing a canonical PAZ domain, catalytically inactive long-B pAgos may have various levels of PAZ domain reduction, including its complete loss.^21^ The majority of pAgos (∼60%) belong to the group of highly divergent short pAgos, which contain only the MID and inactive PIWI domains and completely lack the PAZ domain.^13^ However, short pAgos typically form stable complexes with the associated effector proteins containing a domain historically named APAZ (Analog of PAZ^22^), which, in fact, is structurally homologous to the N, L1, and L2 domains of long pAgos.^23^ APAZ domains are often N-terminally fused to Sir2 (Silent informator regulator 2) or TIR (Toll-Interleukin-1 Receptor) effectors or nucleases of Mrr or DREN families.^13,23^ Therefore, heterodimeric Effector-APAZ/Agos or single chain Effector-APAZ-Agos resemble long-B PAZ-less pAgos containing a fused effector domain. Upon RNA guide-mediated detection of invading DNA by short pAgos functional complexes, the associated effector domains trigger a defense response. Sir2 and TIR domains in SPARSA (Short Prokaryotic Argonaute Sir2-APAZ) and SPARTA (Short Prokaryotic Argonaute TIR-APAZ) systems, respectively, deplete cellular NAD^+^, DREN domains in SPARDA (Short Prokaryotic Argonaute, DNase associated) degrade both the invader and the host genome.^3,14,15^ This ultimately kills the host cell, preventing the spread of the invader within the population.^3,14,15^

To date, detailed structural mechanisms of action have been proposed only for SPARTA and SPARSA short pAgo systems employing NADase effector domain, but not SPARDA or other short pAgos possessing nuclease effector domain.^23–34^ In both SPARTA and SPARSA, the apo form corresponds to a binary monomer composed of the Effector-APAZ and Ago proteins. The binding of an RNA guide and a complementary ssDNA target induces conformational changes that lead to the dimerization of the complexes *via* the Ago-Ago surface, followed by tetramerization *via* the TIR or Sir2 domains. In SPARTA, the tetramer of TIR domains contains two composite NADase active sites, while in SPARSA, all four Sir2 domains have catalytically competent active sites. Like CRISPR Cas12, activated SPARDA exhibits collateral nuclease activity, demonstrating potential in nucleic acid (NA) diagnostics.^14,35^ However, unlike Cas12 systems, SPARDA uses short RNA guides (∼20 nt) for NA detection and does not require a PAM sequence for target recognition. Therefore, elucidating the detailed structural mechanism of SPARDA is important for fundamental knowledge and expanding its potential for biotechnological applications.

Here, we present detailed functional, structural, biochemical, and single-molecule studies of two homologous SPARDA systems. Our results reveal SPARDA activation *via* filament assembly and disclose an allosteric signal transmission mechanism common for all clades of pAgos.

## RESULTS

### SPARDA acts as an anti-plasmid defense system, inducing cell death via nonspecific DNA degradation

To perform functional, biochemical, and structural studies of SPARDA systems, we selected two homologous yet uncharacterized SPARDAs from mesophilic *Xanthobacter autotrophicus* strain Py2 (XauSPARDA) and thermophilic *Enhydrobacter aerosaccus* (EaeSPARDA) belonging to the S2B clade of the short pAgos (Figures 1A and 1B). They are both composed of a catalytically inactive short pAgo and a putative DREN-APAZ effector bearing DUF4365/DREN (**D**NA and **R**NA **e**ffector **n**uclease) domain that belongs to the PD-(D/E)XK superfamily of metal-dependent nucleases (Figures 1A, S1A and S1B).^14^ XauSPARDA and EaeSPARDA genes were heterologously expressed in *E. coli* cells by cloning them into pBAD expression vectors under a P_BAD_ promoter (Figure 1C, Table S1) and challenged them with phages or plasmids. Unlike the previously characterized Nba or CmeSPARDA,^14^ which showed defense against some phages, we did not observe robust protection of *E. coli* cells against 12 selected phages under our experimental conditions (Figures S2A and S2B). However, both overexpressed WT Xau and EaeSPARDA, but not the catalytic mutants of the DREN-APAZ (the active site E37A mutation within the DREN domain) or the XauAgo mutant protein (a bulky 29 aa HSH-tag at the C-terminus that is important for nucleic acid binding in other Agos), reduced the viability of *E. coli* (Figure S2C). However, the expression level of the mutants was similar to that of the WT SPARDAs (Figure S2D). Thus, both functional DREN-APAZ and the Ago proteins are required to induce cell death by the SPARDA system.

**Figure 1.**
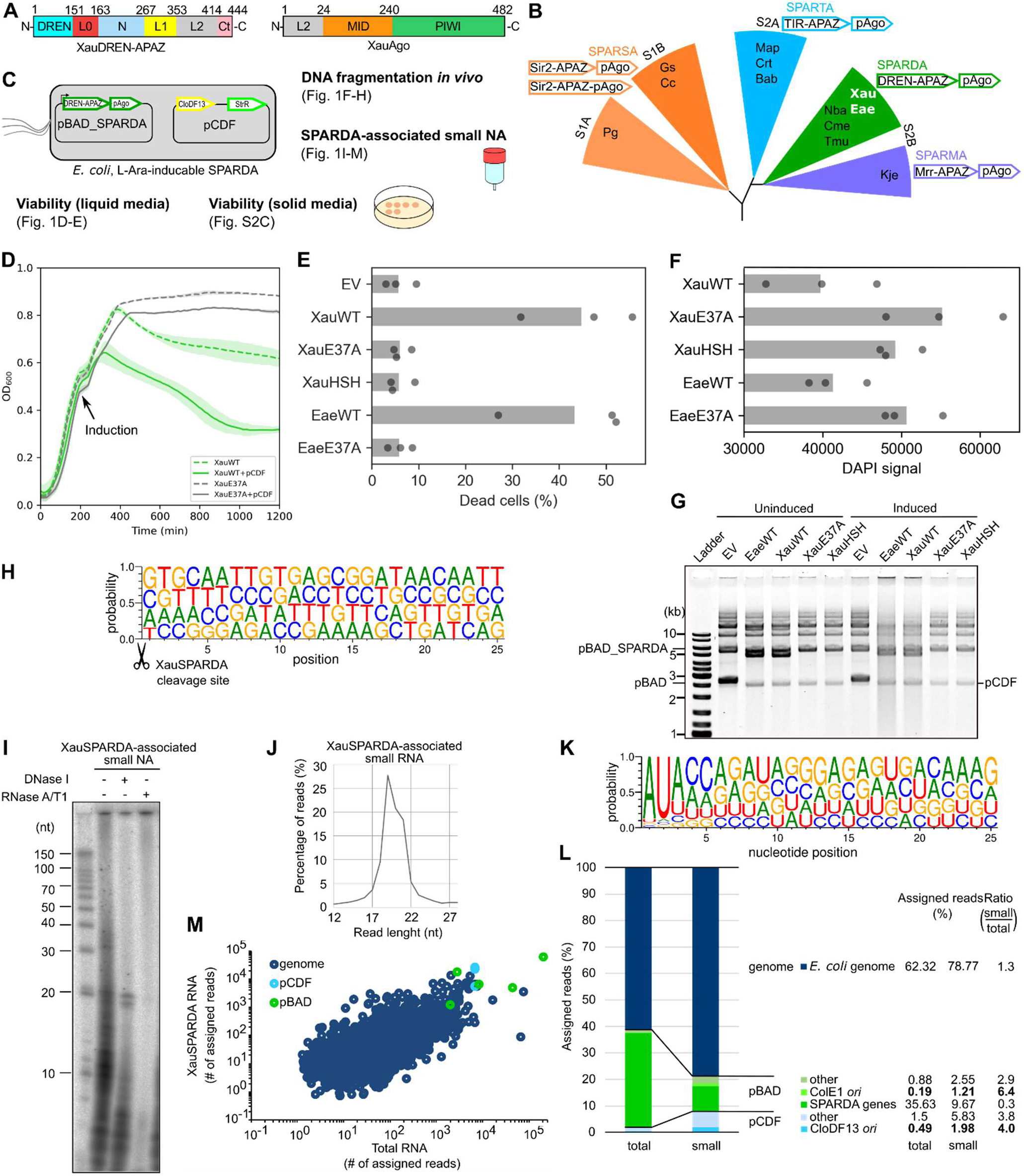
SPARDA is an anti-plasmid defense system. **(A)** Schematic diagram of XauSPARDA domain organization. **(B)** Schematic unrooted phylogenetic tree of short pAgo systems (based on ^14^) with previously studied examples^3,14,15,23,35,38^ and two SPARDA systems (Xau and Eae from *Xanthobacter autotrophicus* strain Py2 and *Enhydrobacter aerosaccus*, respectively) addressed in this study. Schematic operon structure(s) for each clade is also provided. **(C)** Overview of *in vivo* assays, where *E. coli* cells bear the XauSPARDA system cloned into pBAD vector under L-Ara-inducible promoter and the second pCDF plasmid. **(D)** Growth curves of *E. coli* cultures expressing XauSPARDA with and without pCDF plasmid. Data are shown as mean ± SD across three biological replicates. **(E)** Viability of *E. coli* cells expressing SPARDA systems in the presence of pCDF plasmid. Dead cells were counted by flow cytometry using PI staining. The average of three biological replicates is shown in bars. EV - empty vector. **(F)** An integrated DAPI signal of stained *E. coli* cells expressing SPARDA systems in the presence of pCDF plasmid was measured by flow cytometry. The average of three biological replicates is shown in bars. **(G)** DNA fragmentation within *E. coli* cells with (un)induced SPARDA systems. Supercoiled forms of pBAD, pBAD_SPARDA, and pCDF plasmids are indicated in the figure; other forms of these plasmids and their concatemers are also present in the gel. **(H)** Nucleotide distribution at the XauSPARDA cleavage site (indicated by scissors). **(I)** Identification of XauSPARDA-bound nucleic acids. Nucleic acids copurified with XauSPARDA were radiolabeled, treated with DNase I or RNase A/T1, and resolved on a denaturing polyacrylamide gel. **(J)** Length distribution of small RNA copurified with XauSPARDA as determined by sequencing. **(K)** Nucleotide bias of small RNAs copurified with XauSPARDA. **(L)** Percentages of total RNA (*E. coli* expressing XauSPARDA) and small RNA (copurified with XauSPARDA) reads that align to specific genomic or plasmid sequences. **(M)** Correlation between XauSPARDA-associated small RNA and *E. coli*-extracted total RNA sequences. See also Figures S1, S2 and S3.

Previous studies of long-B pAgos and SPARSA defense systems revealed that they are activated by a plasmid (e.g., pCDF) in the cell.^15,20^ Indeed, the pCDF plasmid (in addition to the expression pBAD vector) increased the WT Xau and EaeSPARDA cytotoxicity by ∼8.5×10^3^ and ∼1.6×10^4^-fold, respectively (Figures 1C and S2C). The cell death-triggering effect of pCDF plasmid was also observed in a liquid medium by recording the growth curves of WT Xau and EaeSPARDA cultures (Figures 1D and S2E). According to the flow cytometry using *E. coli* cells staining with propidium iodide (PI), which penetrates dead cells with a damaged membrane, both WT SPARDAs, but not their mutants, caused significant *E. coli* cell death (up to ∼45%) after 2h induction of their expression in the presence of pCDF plasmid (Figure 1E). Meanwhile, the signal of the same DAPI-stained *E. coli* cells bearing WT Xau and EaeSPARDA decreased compared to their functionally inactive mutants, indicating a possible intracellular DNA degradation (Figures 1F and S2F). Next, using a plasmid purification kit, we purified DNA from the same *E. coli* cells (2h after induction of SPARDAs) and analyzed its integrity in an agarose gel (Figure 1G). Nonspecific DNA fragmentation was observed only when DNA was purified from the cells expressing the WT Xau and EaeSPARDA systems. To determine the source of the DNA fragments (plasmid or genomic DNA) and positions of WT XauSPARDA introduced double-stranded DNA breaks (DSB), we adapted our CRISPR/Cas9 off-target detection method CROFT-seq.^36^ Sequencing analysis showed that the WT XauSPARDA-generated DNA fragments originated from both *E. coli* genomic DNA (∼59% of aligned reads) and plasmid DNA (pCDF and pBAD-XauSPARDA, ∼27% and ∼14%, respectively) (Data S1). Assuming that there could be ∼40 copies of the pCDF and pBAD plasmids in a cell,^37^ the activated XauSPARDA could proportionally generate only ∼3% and ∼6% of DNA fragments from them, respectively, suggesting there might be somewhat specific plasmid targeting (more pronounced for the pCDF plasmid). DSB sites are evenly distributed throughout the genomes, not forming localized clusters, indicating that XauSPARDA is not targeting specific genes. Analysis of the DNA sequences adjacent to the DSB revealed no specificity or preference for any particular DNA sequences (Figure 1H). Taken together, XauSPARDA activated by pCDF plasmid nonspecifically degrades both genomic and plasmid DNA *in vivo*.

### In vivo SPARDA preferentially binds small 5’AU-RNAs originating from abundant transcripts

Previously, it was experimentally demonstrated that catalytically inactive short and long-B pAgos *in vivo* bind small RNAs originating from cellular transcripts, which they use as guides to bind DNA targets.^3,15,20^ To identify nucleic acids (NAs) bound by our SPARDAs *in vivo*, we purified the Xau and EaeSPARDA-NA complexes from *E. coli* transformed with the pBAD_SPARDA expression vector and the pCDF plasmid, extracted NAs, and analyzed them by digestion with nucleases and by sequencing. Xau and EaeSPARDA are associated with small (predominantly ∼20 nt) RNAs with 5’AU dinucleotide preference (Figures 1I-1K and S3A-S3C). Copurified small RNAs originated from all genomes: *E. coli*, pBAD_SPARDA, and pCDF plasmids (Figures 1L and S3D, Data S1). Interestingly, comparing Xau and EaeSPARDA total and small RNA-seq data, a slight enrichment (∼4-8-fold) of pCDF and pBAD transcripts from CloDF13 and ColE1 *ori* regions, respectively, among copurified small RNAs was observed in all cases (Figures 1L and S3D, Data S1). The amounts of copurified small RNAs correlate with the abundance of cellular transcripts, determined by total RNA sequencing (Spearman correlation coefficients are 0.60 and 0.61 for Xau and EaeSPARDA, respectively) (Figures 1M and S3E). However, no correlation was found between total RNA transcripts, from which small XauSPARDA-bound RNAs are formed, and DSB sites (Spearman correlation coefficient of 0.022 between RNA- and DNA-seq reads for all annotated genes), indicating that DNA targeting and cleavage sites of XauSPARDA are unrelated (Figure S3F). Thus, Xau and EaeSPARDA *in vivo* preferentially bind small 5’AU-RNAs originating from abundant cellular transcripts, with a slight enrichment of small RNAs of transcripts originating from the *ori* regions of the pCDF and pBAD plasmids.

### Apo SPARDA is a dimer of binary DREN-APAZ:Ago monomers

To characterize the Xau and EaeSPARDA systems biochemically, we coexpressed their DREN-APAZ and pAgo genes in *E. coli* and then purified the corresponding proteins by liquid chromatography (Figure S3G). Although only the effector DREN-APAZ protein contained a His-tag, both proteins copurified on the Ni^2+^-affinity and subsequent columns (Figure S3G), indicating that they form a stable complex as in other published SPAR(D/S/T)A cases.^3,14,15,35^ However, in contrast to other non-activated apo SPAR(D/S/T)As complexes (binary Effector-APAZ:Ago monomers), the SEC-MALS data showed that both apo Xau and EaeSPARDA exist as dimers of two SPARDA complexes (with DREN-APAZ_2_:Ago_2_ stoichiometry) in solution (Table S3).^3,14,15,35^ Functionally compromised XauE37A, EaeE37A, and XauHSH variants also formed a dimeric SPARDA complex, showing that the introduced mutations abolished the activity *in vivo* but did not affect the protein complex composition (Table S3). Thus, apo SPARDA forms a stable dimer of binary SPARDA complexes.

### Activated SPARDA is a nonspecific dsDNA endonuclease in vitro

Since WT Xau and EaeSPARDA degrade genomic and plasmid dsDNA *in vivo* (Figure 1G), we performed their further biochemical characterization *in vitro* using a short (42 bp long) double-stranded (ds) DNA as a substrate added *in trans* and single-stranded (ss) 17 nt DNA or RNA guides and targets. WT Xau and EaeSPARDA nonspecifically degraded the dsDNA substrate only using a guide RNA and a complementary DNA target in the presence of Mg^2+^ ions as a cofactor (Figures 2A and S3I). Since previously characterized SPARDA systems were able to hydrolyze not only dsDNA substrates *in vitro,*^14^ we studied the nucleolytic activity of our SPARDAs on ssDNA, ssRNA, DNA/RNA heteroduplex, and dsRNA substrates (Figures 2B and S3J). However, Xau and EaeSPARDA showed activity only on the dsDNA substrate. They also nonspecifically degraded long dsDNAs with various topologies: circular supercoiled pUC19 plasmid, linear phage lambda, and *E. coli* genomic DNA (Figures 2C and S3K). The SPARDA’s nuclease DREN domain bearing a canonical PD-(D/E)XK active site (Figure S1A) uses Mg^2+^ and Mn^2+^ ions as cofactors for DNA cleavage, but not Ca^2+^ and Ni^2+^ (Figure S3L). Taken together, the guide/target-bound XauSPARDA and EaeSPARDA complexes act as catalytically activated nonspecific dsDNA endonucleases *in vitro*.

**Figure 2.**
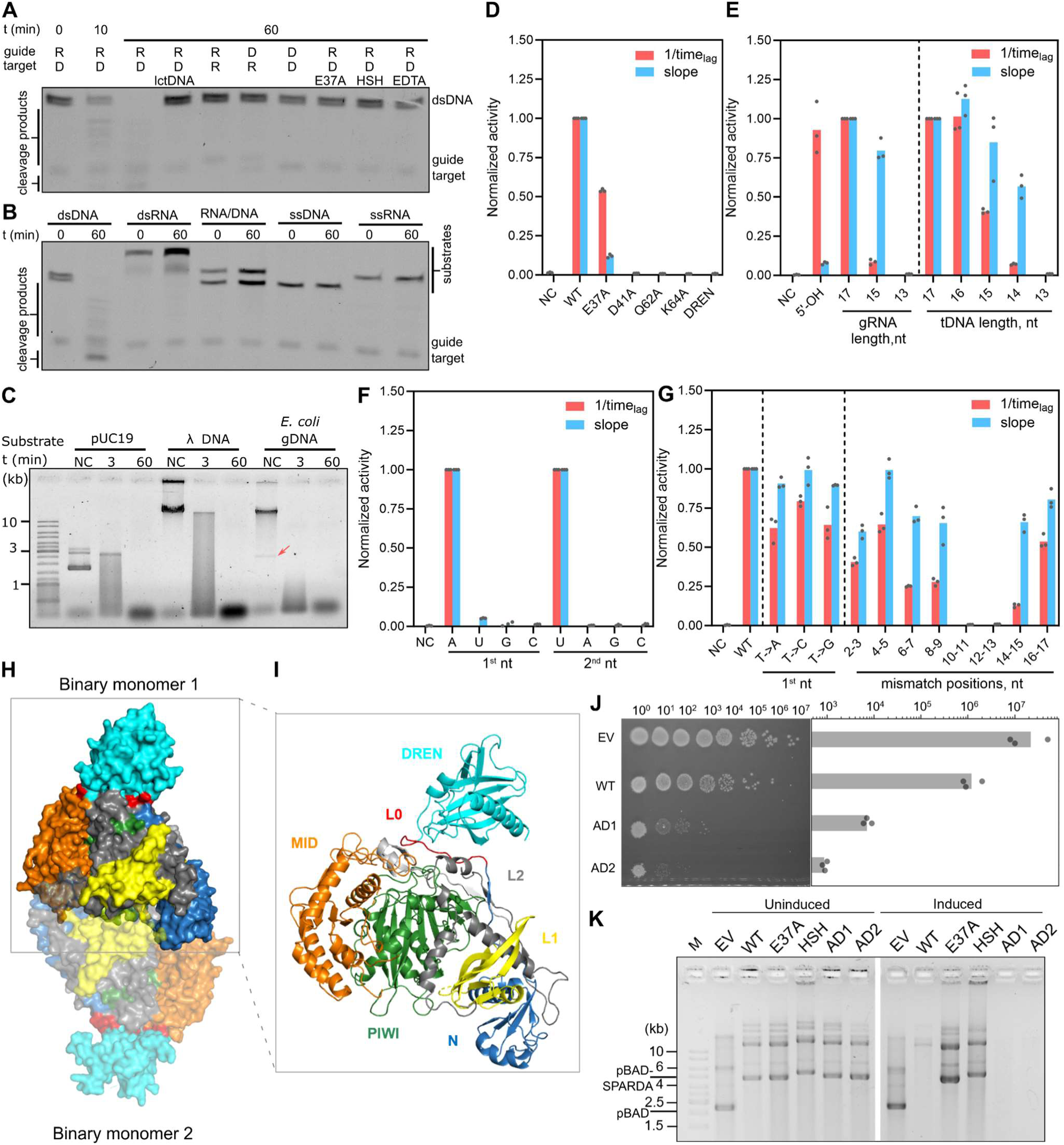
XauSPARDA acts as a nonspecific dsDNA endonuclease. **(A)** Analysis of the specificity of XauSPARDA for gRNA and tDNA with a dsDNA substrate. **(B)** Collateral cleavage activity of XauSPARDA with various short NA substrates. **(C)** Collateral cleavage activity of XauSPARDA with various long dsDNA substrates. NC (negative control) - reaction without protein. The red arrow indicates the pBAD plasmid. **(D)** Cleavage activity of XauSPARDA active center mutants. DREN - isolated DREN domain; WT - reaction with fully complementary gRNA (Mz-1800) and tDNA (Mz-1455). **(E)** Cleavage activity of XauSPARDA with gRNAs and tDNAs of various lengths. 5′-OH depicts dephosphorylated gRNA of 17 nt in length. **(F)** Effect of substitutions of the first or second 5′-nucleotide in gRNA on the cleavage activity of XauSPARDA**. (G)** Effect of single or double mismatches within tDNA on XauSPARDA activation. The numbers show a mismatch position according to 5′-gRNA. **(H)** Crystal structure of apo XauSPARDA dimer. Protein domains are colored as in Figure 1A. (**I**) XauSPARDA binary monomer **(J)** Cytotoxic effect of XauSPARDA system on *E. coli* cells expressing WT and apo dimerization mutants. EV - empty vector. Left: serial dilutions on agar plates, right: colony forming units (CFU) were counted, and the average of three biological replicates is shown in bars. **(K)** DNA fragmentation within *E. coli* cells with (un)induced SPARDA systems. Supercoiled forms of pBAD and pBAD_SPARDA plasmids are indicated in the figure; other forms of these plasmids and their concatemers are also present in the gel. See Figures S1, S3, and S4.

Next, to further characterize the *in trans* cleavage (collateral) activity of our SPARDAs in more detail, we monitored nuclease activity using a fluorophore-quencher substrate. A typical fluorescence signal-time curve initially had a lag phase followed by a signal increase. Therefore, we used two parameters to estimate SPARDA activity: 1/time_lag_, reflecting how fast the cleavage begins after the reaction start, and the slope of the linear part of the rising curve, indicating the cleavage rate (Figure S3M). Replacement of conserved catalytic residues within the XauSPARDA active site (Figure S1A) with alanine significantly reduced (E37A, also in EaeSPARDA) or completely abolished (D41A, Q62A, K64A) activity (Figures 2D and S3N). The isolated XauDREN domain shows no activity, indicating that the remaining SPARDA part is required for activation (Figure 2D).

Both Xau and EaeSPARDA require a 5′-P RNA guide (instead of 5′-OH) for their efficient activity (Figures 2E and S3N), and no activity was observed using a 13 nt long RNA guide (Figures 2E and S3N). Xau and EaeSPARDA tolerate DNA target strand shortening up to 14 and 16 nt, respectively (Figures 2E and S3N). XauSPARDA possesses high and moderate cleavage activity on the ssDNA target and the ssDNA target within an internal loop, respectively (Figure S3O). An entirely complementary dsDNA target does not activate the cleavage activity of XauSPARDA, indicating that it cannot unwind a stable DNA duplex (Figure S3O). XauSPARDA can also use a pre-annealed gRNA/tDNA duplex for its activation; however, the DNA cleavage is less efficient than using the separate RNA guide and ssDNA target (Figure S3P).

Our *in vivo* data show that Xau and EaeSPARDA prefer binding small 5′-AU-RNAs (Figures 1K and S3C) that may originate from cellular transcripts during their degradation after RNase E cleavage at its AU-rich recognition motif.^39–42^ To determine whether our SPARDAs bind the most abundant small intracellular RNAs nonspecifically or recognize the first two bases at the 5′-end of the guide, we performed experiments using RNA guides with different bases at their 1^st^ and 2^nd^ positions (Figures 2F and S3Q). Indeed, Xau and EaeSPARDA were active only with the 5′-AU-RNA guide, suggesting they have evolved to specifically bind the degradation products of RNA transcripts.^39–42^

Next, to determine how our SPARDAs tolerate changes in the complementarity of the DNA target strand, we used a set of DNA targets in which a nucleobase was varied at the 1^st^ position, and 2 nt mismatches were introduced at various positions along the entire length of the strand (Figures 2G and S3R). Most of the changes either had no effect or only slightly reduced activity. However, introducing mismatches at positions 10-13 and 10-15 in the case of Xau and EaeSPARDA, respectively, completely abolished their activity (Figures 2G and S3R).

### Structure of apo SPARDA dimer

To elucidate the molecular mechanism of action of the SPARDA system, we determined the crystal structure of the apo WT XauSPARDA (at a resolution of 3 Å) and its variant, which lacks the catalytic DREN domain (1.5 Å), and WT EaeSPARDA (2.45 Å) (Table S2). Both apo SPARDAs are dimers of two binary DREN-APAZ:Ago monomers (Figures 2H and S4A) as determined by SEC-MALS in solution (Table S3). The overall structure of the binary Xau and EaeSPARDA monomer (DREN-APAZ:Ago complex) is typical for structurally characterized SPAR(S/T)A complexes, except that effector domains occupy different positions relative to the conserved APAZ:Ago core (Figures S4A and S4B). In the XauDREN-APAZ protein, the N-terminal catalytic DREN domain is connected by the flexible linker L0 to the APAZ part that structurally corresponds to the N-L1-L2 region of long pAgos lacking the PAZ domain (Figure 2I). The XauAgo protein possesses a canonical arrangement of MID and PIWI domains, as in other pAgos and eAgos.

### Apo dimerization of XauSPARDA represses its activity

In the apo Xau and EaeSPARDA dimer, the binary DREN-APAZ:Ago monomers dimerize, locating the DREN domains on opposite sides of the complex (Figures 2H and S4A). However, such apo dimerization mode of SPARDA complexes completely differs from the activated SPAR(S/T)A dimers (see below, Figure S6B). All four protein subunits of XauSPARDA participate in apo dimerization contacts, resulting in a total contact surface of ∼2667 Å^2^ (Table S4). To determine the importance of SPARDA apo dimerization for its function, we introduced multiple mutations at the **a**po **d**imerization interface (mutants AD1 and AD2) to disrupt dimerization-relevant interactions (Figure S4C). According to our SEC-MALS results, both purified mutants are monomers in solution (Table S3). *In vitro,* these mutants were active but showed a shorter lag phase and were slightly more tolerant to some guide/target mismatches and the target strand shortening than WT XauSPARDA (Figures 2E, 2G, S4D, and S4E). Compared to WT XauSPARDA, both mutants exhibited significantly higher toxicity to *E. coli* cells, probably due to more efficient DNA fragmentation *in vivo* (Figures 2J-2K), given that all (WT and mutant) variants had similar expression levels (Figure S3H). These results indicate that apo dimerization is essential for controlling the XauSPARDA activity and increasing its fidelity to avoid nonspecific damage to the host cell.

### Activated XauSPARDA forms a filament

Next, we determined a cryo-EM structure of the activated XauSPARDA-guide/target complex with a 42 bp DNA substrate. Unexpectedly, this complex formed helical filaments whose structure was reconstituted at a resolution of 3 Å (Figures 3A-3C, S5A, and S5B). The activated XauSPARDA filament consists of “steps” of two binary APAZ:Ago monomers binding the RNA guide and DNA target heteroduplexes (Figures 3D-3F). The intrastep dimerization surface of XauSPARDA covers ∼1200 Å^2^ (Table S4) and involves the same Ago secondary structure elements as in the case of activated SPARTA dimers^23^ and slightly differs from the asymmetric dimerization reported for activated GsuSPARSA^32^ (Figure S6B). It should be emphasized that the entirely different dimerization in apo XauSPARDA is incompatible with the intrastep dimerization observed in the activated XauSPARDA (Figures S6C and S6D). Also, during the activation of XauSPARDA, when the guide and the target are bound, the apo dimer must disintegrate since it could not sterically accommodate the bound duplex (Figure S6D), and the formed activated monomers dimerize using other surfaces to assemble the filament further.

**Figure 3.**
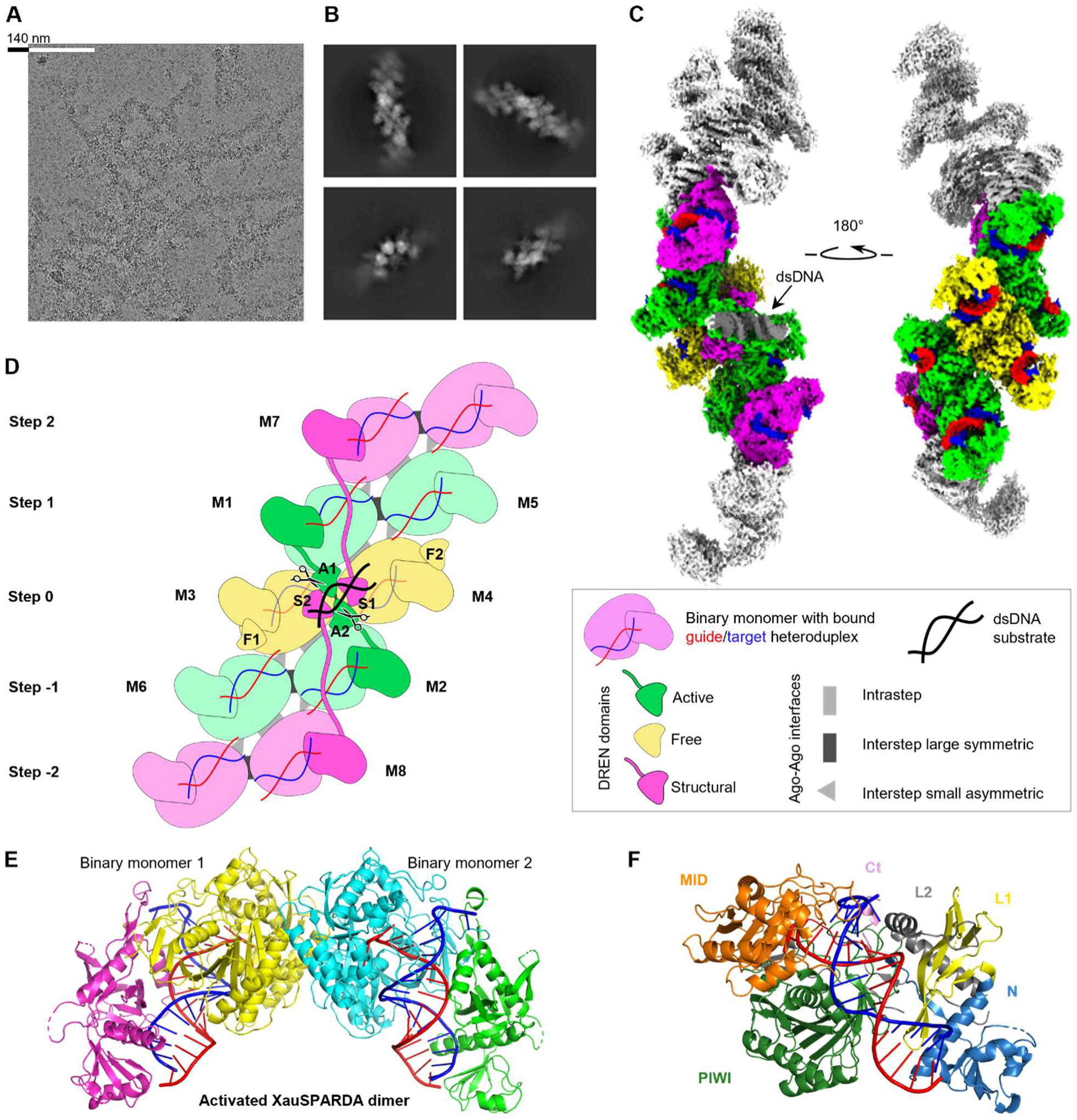
Cryo-EM structure of the activated XauSPARDA filament. **(A)** Representative micrograph showing XauSPARDA filaments. **(B)** Representative 2D classes of filament assembly. **(C)** Cryo-EM map of the filament fragment containing DREN-domain assembly with the bound dsDNA. Part of the map covered by the model is colored. The filament is formed by dimeric “steps”: the central step (yellow) carries the DREN domains tetrameric assembly, which binds the DNA substrate (dark grey). The neighboring steps (green) donate the DREN domains into the active center DREN tetramer. The distant steps, represented in the model by only one monomer (magenta), donate DREN domains stabilizing active center assembly. gRNA colored red, tDNA - blue. **(D)** Schematic of XauSPARDA activated complex assembly. Binary monomers are colored as in C. **(E)** Dimer of two binary XauSPARDA-gRNA/tDNA monomers within the “step”. No DREN domains are shown. **(F)** The structure of the XauSPARDA binary monomer with bound gRNA/tDNA (without DREN domain) is colored as shown in Figure 1A. See Figures S5, S6, and Table S4.

The steps within the filament are connected by two interstep surfaces formed by the Ago proteins: a large symmetric interface (contact area ∼735 Å^2^) and a small asymmetric interface (contact area ∼420 Å^2^) (Figure 3D, Table S4). A tetramer of DREN domains bound to the dsDNA substrate is also involved in filament formation (its detailed structure will be presented below).

### XauSPARDA recognizes the 5′-AU dinucleotide of the RNA guide

In the activated XauSPARDA complex, the gRNA/tDNA heteroduplex is bound in the positively charged groove formed between the APAZ domain and the XauAgo MID and PIWI domains, similarly as in the case of the activated SPAR(S/T)A complexes (Figures 3E, 3F, and S6B). The gRNA and the tDNA strands make multiple salt-bridge and hydrogen-bonding interactions with the APAZ:Ago part (Figure 4A).

**Figure 4.**
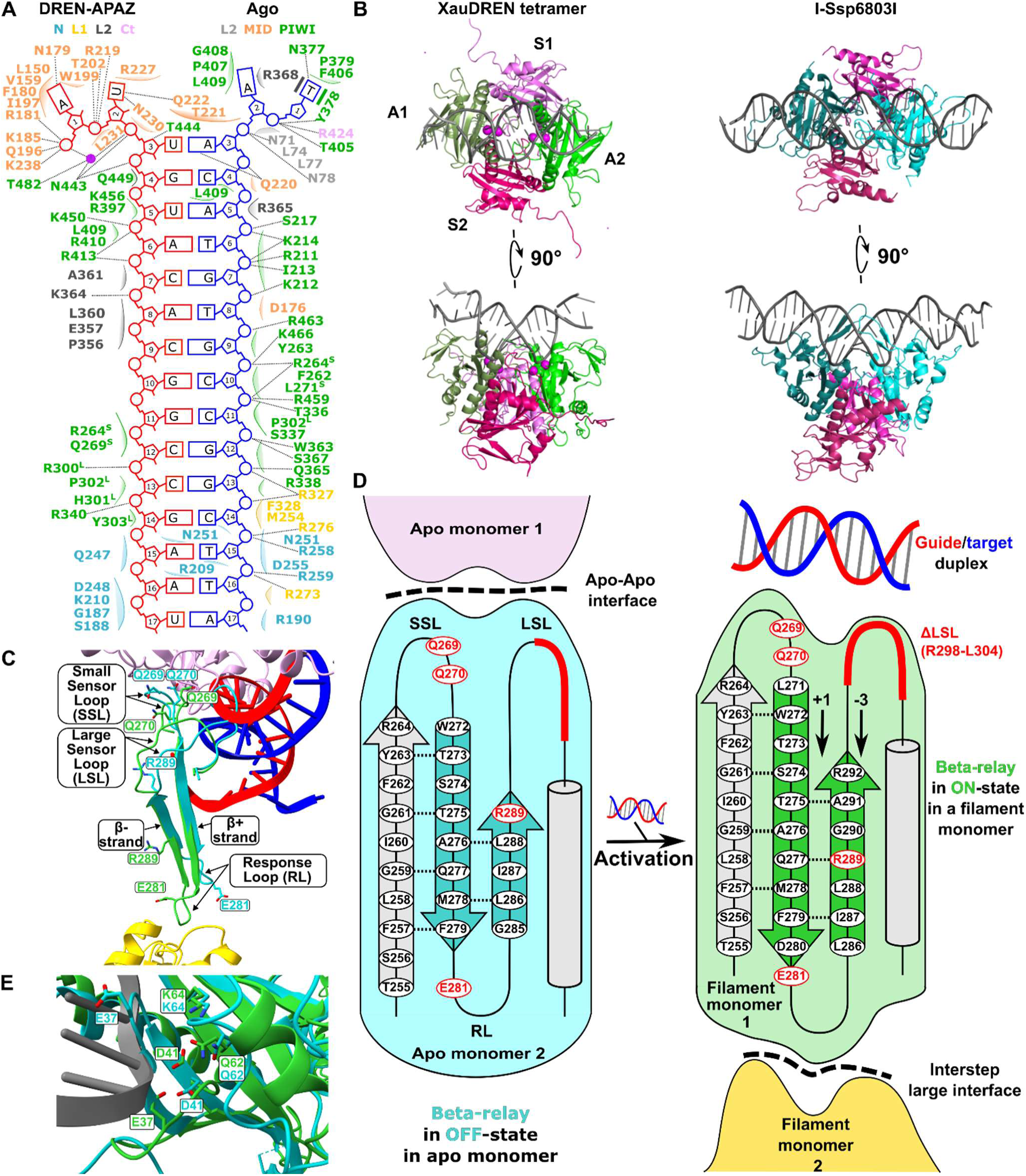
XauSPARDA uses the beta-relay mechanism for activation. **(A)** Schematic of the XauSPARDA binary monomer interactions with the gRNA (red) and tDNA (blue) strands. Interacting residues are colored according to the domain scheme. H-bonds or salt bridges are shown as dotted lines. Ca^2+^ is shown as a magenta sphere. Concave surfaces mark vdW interactions. Stacking is marked by bars. S and L superscripts indicate residues from small and large sensor loops, respectively. **(B)** DREN domain tetramer of XauSPARDA corresponds to homing endonuclease I-Ssp6803I. A1/2 and S1/2 indicate the active DREN (binds dsDNA) and the structural DREN domains, respectively. Ca^2+^ is shown as magenta (XauSPARDA) and cyan (I-Ssp6803I) spheres. **(C)** Beta-relay in the context of XauSPARDA interchain interactions. Components of the beta-relay are indicated. Apo (turquoise) and gRNA/tDNA duplex-bound (green) monomers of XauSPARDA are overlapped. Apo monomers (turquoise and pink) interact within the apo dimer; gRNA/tDNA-bound monomers (green and yellow) interact within the filament via the large interstep interface. **(D)** Schematic depiction of the beta-relay action upon XauSPARDA activation. The mutated residues (including LSL deletion) are colored red. Dotted lines indicate hydrogen bonds between β-strands. Arrows with values show the register shifts of the beta-relay upon gRNA/tDNA duplex binding. **(E)** Conformational changes of the XauSPARDA nuclease active center upon transition from the apo form (turquoise) to the active DREN conformation (green). DNA substrate is colored grey. For clarity only a single XauSPARDA monomer of each form is shown. See also Figures S1, S6, S7 and Table S4.

The first two base pairs of the bound heteroduplex are disrupted in the activated XauSPARDA complex as in the activated SPAR(S/T)A complexes (Figure S7A).^24–31^ The 5′-terminal phosphate group of the guide RNA is fixed in a highly conserved MID domain pocket and coordinated by a Ca^2+^ ion (used instead of Mg^2+^ to preclude DNA cleavage in cryo-EM experiments) as observed for other pAgo-gRNA complexes (Figure S7B).^23,43–45^ The first guide A base (g1A) is flipped out of the bound heteroduplex into the MID domain pocket, where beside several nonspecific vdW interactions, the side chain of the N179 residue makes a base-specific contact with the N6 atom of the adenine (Figures 4A and S7B). The second guide U base (g2U) makes base-specific contacts with the Q222 and N230 residues (Figures 4A and S7C). No base-specific contacts are made with further RNA guide bases (g3-17) (Figure 4A). These structural data agree with our *in vivo* and *in vitro* results that the 5′-AU dinucleotide of the cellular RNA guides bound by XauSPARDA is critical for XauSPARDA activity (Figures 1K and 2F). The first two unpaired bases of the target are bound mainly by the MID and PIWI domains (Figures 4A and S7A). The target DNA strand makes mostly sequence non-specific contacts (Figure 4A). Although there is some structural specificity for the first flipped T base of the target (t1’T), our *in vitro* data show that its replacement with another base had no significant effect on XauSPARDA activity (Figure 2G).

### dsDNA substrate is bound by the DREN domain tetramer characteristic of homing endonucleases

The dsDNA substrate is bound by a tetramer of the nucleolytic DREN domains (the total tetramerization surface is ∼ 4900 Å^2^), where two DREN domains bind the substrate DNA (named **A**ctive A-DREN domains coming from the ±1 steps), while the other two support them (**S**tructural S-DREN domains coming from the ±2 steps) (Figure 3D). The dsDNA substrate is bound nonspecifically (overall contact area is ∼1400 Å^2^, Table S4), which is in line with our results demonstrating a nonspecific DNA cleavage by XauSPARDA *in vivo* and *in vitro* (Figures 1G, 1H, and 2A-2C). Surprisingly, the tetrameric assembly of DREN domains is very similar to the tetrameric arrangement of homing endonuclease I-Ssp6803I, which also belongs to the PD-(D/E)XK nuclease family. The active sites in both are located at the minor groove side of the bound DNA backbone and coordinate one and two Ca^2+^ ions in I-Ssp6803I and XauSPARDA, respectively (Figures 4B and S7E).^46^

### Significant conformational changes in apo monomer occur upon XauSPARDA activation

To find out in more detail the structural mechanism of XauSPARDA activation when drastic changes occur even in the oligomeric state of the protein (from apo dimer to the activated polymer), we analyzed the conformational changes of the protein during the transition from the apo to its filamentous form.

First, the negatively charged C-terminal tail (Ct, residues 414-444, Figure S1A) of the DREN-APAZ protein is sterically displaced from the positively charged NA binding groove by the target strand of the bound guide/target heteroduplex and becomes invisible (Figure S7F). In the case of SPARTA systems, it was shown that Ct performs an autoinhibitory function during the target binding, thereby preventing uncontrolled activation of these systems.^23^

Second, the disordered region (199-223 residues) in the apo form becomes ordered and interacts with the backbone of the DNA target strand (t3’-7’) (Figure S7G). The main chain of the 220-222 residues and the side chain of Q222 disrupt the first two base pairs of the gRNA/tDNA heteroduplex. The flipped t1’-t2’ bases are bound within the pocket formed mainly by the XauAgo protein (Figure S7G).

Third, the gRNA/tDNA duplex binding triggers a cascade of large-scale rearrangements in a structural subregion of the PIWI domain we named beta-relay. It is composed of: i) two sensor loops (the large sensor loop (LSL), which was previously identified as the sensor loop in structurally homologous SPAR(S/T)A,^23,32^ and the small sensor loop (SSL)) connected to ii) two β-strands (named β+ and β-denoting the directions of their movement relative to the rest of the β-sheet during XauSPARDA activation) ending by iii) a response loop (RL) (Figures 4A, 4C and 4D). In apo XauSPARDA, the beta-relay is in the inactive “OFF” state, where its SSL participates in apo dimerization *via* Q269 and Q270 residues (Figures 4C and 4D). The beta-relay is activated and switched into its “ON” state when the bound complementary gRNA/tDNA duplex sterically replaces the second apo monomer and its 10-13 base pairs push both SLs, causing a displacement of the β+ strand into the direction of its C-terminus by one residue (+1), while the β-strand gets pushed toward its N-terminus by three residues (−3) (Figure 4D). Since these two β-strands are antiparallel, they move in the same spatial direction, extending RL into an appropriate conformation to interact with another activated XauSPARDA monomer within the filament through the large interstep interface (*via* E281 and R282 residues) (Figure 4D). This structural model is in good agreement with our *in vitro* and *in vivo* results where the mismatches at the 10-13 bp positions of the gRNA/tDNA duplex, either a shortening of the RNA guide or the DNA target strand to 13 nt and LSL deletion (XauΔLSL mutant, where 298-304 residues are deleted) completely abolished the activity of XauSPARDA (Figures 2E, 2G, 4A and 5A-5C). Thus, the beta-relay is responsible for “sensing” the binding of the proper complementary gRNA/tDNA duplex and transferring this recognition signal into the XauSPARDA filament interface.

Fourth, the DREN domain undergoes a transition from the “closed” orientation, where it contacts the remaining part of the monomer (including Ct) *via* multiple interactions, to the “open” orientation, where its interactions with the remaining part of the monomer are lost (Figure S7H). One may assume the bound gRNA/tDNA duplex displaces Ct that holds the DREN domain in the “closed” orientation (interactions R424:E31 and S241:R33), releasing the DREN domain into the “open” orientation required for binding and cleavage of the dsDNA substrate (Figure S7I).

Fifth, the active site of the DREN domain undergoes a structural reorganization, shifting from a conformation that sterically hinders dsDNA substrate binding to one that is catalytically competent (Figure 4E). The β-hairpin bearing the catalytic E37 residue undergoes the most prominent structural changes: the β-strands shorten, elongating the E37 loop that adopts a conformation suitable for dsDNA binding and cleavage (Figure 4E).

All of the cascade conformational changes of XauSPARDA presented above explain the transition from an inactive apo to a catalytically competent filamentous form safeguarding against its premature activation.

### Filament formation is essential for XauSPARDA activity

To determine the importance of filament formation for the enzymatic activity of XauSPARDA, we performed mutagenesis of the interfaces formed between monomers in the filament. Multiple mutations were introduced into the XauAgo protein (**FA** mutants from **f**ilamentation by **A**go) to disrupt the intrastep (FA1 mutant), large asymmetric (FA2 mutant), and small symmetric (FA3 mutant) interstep interfaces and into the XauDREN-APAZ protein (**FD** mutants from **f**ilamentation by **D**REN) to disrupt different interfaces of the DREN tetramer (FD1, FD2; FD3) (Figure 3D, Table S4). The expression of all these mutants in *E. coli* cells was similar to that of WT XauSPARDA (Figure S3H); the mutant proteins were also successfully purified (Figure S3G). Unlike WT XauSPARDA, all mutants were non-toxic *in vivo* (Figures 5A-5C) and were (almost) inactive DNases *in vitro* (Figure 5D). Thus, the intact interfaces between the activated monomers within the filament are essential for the functional activity of XauSPARDA.

**Figure 5.**
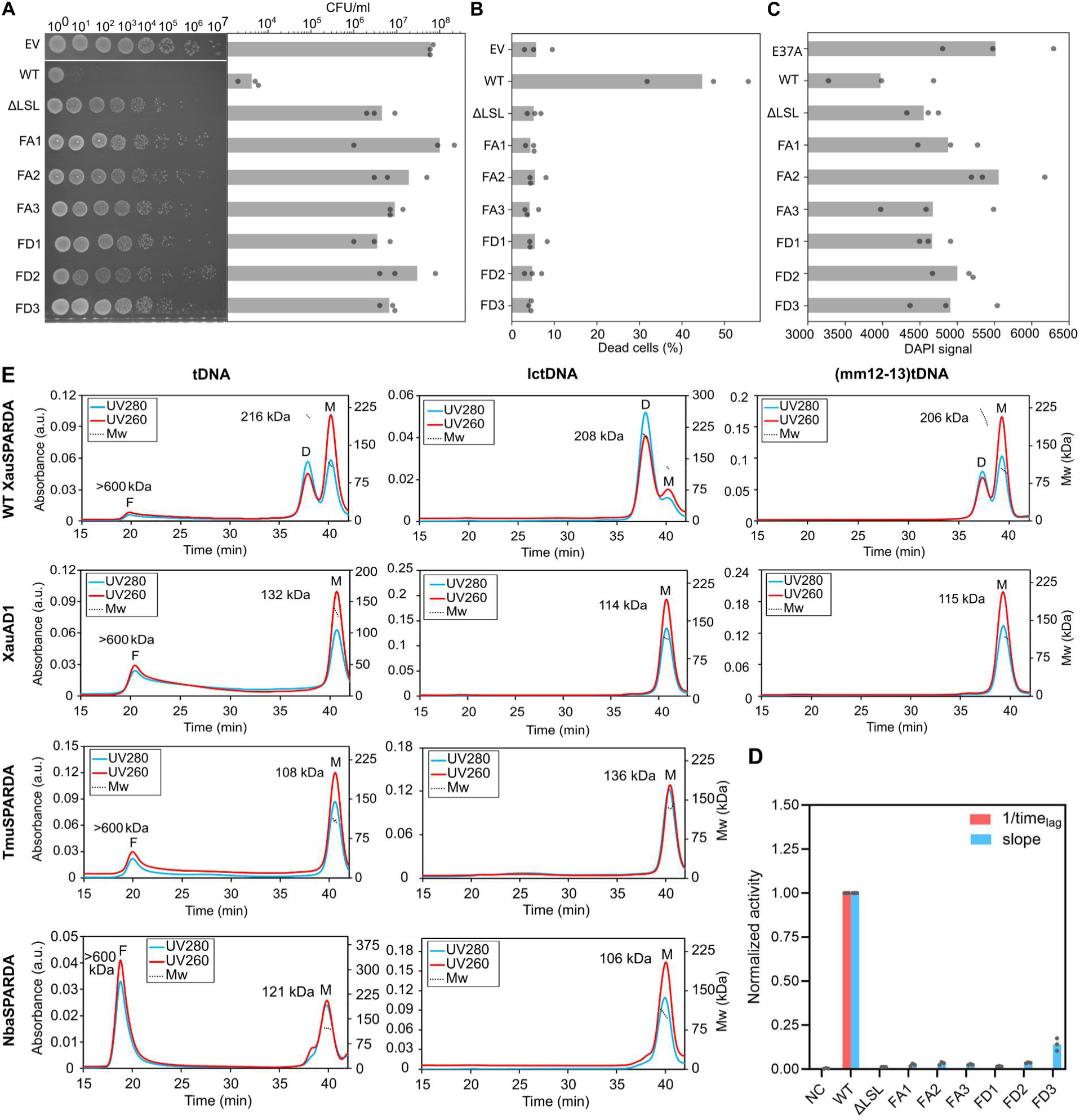
Filament formation is essential for XauSPARDA activity. **(A)** Cytotoxic effect of XauSPARDA system on *E. coli* cells expressing WT and mutants. EV - empty vector. Left: serial dilutions on agar plates, right: colony forming units (CFU) were counted, and the average of three biological replicates is shown in bars. **(B)** Viability of *E. coli* cells expressing SPARDA systems in the presence of pCDF plasmid. Dead cells were counted by flow cytometry using propidium iodide (PI) staining. The average of three biological replicates is shown in bars. **(C)** DAPI signal of *E. coli* cells expressing SPARDA systems in the presence of pCDF plasmid. An integrated DAPI signal of stained cells was measured by flow cytometry. The average of three biological replicates is shown in bars. **(D)** Cleavage activity of XauSPARDA mutants. NC (negative control) - reaction without protein. **(E)** Filamentation analysis of XauWT, AD1, TmuSPARDA, and NbaSPARDA in the presence of fully complementary tDNA, low-complementarity (8 bp) tDNA or tDNA mismatch in 12-13 position, as determined by SEC-MALS. The absolute molar weight of proteins is indicated. For each sample, OD_280_ and OD_260_ absorbance profiles are shown. Molecular weight (MW) is shown as a dotted line. In each panel, positions of monomeric (M), dimeric (D), and filamentous (F) SPARDA peaks are indicated. See Figures S3, S7, and Table S5.

Next, we investigated the oligomeric state of XauSPARDA in the presence of the RNA guide, the (low) complementary DNA target, and the dsDNA substrate or their combinations using SEC-MALS (Figure 5E, Table S5). The formation of the nucleic acid-bound monomers of WT XauSPARDA and its catalytically inactive DREN domain mutant XauD41A was observed, indicating the apo dimer dissociation upon NA binding (Figure 5E, Table S5). Only in the presence of the RNA guide and the complementary DNA target (with or without added dsDNA), WT XauSPARDA and the XauD41A mutant form large nucleic acid-bound oligomers reaching >600 kDa, which elute with the free volume of the gel filtration column (Figure 5E, Table S5). These specific XauSPARDA oligomers (formed only in the presence of gRNA and tDNA) may correspond to the activated filament forms of various lengths. Interestingly, the XauAD1/2 mutants, which are already present as monomers due to the mutations disrupting the apo dimerization interface, form the specific oligomers much more efficiently than the WT XauSPARDA (Figure 5E, Table S5). Neither WT XauSPARDA nor its XauAD1 mutant formed the specific oligomers in the presence of the DNA target with the mismatches at the 10-11 and 12-13 bp positions. SSL and LSL of the beta-relay contact these positions, which are critical for the XauSPARDA activation (Figures 2G, 4A, and 5E, Table S5). Despite the XauΔLSL mutant (the LSL of the beta-relay is deleted) dissociating into monomers upon binding the guide and target, it fails to form the specific oligomers (Table S5). Disruption of the large asymmetric filament interface formed by the XauAgo protein (XauFA2 mutant) did not result in the specific oligomerization under all tested conditions (Table S5). These results support our previous structure-based hypothesis that, upon binding the complementary DNA target, the XauSPARDA apo dimer dissociates into monomers, then the activated monomers assemble into a filament, and the beta-relay plays a critical role in this process (Figure 4D). It should be noted that the isolated DREN domain is a monomer in solution and does not exhibit DNase activity (Figure 2D, Table S5). Thus, it appears that the filament assembly is the key to DREN nuclease activation as it brings DREN domains into close proximity with each other, enabling them to form a catalytically active tetramer.

### Real-time visualization of filamentation at a single-molecule level

Although in the cryo-EM structure of the activated XauSPARDA filament, the DREN tetramers nonspecifically bind short (42 bp) dsDNA substrates *in trans* (Figure 3C), *in vivo* activated SPARDA interacts with and cuts longer dsDNA substrates (> 1 kb) such as plasmids or genomic DNA (Figure 1G). Therefore, during the formation of the activated SPARDA filament *in vivo*, the DREN tetramers theoretically should bind many sites of long dsDNA molecule, thus compacting and cleaving it. We performed *in vitro* single-molecule fluorescence microscopy to observe the real-time interaction of the activated XauSPARDA with single end-immobilized phage λ DNA molecules in solution. WT XauSPARDA-gRNA/tDNA complex was injected into such flow cell in the presence of Ca^2+^ ions to prevent hydrolysis of the dsDNA substrate (Figures 6A–6B). The kymograph (Figure 6C) and movie (Video S1) illustrate how surface-immobilized continuous slow buffer flow-stretched DNA molecules begin to shorten upon injection of the WT XauSPARDA-gRNA/tDNA complex until DNA is fully compacted into a bundle (Figure 6B). These experiments demonstrated that ATTO647N-labeled tDNA puncta colocalize with the compacted DNA bundles, indicating the formation of a compacted λ DNA-XauSPARDA-gRNA/tDNA complex (Figure 6E and Video S2). The activated WT XauSPARDA compacts dsDNA reversibly: the XauSPARDA complex dissociates from DNA upon prolonged exposure to higher ionic strength buffer, and free DNA can be re-compacted by injection of the XauSPARDA complex again (Figure 6B). In addition, the exchange of Ca^2+^ ions into Mg^2+^ ions in the flow cell, where λ DNA is compacted by XauSPARDA-gRNA/tDNA, leads to nearly complete DNA cleavage – only some short DNA fragments in the compacted DNA bundles remain (Figure 6B). Such compaction of λ DNA molecules into bundles and its subsequent degradation was not observed in the presence of the low complementarity DNA target (lctDNA), which also did not support the specific oligomer formation in the SEC-MALS experiments and the nuclease activity *in vitro* (Figures 5E and S7J). The XauD41A (the active site mutant) specifically compacted the immobilized λ DNA but did not cleave it (Figures 6F and 6G). The apo dimerization mutants AD1 and AD2, like the WT XauSPARDA, were able to compact the immobilized λ DNA and then degrade it (Figures 6F and 6G). XauSPARDA mutants FA1, FA2, FA3, FD1, and FD2, in which the filament-forming interfaces were disrupted, did not compact the immobilized λ DNA (Figure 6F) nor showed DNA cleavage activity *in vitro* (Figure 5D) even in the presence of the complementary DNA target.

**Figure 6.**
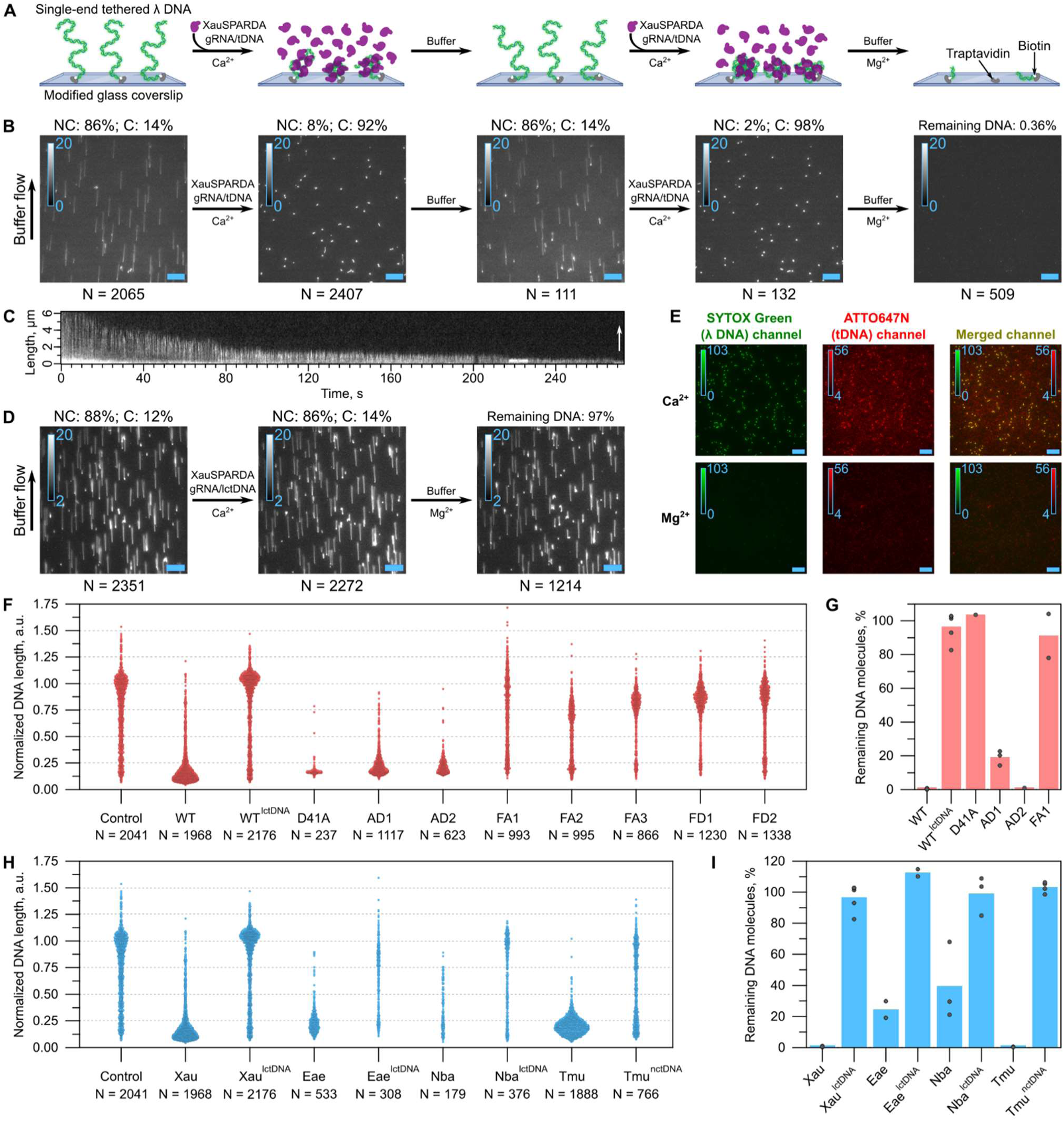
Direct real-time single-molecule total internal reflection fluorescence microscopy (TIRFM) *in vitro* reveals SPARDA acting as a dsDNA compacting endonuclease. **(A)** Schematics illustrating a DNA flow-stretch assay employed to investigate SPARDA and dsDNA interaction dynamics at the single-molecule level. Arrows indicate the essential components injected into the flow cell’s channel during each step. **(B)** The reversible compaction of single-end immobilized lambda bacteriophage DNA (λ DNA) by the WT XauSPARDA-gRNA/tDNA complex (in the presence of Ca^2+^) and its cleavage (in the presence of Mg^2+^) as depicted in (A). All TIRFM images were acquired with the enabled buffer flow (indicated by a vertical arrow). N - the total number of λ DNA molecules analyzed, NC - the mean percent value of non-compacted DNA, and C - compacted DNA. **(C)** A representative kymograph demonstrates a gradual shortening of a single λ DNA molecule by the activated WT XauSPARDA in the presence of Ca^2+^ ions and buffer flow (white arrow) during the entire time-lapse series acquisition. **(D)** Response of λ DNA molecules to the WT XauSPARDA-gRNA/lctDNA complex in the presence of Ca^2+^ and Mg^2+^ (where lctDNA - low complementarity target DNA). **(E)** Co-localization of λ DNA (SYTOX, green) and the WT XauSPARDA-gRNA/tDNA complex containing ATTO647N-labeled tDNA (red). λ DNA was first compacted (in the presence of Ca^2+^), then cleaved (in the presence of Mg^2+^). TIRFM images shown here were acquired without the buffer flow. **(F)** Distribution of measured λ DNA lengths after incubation (group ‘Control’ - before incubation) with the complexes of XauSPARDA proteins and the complementary gRNA/tDNA duplex (or gRNA/lctDNA, where indicated) in the presence of Ca^2+^. **(G)** Cleavage activity of XauSPARDA variants in the presence of Mg^2+^. Bars represent an average percentage of the remaining non-cleaved λ DNA molecules, whereas dots indicate individual values determined for every separate surface position imaged. In each case, the collected N>500. **(H)** Complexes of homologous SPARDAs with gRNA/tDNA compact λ DNA in the presence of Ca^2+^. The λ DNA lengths are displayed in a scatter-plot representation. In control experiments, gRNA/lctDNA or gRNA/nctDNA were used (where nctDNA – non-complementary target DNA). **(I)** Activated homologous SPARDAs cleave immobilized λ DNA molecules in the presence of Mg^2+^. The collected N>300 was in each case except for NbaSPARDA-gRNA/tDNA (N = 68). Here, data for each sample was taken from at least three different surface positions. The scale bar (light blue) in all images is 10 µm. The calibration bar with indicated minimum and maximum gray or color scale values is provided at the upper left of each image.

### Filamentation is also characteristic of homologous SPARDAs

To gain insight into the prevalence of filamentation in other SPARDA systems, we modeled them as dimers using AlphaFold-Multimer. Our initial modeling attempts revealed that AlphaFold often prioritizes dimer interface formation through DREN domains at the expense of Ago-Ago interactions. To alleviate this issue, we resorted to modeling the truncated DREN-APAZ:Ago dimer models (TDMs) by removing DREN domains. Additionally, by splitting off the nuclease Mrr domain, we similarly modeled the SPARMA system from *Kordia jejudonensis* (Kje system) belonging to the branch of short pAgos, different from SPARDA (Figure 1B).^14,38^ The resulting TDMs were grouped by their structural similarity, resulting in three large groups of models: two groups correspond to XauSPARDA filament intrastep and large interstep interfaces, respectively, while the third one closely resembles Xau and EaeSPARDA dimeric apo form (Figure S7K and S7L). Only Xau and EaeSPARDA TDMs were present in the third group, matching the Xau and EaeSPARDA dimeric apo form obtained from X-ray crystallography. This is in line with our and others’ published experimental results showing that unlike dimeric apo Xau and EaeSPARDA, the apo forms of the Kje system, Tmu and NbaSPARDA exist in solution as monomers (Table S3).^14,35,38^ However, most of the remaining TDMs resembled either of the two filament interfaces, suggesting that all the selected systems may assemble into filaments.

To test whether the filament formation is also characteristic of other previously characterized SPARDA, as suggested by the AlphaFold modeling experiments, we purified NbaSPARDA and TmuSPARDA proteins.^14,35^ According to SEC-MALS data, unlike apo dimeric XauSPARDA, both apo Nba and TmuSPARDA proteins are monomers in solution as determined previously (Tables S3 and S5). However, like XauSPARDA, both proteins form specific oligomers in the presence of the RNA guide and complementary DNA target (and dsDNA) (Figure 5E, Table S5). Moreover, in the single-molecule experiments, all of them (including EaeSPARDA) efficiently compacted immobilized λ DNA and cleaved it only in the presence of a complementary DNA target, with which their nuclease activity was observed in bulk experiments (Figures 6H, 6I and S7J). Thus, filament formation during activation may be a common feature of SPARDA enzymes.

### Beta-relay sensor mechanism is widespread among pAgos

Discovery of the beta-relay sensor mechanism in XauSPARDA prompted us to explore whether this feature is unique to XauSPARDA or is also present in other pAgos. To this end, we collected PDB structures of pAgos solved in both the apo (inactive) and guide/target bound (active) forms. We then compared the structures of these two forms and looked at whether the register of β-strands corresponding to those involved in the XauSPARDA beta-relay differs between these two forms. We found that the beta-relay is present not only in other clades of short pAgos but also in representatives of long-B and long-A pAgos (Figure 7A). Thus, we identified the beta-relay in three different SPARTA systems^24–31^ and GsuSPARSA^32–34^ that are all members of short pAgos We also identified the presence of the beta-relay in fAfAgo^21^ (split long-B pAgo) and in very structurally similar *Sulfolobus islandicus* SiAgo system.^47^ Among pAgos from the long-A clade, we identified the beta-relay in PfAgo,^48,49^ TtAgo,^6,50^ and KmAgo^51^ but not MpAgo.^52,53^ Since our structural analysis linked the beta-relay activation to the guide/target binding, we were curious whether distortions within the guide/target duplex, such as mismatches and bulges, may affect this switch. We analyzed TtAgo structures solved with guide/target duplexes containing bulges in various positions. It turned out that the beta-relay was activated in all these structures except the one with the bulge formed between the 9’ and 10’ positions of the target strand (PDB ID: 5XPA). In this case, the guide/target distortion enabled accommodation of the small sensor loop without the shift of the β-strands.

**Figure 7.**
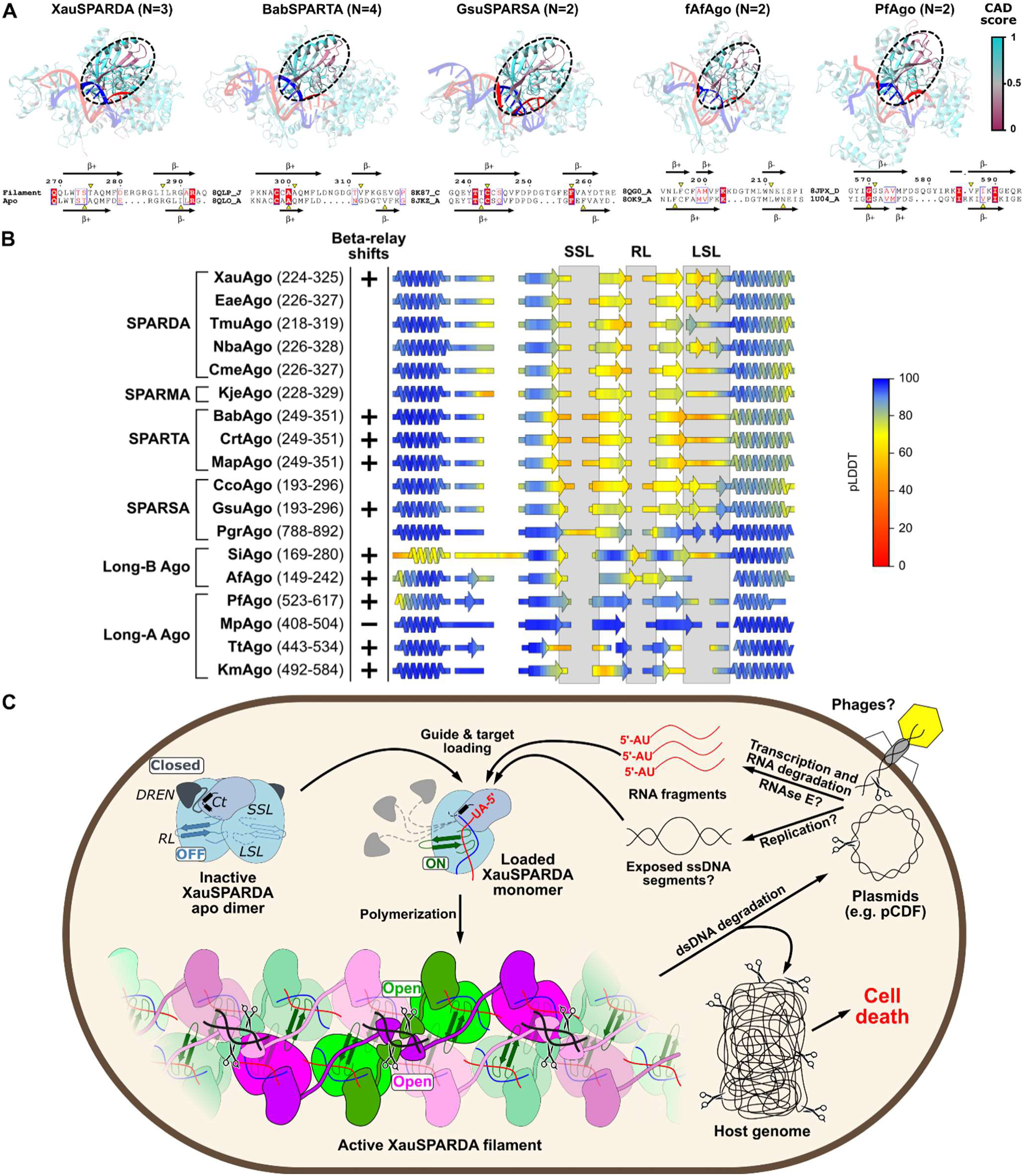
Beta-relay sensor mechanism is widespread among pAgos. **(A)** Main conformational changes in a diverse set of pAgo systems upon activation, as indicated by CAD-score. Low CAD-score values (colored in maroon) indicate significant local conformation changes in compared structures. Dashed lines outline beta-relay locations in each structure. The beta-relay location in each structure is highlighted. Guide nucleic acids are colored red, while target DNA molecules are colored blue. N values denote the magnitudes of the outer β-strand shifts upon beta-relay activation. The pairwise alignments depict the spatial correspondence of the residues in the outer β-strand and the neighboring residues in both the “ON” and “OFF” states. The yellow triangles highlight the positions of arbitrarily chosen residues in both beta-relay states. **(B)** Summary of available experimental data and top AlphaFold model data on the beta-relay region in various pAgos. Colored protein secondary structure elements are derived from the structural alignment of top Alphafold models for minimal units (monomers/heterodimers) of pAgo systems. “+” and “-” symbols indicate whether the beta-relay shifts are observed when comparing experimental active and inactive protein state structures. The absence of symbols indicates that the pair of experimental protein structures is unavailable. **(C)** Mechanism of action of XauSPARDA. See also Figure S7.

We found that the extent of β-strand register shifts in different pAgo systems may differ. Interestingly, in all beta-relays, the inner β-strand (β+) consistently shifts forward by a single position, while the outer β-strand (β-) shifts backward by N positions, where N ranges from 0 to 4 and is system-dependent (Figure 7A). Notably, in many structures, the outer β-strand displays significant conformational heterogeneity, and in some cases, the extent of its register shift is ambiguous. Despite some ambiguity regarding the outer β-strand, the outcome of beta-relay activation is unambiguous. Since the two β-strands of the beta-relay are antiparallel, the residues of both strands are moving in the same direction relative to the rest of the pAgo structure. The consequence of these β-strand register shifts is that part of the β-sheet surface is remodeled, LSL and SSL become shorter while RL is extended (Figure 7A).

AlphaFold typically assigns lower confidence values to dynamic regions of proteins.^54^ Since the beta-relay involves significant rearrangement of the local structural environment, AlphaFold might be expected to assign lower confidence values to the corresponding region. To test this idea, we remodeled monomeric structures for pAgos used in structural analysis. In addition, we modeled several additional short pAgos that have been studied experimentally but did not have PDB structures. All resulting models had high global confidence scores. However, regions corresponding to beta-relays in short pAgo models consistently had noticeably lower local confidence scores (pLDDT) than the neighboring regions. In the long pAgo models, such a drop in pLDDT scores was either less pronounced or not present (Figure 7B). These findings suggest that the low local confidence in AlphaFold models might be a strong indicator of the beta-relay presence, whereas the high local confidence in this regard is less symptomatic.

## DISCUSSION

Here, we present detailed functional, biochemical, single-molecule, and structural studies of the yet uncharacterized SPARDA systems consisting of short pAgo proteins and the associated DREN-APAZ effectors, allowing us to propose a detailed mechanism of their action. Our studied Xau and EaeSPARDA are anti-plasmid protection systems, like the previously characterized Nba and CmeSPARDA, which were also able to protect *E. coli* cells from phages.^14^ It is not excluded that, given the proper experimental conditions, our SPARDAs would also exhibit anti-phage activity. Activated XauSPARDA nonspecifically degrades both the trigger plasmid and *E. coli* genomic DNA by introducing DSBs, which causes the host’s death, thus preventing the further spread of the mobile element in the population. Such a strategy of host suicide-based protection against foreign genetic information is common among prokaryotic defense systems.^55,56^

Based on the structures in apo (X-ray) and the activated filament (cryo-EM) forms, along with *in vivo* and *in vitro* biochemical and single-molecule experiments, we propose the following molecular mechanism of XauSPARDA function (Figure 7C, Video S3). Initially, the apo binary dimer (DREN-APAZ:Ago) exists in an inactive autoinhibited state. In this state: i) the DREN domains with the catalytically incompetent active sites are positioned on the opposite sides of the complex and are locked in a “closed” conformation by interactions with other regions, including the negatively charged Ct-tail of DREN-APAZ; ii) the positively charged groove for binding of the gRNA/tDNA duplex is occupied by the C-tail; iii) the beta-relay, responsible for the “sensing” of the complementary gRNA/tDNA duplex is in the “OFF” state, as the SSL and LSL loops of the beta-relay participate in the dimerization interface. The activation begins when XauSPARDA binds the 5′-AU-RNA guide derived from invader transcripts (e.g., RNase E degradation products of pCDF plasmid CloDF13 *ori* region transcripts) and the target DNA, complementary to the guide. The binding of the RNA guide and the DNA target likely occurs sequentially since the APAZ:pAgos module loaded with the guide should make the DNA target search more efficient. While XauSPARDA can also bind a preformed gRNA/tDNA duplex, this process is less efficient (Figure S3P). XauSPARDA can only be activated by the single-stranded DNA target as it cannot unwind a stable DNA duplex (Figure S3O). A sufficient length of ssDNA might be formed during active replication of the invader’s DNA as the transcription bubble (9 bp) might be too small and inaccessible.^57^ An illustrative example is the replication of the pCDF plasmid, which triggers XauSPARDA activity. It begins with forming a long R-loop, which could serve as a ssDNA target,^58^ whereas the transcripts of the pCDF *ori* region preferentially bound by XauSPARDA may serve as a source of RNA guides. Upon binding a gRNA/tDNA duplex, Ct is removed from the NA binding groove due to steric clashes. In turn, the Ct interaction with the DREN domain is disrupted, and the DREN domain detaches from the APAZ:pAgo module. Ct may also facilitate the efficient exchange of RNA guides until a stable interaction with the complementary DNA target is established. The clashes between the SSL and LSL loops and the bound gRNA/tDNA duplex (at positions 10-13) switch beta-relay “ON” by initiating shifts of the β+ and β-strands in the PIWI domain. As a result, the RL lengthening, accompanied by extensive surface remodeling, produces new interaction surfaces necessary for filament formation. Since the apo dimer is structurally incompatible with both gRNA/tDNA binding and filament formation, the dimer has to dissociate into NA-bound monomers, as confirmed by our SEC-MALS experiments (Figures 5E and S6D). The gRNA/tDNA-bound monomers then polymerize *via* intra- and interstep Ago interfaces, producing a functionally active filament. Once the filament is formed, the DREN domains, detached from APAZ/Ago, transition into the “open” conformation and tetramerize due to their spatial proximity. Within the tetramer, the two A-DREN domains can bind and cleave the dsDNA substrate, while the other two S-DREN domains provide structural support (Figure 4B). Presumably, the formation of an activated filament is favored when rapid transcription and replication of the invader produces many uniform RNA guides complementary to the relatively small plasmid or phage genomes as opposed to the much larger host genome. High local concentration of the gRNA/tDNA-bound XauSPARDA complexes also likely promotes filament assembly. Since the activated XauSPARDA efficiently cleaves any dsDNA, the filament initiated by the invading DNA will hydrolyze both the invader and the host genome, causing cell death and thereby preventing further spread of the invader in the cell population.

We were surprised that the beta-relay mechanism uncovered in our study is not confined to SPARDA systems. We identified beta-relays not only in the evolutionarily closest SPARTA and SPARSA systems, representing short pAgos, but also in pAgos of long-B and long-A clades. The activated SPARTA and SPARSA are tetramers, in which Ago subunits form dimers.^23–32^ In line with the close evolutionary relationship of Ago components, the dimer interfaces in SPARTA and SPARSA are reminiscent of the intrastep interface in XauSPARDA, but the orientation between Ago subunits differs slightly. Despite these differences, the beta-relay in both SPARTA and SPARSA is involved in higher-order assembly. For example, RL (loop 8-9) of the CrtSPARTA masks a positively charged surface pocket when the system is inactive but is displaced when a CrtSPARTA monomer binds the gRNA/tDNA duplex thereby facilitating dimerization.^28^ In the GsuSPARSA tetramer, RL (e.g., Thr 253) also participates in the Ago-Ago dimerization interface.^34^ The function of the beta-relay in long-B pAgo systems is not entirely clear. However, as most long-B pAgos are associated with distinct effector proteins,^20^ it is tempting to speculate that the beta-relay may be used to modulate interactions with these effectors and thereby regulate their activities. The observation that the SiAgo system stably interacts with its transmembrane effector, Aga2, only when loaded with the guide/target duplex^47^ suggests a possible involvement of the beta-relay, but this has yet to be demonstrated experimentally. In long-A pAgos, the beta-relay appears to play a key role in the conformational changes of the sensor loops. In PfAgo and TtAgo, before target binding, the loop (LSL in our notation) hosting the catalytic glutamate (glutamate finger) is in the ‘unplugged’ conformation, far from the catalytic site (for review, see ^59^). In the apo form of KmAgo, both the glutamate finger and one of the catalytic aspartates are ‘unplugged’.^51^ Upon guide and target binding by these pAgos, SSL and LSL rearrange, allowing positioning of the glutamate finger into the ‘plugged-in’ conformation to complete the catalytic DED(D/H) tetrad.^49^ In PfAgo, the activation is also accompanied by dimerization involving residues from both SSL and LSL.^49^ Whether or not RL lengthening plays some role in the beta-relay signaling in long-A pAgos remains to be explored.

The apparent reason for the beta-relay activation is the engagement of the sensor loops (SSL and LSL) with the central region of the bound guide/target duplex. This is consistent with our results showing that only target strands longer than 13 and 15 nt activate Xau and EaeSPARDA, respectively (Figures 2E and S3N). Similar length requirements of the target strand were experimentally observed for short pAgos GsuSPARSA,^25^ and MapSPARTA^25^ and for long-A pAgos PfAgo,^49^ and TtAgo.^50^ The geometry of the guide/target duplex in the region facing the sensor loops is also important as some mismatches or bulges in this region also affect the beta-relay activation (Xau and EaeSPARDA (Figures 2G and S3R, respectively), GsuSPARSA,^25^ Crt^3^ and MapSPARTA.^3,25^ Additionally, the interplay between sensor loops and a guide/target duplex here and in other studies was shown by the lack of activity of pAgos with modified or deleted LSL even when a complementary target of an optimal length was used (XauSPARDA (Figure 5D), GsuSPARSA and MapSPARTA^25^). Thus, several types of biochemical data directly support the beta-relay mechanism discovered here using structure-based analysis.

We did not detect the beta-relay in eAgos, but this does not necessarily mean that eAgos do not use such a signaling mechanism. In many cases, there are no pairs of eAgo structures in both inactive and fully activated (bound to long complementary guide/target duplexes) states (reviewed in ^59^). Also, some inactive-active eAgo structure pairs have relatively low resolution, which might preclude the identification of residue shifts in β-strands that are part of the beta-relay substructure.

Recent discoveries of new prokaryotic defense systems and studies of their mechanisms of action have revealed the emerging trend of supramolecular assemblies (including filaments) in bacterial immunity^60–66^ (reviewed in ^67^). Moreover, it was observed that the organization of immune system components into supramolecular assemblies is not confined to bacteria. In eukaryotic immunity, the concepts of ‘signaling via cooperative assembly formation’ (SCAF) and ‘supramolecular organizing centers’ (SMOCs) describe processes whereby components of the immune systems self-assemble into scaffolds that activate downstream effectors, leading to the effective immune response.^67–69^ As in other prokaryotic defense systems, pAgo-based supramolecular assemblies facilitate the formation of the effectors’ composite active site (TIR domain in SPARTA^23–31^) or allosteric activation of the effector domains (Sir2 domain in SPARSA^32^). In the case of XauSPARDA and likely other SPARDAs, filament assembly facilitates the formation of a DREN tetramer, coupling this with allosteric changes that produce an active site competent for dsDNA cleavage. It is thought that the supramolecular assembly of immune system components evolved as a way to control the cytotoxicity of effectors, reducing the risk of autoimmunity from spurious activation.^67^ In the case of SPAR(D/S/T)As it is likely that a concentration of gRNA/tDNA duplex-bound pAgos exceeding the threshold for the formation of activated supramolecular assemblies is achieved only as a result of infection. Otherwise, the host cell harboring one or more of such defense systems would be in constant danger of their accidental activation, leading to cell death. Finally, an intriguing question remains whether eAgo, similarly to homologous pAgos, also utilizes the beta-relay mechanism to regulate their activity via homodimerization (as in long-A pAgos) or interaction with other proteins.

### Limitations of the study

In this study, the experiments with SPARDAs were performed *in vitro* using purified proteins or *in vivo* using heterologous expression systems in *E. coli* under artificial promoters. It is possible that SPARDAs could protect against bacteriophage infection in their native hosts. We also didn’t directly demonstrate filament formation *in cellulo*, as such a study poses significant technical challenges and requires a separate project.

## RESOURCE AVAILABILITY

### Lead contact

Further information and requests for resources and reagents should be directed to and will be fulfilled by the lead contact, Mindaugas Zaremba.

### Materials availability

Plasmids generated in this study are available upon request from the lead contact with a completed Materials Transfer Agreement.

### Data and code availability

- The cryo-EM 3D reconstruction and crystal structure have been deposited in the Electron Microscopy Data Bank and Protein Data Bank. The accession numbers of publicly available data analyzed in this paper are listed in the key resources table.
- This paper does not report the original code.
- Any additional information required to reanalyze the data reported in this paper is available from the lead contact upon request.

## ACKNOWLEDGEMENTS

We thank Prof. Virginijus Siksnys and all colleagues for their comments on earlier versions of this manuscript. We thank Audrone Ruksenaite for the mass spectrometry of SPARDA proteins and Jokubas Siauciunas for cytotoxicity experiments of *E. coli* expressing SPARDA with pCDF. The authors also acknowledge Gleb Bourenkov for his assistance in using the beamline. Access to the EMBL beamlines at PETRA III (DESY) has been supported by iNEXT, project number 653706, funded by the Horizon 2020 program of the European Commission. The cryo-EM data collection by E.M. was supported by funding source 01.2.2-CPVA-V-716-01-0001 ‘EMBL partnership institution’. This work was supported by the Research Council of Lithuania (LMTLT) [S-MIP-24-40 to E.Z.] and Vilnius University Research Promotion Fund [MSF-JM-20/2023 to E.Z.]. Single-molecule microscopy was supported by the Research Council of Lithuania (LMTLT) [S-MIP-24-58 to M.T.], Horizon Europe HORIZON-MSCA-2021-SE-01 project FLORIN [Grant agreement ID: 101086142 to M.T.]. RNA-seq and DNA-seq were supported by the Research Council of Lithuania (LMTLT) [S-MIP-23-131 to M.Z.]. CROFT-seq adaptation was supported by Vilnius University Research Promotion Fund [MSF-JM-05/2024 to P.T.].

## AUTHOR CONTRIBUTIONS

Conceptualization, Č.V., and M.Z.; methodology and validation, E.Z., and M.Z.; formal analysis, E.J., and E.Z.; investigation, E.J., E.M., E.Z., A.K., S.A., I.D., U.T., A.S., P.T., A.G., M.T., and Č.V.; writing – original draft, E.Z., Č.V. and M.Z.; writing – review and editing, E.Z., Č.V., and M.Z.; visualization, E.Z. and M.Z.; supervision, M.Z.; project administration, and funding acquisition, E.Z. and M.Z.

## DECLARATION OF GENERATIVE AI AND AI-ASSISTED TECHNOLOGIES

During the preparation of this work, the authors used Grammarly and Google Translate in order to improve the readability and language of the manuscript. After using this tool, the authors reviewed and edited the content as needed and take full responsibility for the content of the published article.

## SUPPLEMENTAL INFORMATION

**Table S1. Plasmids and oligonucleotides used in this study. Related to Methods.**

**Table S2. X-ray and cryo-EM data collection, refinement, and validation statistics, related to Figures 2, 3 S4, S5, and S6**

**Table S3. SEC-MALS and Mw. Related to Figures 2 and S4.**

**Table S4. Contacts in different surfaces in apo and filament form of XauSPARDA system. Related to Figures 2, 3, 4, 5 and S6.**

**Table S5. SEC-MALS graphs. Related to Figures 2, 4, and 5.**

**Video S1. Fluorescence movie showing the activated WT XauSPARDA-mediated compaction of individual single end-tethered flow-stretched λ DNA molecules in real-time. Related to Figure 6.**

**Video S2. Dual-color footage of SYTOX Green-labeled λ DNA (green) real-time compaction performed by the WT XauSPARDA-gRNA/tDNA-ATTO647N complex (red) under a continuous buffer flow. Related to Figure 6.**

**Video S3. Structural changes of XauSPARDA upon its activation. Related to Figure 7.**

**Data S1. RNA- and DNA-seq results and analysis. Related to Figures 1 and S3.**

## METHODS

### KEY RESOURCES TABLE

| REAGENT or RESOURCE | SOURCE | IDENTIFIER |
| --- | --- | --- |
| **Antibodies** |  |  |
| 6x-His Tag monoclonal antibody | ThermoFisher | MA1-21315 |
| Goat anti-Mouse IgG (H+L) Secondary Antibody | ThermoFisher | 31320 |
| StrepTag II Monoclonal Antibody | ThermoFisher | MA5-37747 |
| **Bacterial and virus strains** |  |  |
| DH5α *E. coli* | Invitrogen | 18265017 |
| *E. coli* MG1655 | Blattner, F. *et al.*<sup>70</sup> | NCBI:txid511145 |
| *E. coli* BL21-AI | Bhawsinghka, N. *et al.*<sup>71</sup> | CP047231.1 |
| *E. coli* DH10B | Grant, S.G.N. *et al.*<sup>72</sup> | NCBI:txid316385 |
| Bacteriophage T2 | Gift from L. Truncaite lab collection | LC348380 |
| Bacteriophage T4 | Gift from L. Truncaite lab collection | NC_000866 |
| Bacteriophage T6 | Gift from L. Truncaite lab collection | MH550421 |
| Bacteriophage T3 | Gift from L. Truncaite lab collection | KC960671.1 |
| Bacteriophage T5 | ATCC collection, #11303-B5 | AY587007.1 |
| Bacteriophage T7 | Gift from L. Truncaite lab collection | NC_001604 |
| Bacteriophage FV3 | Gift from L. Truncaite lab, Truncaite *et al.*, 2012<sup>73</sup> | JQ031132 |
| Bacteriophage Alf5 | Gift from L. Truncaite lab, Alijošius *et al.*, 2017<sup>74</sup> | KX377933.1 |
| Bacteriophage P1 | ATCC collection, 25404-B1 | AF234172 |
| Bacteriophage lambda-vir | Gift from L. Truncaite lab collection | N/A |
| Bacteriophage Bas18 | BASEL, Maffei *et al.*, 2021<sup>75</sup> | MZ501095 |
| Bacteriophage Bas69 | BASEL, Maffei *et al.*, 2021<sup>75</sup> | MZ501049 |
| **Chemicals, peptides, and recombinant proteins** |  |  |
| Yeast Extract | Difco | 210941 |
| Tryptone | Difco | 211921 |
| Kanamycin sulfate | Sigma-Aldrich | 60615 |
| Carbenicillin | Sigma-Aldrich | C1389-10G |
| Ampicillin | Fisher Scientific | BP1760-25 |
| Streptomycin sulfate | Gibco | 11860038 |
| Anhydrotetracycline hydrochloride | Fisher Scientific | J66688.MB |
| L-(+)-arabinose | Thermo Fisher | A11921.30 |
| Ni<sup>2+</sup>-charged HiTrap chelating HP column | Cytiva | 17-0409-03 |
| StrepTrap XT column | Cytiva | 29401323 |
| HiLoad 16/600 Superdex 200 column | Cytiva | 28-9893-35 |
| Superdex 200 Increase 10/300 GL column | Cytiva | 28-9909-44 |
| B-PER™ Bacterial Protein Extraction Reagent | ThermoFisher | 78243 |
| Lysozyme | ThermoFisher | 89833 |
| 4',6-diamidino-2-phenylindole, dihydrochloride (DAPI) | Invitrogen | D1306 |
| Propidium iodide (PI) | Sigma-Aldrich | 537059 |
| DNase I, RNase free | ThermoFisher | 18047019 |
| RNase A/T1, DNase and protease free | ThermoFisher | EN0551 |
| FastAP Thermosensitive Alkaline Phosphatase | ThermoFisher | EF0651 |
| Roti-phenol/chloroform/isoamyl alcohol | Carl-Roth | A156 |
| 5PRIME Phase Lock Gel tubes | Quantabio | 733-2477 |
| T4 DNA polymerase | ThermoFisher | EP0062 |
| Exonuclease I | ThermoFisher | EN0581 |
| Streptavidin-coated MyOne C1 magnetic beads | Invitrogen | 65002 |
| Phusion Plus DNA polymerase | ThermoFisher | F630L |
| NaOH | Sigma-Aldrich | 221465 |
| Alcojet | Alconox | 1404-1 |
| Absolute ethanol | Honeywell | 32221 |
| Sulfobetaine silatrane | Gift from Chun-Jen Huang lab<sup>76</sup> | N/A |
| 1-(3-Aminopropyl)silatrane | This study<sup>77</sup> | N/A |
| Glacial acetic acid | Sigma-Aldrich | 695092 |
| Biotin-PEG4-NHS ester | Lumiprobe | 2893 |
| PBS (10X), pH 7.4 | Thermo Fisher | 70011044 |
| SYTOX Green Nucleic Acid Stain | Thermo Fisher | S7020 |
| TRIS | Carl Roth | 2449 |
| Hydrochloric acid | Carl Roth | 4625 |
| Potassium chloride | Carl Roth | 6781 |
| Traptavidin | This study<sup>78,79</sup> | N/A |
| FastDigest Bsp120I | Thermo Fisher | FD0134 |
| EDTA | Carl Roth | CN06 |
| Sodium chloride | Carl Roth | 9265 |
| Calcium chloride, dihydrate | Amresco | 0556 |
| Magnesium chloride hexahydrate | Carl Roth | 2189 |
| Streptavidin lyophilized, salt-free | SERVA | 35490 |
| **Critical commercial assays** |  |  |
| DNase Alert kit | IDT | 11-04-02-04 |
| NEBuilder HiFi DNA Assembly Master Mix | New England Biolabs | E2621 |
| QuikChange Lightning Multi Site-Directed Mutagenesis Kit | Agilent | 210516 |
| SPLIT RNA Extraction Kit | Lexogen | 008 |
| RiboCop for Gram negative Bacteria | Lexogen | 126 |
| Small RNA-Seq Library Prep Kit | Lexogen | 052 |
| CORALL Total RNA-Seq Library Prep Kit | Lexogen | 095 |
| **Deposited data** |  |  |
| Raw NGS data | This study,<br>https://www.ncbi.nlm.nih.gov/bioproject/?term=PRJNA1142218 | PRJNA1142218 |
| XauSPARDA apo | This study | 9QBQ |
| XauSPARDA-gRNA/tDNA-dsDNA | This study | 9QCC (EMD-53008) |
| EaeSPARDA apo | This study | 9QBP |
| XauSPARDA-ΔDREN | This study | 9QBL |
| **Oligonucleotides** |  |  |
| RNA oligos | IDT | See Table S1 |
| DNA oligos | IDT | See Table S1 |
| **Recombinant and genomic DNA** |  |  |
| pBADHisA | ThermoFisher | V43001 |
| pCDF | Tolia & Joshua-Tor<sup>37</sup> | N/A |
| pUC19 | Thermo Scientific | SD0611 |
| λ DNA | Thermo Scientific | SD0011 |
| plasmid DNA | This study | See Table S1 |
| *E. coli* BL21-AI genomic DNA | This study | CP047231.1 |
| **Software and algorithms** |  |  |
| Biotite | https://www.biotite-python.org/0.41.0/index.html | v0.41.0 |
| ColabFold | Mirdita *et al.* 2022<sup>80</sup> | v1.5.5-cuda12.2.2 |
| DALI server | http://ekhidna2.biocenter.helsinki.fi/dali/ | N/A |
| Python | https://www.python.org/ | v3.11.6 |
| Scikit-learn | https://scikit-learn.org/1.4/index.html | v1.4.2 |
| Seaborn | https://seaborn.pydata.org/ | v0.12.2 |
| SSDraw | https://github.com/simonas-asmontas/SSDraw | N/A |
| UCSF ChimeraX | https://www.cgl.ucsf.edu/chimerax/ | v1.8 |
| XDS | https://xds.mr.mpg.de/html_doc/downloading.html | v. Jan26,2018<br>BUILT=20180126 |
| POINTLESS | https://www.ccp4.ac.uk/ | v. 1.10.26 |
| AIMLESS | https://www.ccp4.ac.uk/ | v. 0.7.2:27/05/18 |
| TRUNCATE | https://www.ccp4.ac.uk/ | v. 7.0.061 |
| CCP4 | https://www.ccp4.ac.uk/ | v. 7.1.002 |
| Auto-Rickshaw | http://www.embl-hamburg.de/Auto-Rickshaw/ |  |
| MOLREP | https://www.ccp4.ac.uk/ | v. 11.6.04 |
| REFMAC | https://www.ccp4.ac.uk/ | v. 5.8.0258 |
| PHENIX | https://www.phenix-online.org/ | v.1.20-4459 |
| COOT | https://www2.mrc-lmb.cam.ac.uk/personal/pemsley/coot/ | v. 0.9 |
| EPU | Thermo Fisher Scientific | v.3 |
| CryoSPARC | https://cryosparc.com/ | v.4.4.1 |
| CisTEM | https://cistem.org/ | v.1.0.0-beta |
| USalign | Zhang *et al.* 2022<sup>81</sup> | v20240319 |
| VoroContacts | https://bioinformatics.lt/wtsam/vorocontacts | v.1.22 |
| AdapterRemoval | Schubert *et al.* 2016<sup>82</sup> | v2.3.0 |
| bedtools | Quinlan *et al.* 2014<sup>83</sup> | v2.26.0 |
| BWA-MEM | Li *et al.* 2009<sup>84</sup> | v0.7.17-r1188 |
| csv2transfac.pl | https://github.com/agryb auskas/argonaute-bound-rna-manuscript | v0.1.2 |
| FastQC | https://www.bioinformatics.babraham.ac.uk/projects/fastqc/ | v0.11.8 |
| fragmentation-bias.jl | https://github.com/agryb auskas/argonaute-bound-rna-manuscript | v0.1.2 |
| RSeQC | Wang *et al.* 2012<sup>85</sup> | v4.0.0 |
| SAMtools | Danecek *et al.* 2021<sup>86</sup> | v1.7 |
| WebLogo | Crooks *et al.* 2004<sup>87</sup> | v3.7.4 |
| Anaconda.Navigator | https://www.anaconda.com/products/navigator | v.2.6.0 |
| Jupyter notebook | https://jupyter.org/ | v.7.0.8 |
| microEye | Alsamsam, M. N. microEye, https://github.com/samhi tech/microEye [Computer software] | v2.3.1 |
| Fiji 2.16.0 | Schindelin *et al.* 2012<sup>88</sup> | v1.54p |
| Mars | Huisjes *et al.* 2022<sup>89</sup> | N/A |
| Igor Pro 9.0.5.1 | WaveMetrics, Inc. | Build 56551 |

### EXPERIMENTAL MODEL AND STUDY PARTICIPANT DETAILS

#### DH5α E. coli

DH5α *E. coli* (K-12 F– *endA1 glnV44 thi-1 recA1 relA1 gyrA96 deoR nupG Φ80dlacZ*Δ*M15* Δ*(lacZYA-argF)U169, hsdR17(rK– mK+),* λ–) was used for cloning purposes.

#### MG1655 E. coli

MG1655 *E. coli* (F- λ- *ilvG- rfb-50 rph-1*) was used for bacteriophage assay.

#### BL21-AI E. coli

BL21-AI *E. coli* (B F– *ompT gal dcm lon hsdSB(rB–mB–) [malB+]*K-12(λS) *araB::T7RNAP-tetA*) was used for *in vivo* assays and protein expression and purification, protein-nucleic acids co-purification.

#### DH10B E. coli

DH10B *E. coli* (K-12 F– *endA1 recA1 galE15 galK16 nupG rpsL* Δ*lacX74 Φ80lacZΔM15 araD139* Δ*(ara,leu)7697 mcrA* Δ*(mrr-hsdRMS-mcrBC) StrR* λ–) was used for protein expression and purification.

All *E. coli* strains were cultivated in LB media or on LB agar plates (with 1.5 % agar) supplemented with antibiotics according to plasmid resistance (ampicillin/carbenicillin (Amp/Cb): 100 μg/mL; kanamycin (Kn): 25 μg/mL, tetracycline (Tc): 10 μg/mL; streptomycin (Str): 50 μg/mL). For experiments involving bacteriophages all cultivations were performed in LB media at 37 °C unless stated otherwise. When required, expression was induced by adding arabinose (for *in vivo* assays and protein expression and protein or protein-nucleic acid purification to final concentrations of 0.2 % (w/v)).

### METHOD DETAILS

#### Cloning and mutagenesis

Plasmids with WT XauSPARDA, EaeSPARDA, TmuSPARDA and NbaSPARDA genes used for protein expression and *in vivo* assays were obtained using whole gene synthesis and cloning service provided by Twist Bioscience and are listed in Table S1. Genes were cloned into pBAD expression vectors under a P_BAD_ promoter inducible with arabinose which allowed for expression of the proteins that are linked to an N-terminal His_6_ or/and Strep tags for protein purification. Point and multiple mutations were introduced using the NEBuilder HiFi DNA Assembly Master Mix (New England Biolabs, E2621) or QuikChange Lightning Multi Site-Directed Mutagenesis Kit (Agilent, 210516) and heat-shock transformation into DH5α *E. coli* cells.

To swap streptomycin resistance for kanamycin in the pCDF plasmid, plasmid pCDF was cleaved with NheI and Eco81I (ThermoFisher, FD0374) and isolated using a runView electrophoresis system (Cleaver Scientific). The purified fragments were then ligated into pCDF to yield a pCDF_Kn plasmid.

All plasmids used in this study were sequenced and confirmed by a full-plasmid sequencing service (SeqVision); links to DNA and protein sequences are presented in Table S1.

#### Phage restriction assay

*E. coli* MG1655 (ATCC 47076) cells carrying empty vector pBADHisA and pBADHisA plasmid expressing WT SPARDA system were used for phage infection. Liquid culture phage infection experiments were performed using *E. coli* bacteriophages T2, T4, T6, T3, T5, T7, FV3, Alf5, P1, lambda-vir, Bas18 and Bas69 (see Key Resources Table). After overnight incubation, the liquid suspension of pAgo-lacking (as control used empty vector pBAD) and pAgo-containing *E. coli* cells were diluted 1:40 in LB medium, grown until reached OD_600_∼0.5, then SPARDA expression induced by adding 0.2% L-arabinose (L-Ara), cells were grown for 2 h at 37 °C, and dispensed into a 96-well plate by 90 μl (normalized to OD_600_=0.5 and diluted 5×). Then 10 μl of phage lysate was added to each well in a multiplicity of infection 0.05. Optical density measurements at a wavelength of 600 nm were taken every 15 min using a Clariostar plate reader. Each experiment was performed in three biological replicates, and visualizations were performed using Anaconda.Navigator (v.2.6.0) in the Jupyter notebook environment (v.7.0.8) in graph form with average growth curve with standard deviation shown in shading around the curve.

#### Cytotoxicity assays

*E. coli* BL21-AI strain cells were heat-shock transformed with plasmids: 1) pBADHisA plasmid encoding XauSPARDA or EaeSPARDA (either WT or mutant variants), - under the control of araBAD promoter, and/or 2) pCDF plasmid. Protein expression in selected single or double transformants was induced by adding L-Ara (final concentration of 0.2% (w/v)) into liquid LB culture, and cells were harvested after 2-4 h. Then cells were used: 1) for serial dilutions in plate assay: optical density (OD_600_) equalized, cultures were serially diluted, and aliquots were spotted on an agarose LB medium containing 50 μg/ml ampicillin, 25 μg/ml tetracycline and/or 25 μg/ml streptomycin and/or L-Ara and were grown overnight at 37 °C. The viability of uninduced cells was measured as a control. Plate pictures were taken with UVITEC imaging system (Cambridge) and colony forming units (CFU/ml) were counted on the last visible dilution; 2) to count dead cells using flow cytometer: 100-300 μl of cells were collected by centrifuging 4g 5 min, washed with PBS and dyed with propidium iodide (Thermo Fisher Scientific, P3566) in PBS (final concentration 10 μg/ml) for 15-30 min on ice, the prepared samples were analyzed using BD FACSymphony A1 flow cytometer (BD Biosciences, USA), collecting forward scatter (FSC; height (H), width (W), and area (A)), side scatter (SSC; H, W, A), and fluorescence emission in the PE-CF594 channel (excitation 561 nm, emission 610/20 nm) for PI-stained cells. A minimum of 20,000 events per sample were recorded. Data were analyzed using FlowJo V10 software (BD Biosciences, USA); 3) to measure DNA amount in cells: 100-300 μl of cells were fixed by adding 3:7 ethanol (100 %), fixed overnight at 4 °C, collected by centrifuging 4g 5 min, washed with PBS and dyed with DAPI (Thermo Fisher Scientific, D1306) in PBS (final conc. 10 μg/ml) for 1 h on ice, washed with PBS and resuspended in PBS for analysis. The prepared samples were analyzed using BD FACSymphony A1 flow cytometer, collecting forward scatter (FSC; H, W, A), side scatter (SSC; H, W, A), and fluorescence emission in the BV421 channel (excitation 405 nm, emission 450/50 nm) for DAPI-stained cells. A minimum of 20,000 events per sample were recorded. Data were analyzed using FlowJo V10 software. DNA amount was compared between cells having the same size plasmids: pBADHisA with WT or inactive mutant E37A SPARDA genes; 4) for Western blot analysis using His_6_-Tag antibody (ThermoFisher, MA1-21315) or StrepTag antibody (ThermoFisher, MA5-37747) depending on Tag that Xau or EaeSPARDA proteins have.

Each experiment was performed in three biological replicates, and visualizations were performed using Anaconda.Navigator (v.2.6.0) in the Jupyter notebook environment (v.7.0.8) in bars form of average mean and experimental points on the bars.

#### Purification of protein complexes

All cells containing proteins of interest were collected by centrifugation, resuspended in buffer1 (20 mM Tris-HCl (pH 8.0 at 25 °C), 1000 mM NaCl, 5 mM 2-mercaptoethanol, 2 mM phenylmethylsulfonyl fluoride) and disrupted by sonication. After removing cell debris by centrifugation, the supernatants with XauE37A, XauD41A, XauQ62A, XauK64A, XauHSH, DREN, XauWT, XauAD1, XauAD2, EaeE37A, EaeWT proteins were loaded on a Ni^2+^- charged HiTrap chelating HP column (Cytiva). Proteins were eluted with a linear gradient of increasing imidazole concentration (from 25 to 500 mM) in buffer2 (20 mM Tris-HCl (pH 8.0 at 25 °C), 500 mM NaCl, 5 mM 2- mercaptoethanol). Fractions with proteins were further loaded on the HiLoad 16/600 Superdex 200 gel filtration column (Cytiva) equilibrated with buffer2. The supernatants with XauFA1, FA2, FA3, FD1, FD2, and FD3 proteins were first loaded on a Ni^2+^-charged HiTrap chelating HP column (Cytiva) similar to those described above. Fractions with proteins of interest were loaded onto the StrepTrap XT column (Cytiva), washed with buffer2, and elution of specifically bound protein complexes was performed with 50 mM biotin solution. XauΔLSL, TmuSPARDA, NbaSPARDA proteins were purified using a Ni^2+^-charged HiTrap chelating HP and StrepTrap XT columns (Cytiva) as described previously and additionally was used the HiLoad 16/600 Superdex 200 gel filtration column (Cytiva) equilibrated with buffer2. Fractions with all purified proteins were pooled and dialyzed against a buffer containing 20 mM Tris-HCl (pH 8.0 at 25 °C), 500 mM NaCl, 2 mM dithiothreitol (DTT), and 50% (v/v) glycerol.

#### Size-exclusion chromatography with multi-angle light scattering

Size-exclusion chromatography with a multi-angle light scattering (SEC-MALS) of all protein complexes and their complexes with nucleic acids (200 μl sample containing 2 μM of protein, 2 μM of gRNA/tDNA, 0.5 μM of TK49/MZ949 double-stranded DNA (pre annealed oligoduplex)) were carried out at room temperature using a Superdex 200 Increase 10/300 GL column (Cytiva) equilibrated with buffer (20 mM Tris–HCl (pH 8.0 at 25 °C), 150 mM NaCl, 2 mM MgCl_2_, 1 mM DTT), at 0.4 ml min^−1^ flow rate. The light scattering signals were monitored on a miniDawn TREOS II detector, and concentrations of all samples were measured using an Optilab T-rEX refractive index detector (Wyatt Technologies). Data were analyzed in Astra software (Wyatt Technologies) using a dn/dc value of 0.185 ml g ^−1^.

#### Nucleic acid extraction and analysis

To obtain Xau and EaeSPARDA-bound nucleic acids, *E. coli* DH10B was transformed with two plasmids: 1) pBAD_His_Strep_His_XauSPARDA or pBADHisA_TwinStrep_TEV_EaeSPARDA, and 2) pCDFDuet_Kn plasmids. Cells were grown at 37 °C in LB medium in the presence of 50 μg/ml ampicillin, 25 μg/ml streptomycin and 25 μg/ml kanamycin until OD_600_ = 0.5 was reached. Then, expression was induced by adding 0.2% (w/v) L-arabinose, and cells were harvested after 2 h. Cells were disrupted by incubating for 20 min at room temperature in lysis reagent B-PER (ThermoFisher, 78243) containing 3 mg/ml lysozyme (ThermoFisher, 89833). The XauSPARDA-NA and EaeSPARDA-NA complexes were purified using a StrepTactin column; all buffer solutions contained 100 mM NaCl. Xau and EaeSPARDA-bound nucleic acids were purified using phenol-chloroform isolation as described in Zaremba *et al*., 2022^15^: 800 μL of Roti-phenol/chloroform/isoamyl alcohol (Carl-Roth, A156) was added to the 800 μL of purified protein-NA fractions in 5PRIME Phase Lock Gel tubes (Quantabio, 733-2477). The upper aqueous phase was isolated, and 0.1 volumes of 1M sodium acetate, 3 volumes of 100 % ethanol, and 10 μL glycogen (ThermoFisher, R0561) were added. This mixture was vortexed briefly and incubated at −20 °C for 20 hours. Samples were centrifuged for 20 min, and the supernatant was removed from the pellet. The pellet was washed with cold (−20 °C) 70% ethanol. The pellets containing the copurified nucleic acids were dried for 20 min at room temperature and resuspended in 30 μL water (free of nucleases).

Co-purified nucleic acids were dephosphorylated with FastAP Thermosensitive Alkaline Phosphatase (ThermoFisher, EF0651) and [γ-32P]-ATP (Revvity) labeled with T4 polynucleotide kinase (PNK) (ThermoFisher, EK0031). Labeled nucleic acids were incubated with nucleases (ThermoFisher DNase I, 18047019, RNase A/T1, EN0551) for 60 min at 37 °C. After nuclease treatment, samples were mixed with RNA Gel Loading Dye (ThermoFisher, R0641), heated for 5 min at 95 °C, and resolved on 20% denaturing (8 M Urea) polyacrylamide gels. The molecular weight marker used for RNA size identification was the Decade Marker System (Ambion, AM7778). Radioactivity was captured from gels using phosphor screens and imaged using a Typhoon FLA 7000 laser scanner (GE Healthcare). Each experiment was performed in three biological replicates.

In a control sample a total RNA from induced cells was extracted using a SPLIT RNA Extraction Kit (Lexogen, 008). Then, rRNA was removed using RiboCop for Gram-negative Bacteria (Lexogen, 126). Next, RNA samples were sequenced, as noted below.

#### RNA sequencing and analysis

RNA samples (extracted from protein-NA complexes) were converted to DNA libraries using the Small RNA-Seq Library Prep Kit (Lexogen, 052). The concentration and quality of libraries were measured with a Qubit Fluorometer (ThermoFisher) and a 2100 Bioanalyzer (Agilent).

Libraries were sequenced using Illumina MiniSeq sequencing with single-end reads and 75 bp read length. Single-end reads were processed by trimming adapters with AdapterRemoval. Then, the processed reads were aligned to the *E. coli* str. K12 substr. DH10B genome (GenBank: CP000948.1) and the additional pBAD_His_Strep_His_XauSPARDA or pBADHisA_TwinStrep_TEV_EaeSPARDA and pCDF_Kn plasmids (Table S1) using BWA-MEM. To avoid filtering out shorter reads during the alignment process, aligned reads with MAPQ values greater or equal to 15 were chosen. FastQC was used for read quality control and SAMtools – for indexing, sorting, and analyzing alignment files. Custom scripts (fragmentation-bias.jl and csv2trasnfac.pl) in combination with Weblogo were used to produce nucleotide frequency plots. The custom scripts had to be implemented to ensure that only aligned reads would be used for nucleotide frequency analysis. Gene enrichment analysis was performed with bedtools and FPKM_count.py of the RSeqQC package.

The control DNA library of total RNA was prepared using the CORALL Total RNA-Seq Library Prep Kit (Lexogen, 095). The concentration and quality of the library were measured with a Qubit Fluorometer (ThermoFisher) and a 2100 Bioanalyzer (Agilent) according to the protocol of the manufacturer. Control DNA libraries were sequenced using Illumina MiSeq sequencing with single-end reads and 150 bp read length. The read processing, alignment, and alignment analysis were analogous to those samples from Illumina MiniSeq sequencing.

Sequencing was performed in two biological replicates.

#### In vivo DNA cleavage analysis

To obtain XauSPARDA fragmented DNA *E. coli* BL21-AI was transformed with WT or mutant pBADHisA_XauSPARDA and/or pCDF plasmids. Cells were grown at 37 °C in LB medium in the presence of 50 μg/ml ampicillin, 25 μg/ml streptomycin (if with pCDF), and 25 μg/ml tetracycline until OD_600_ = 0.5 was reached. Then, expression was induced by adding 0.2% (w/v) L-arabinose, and cells were harvested after 2 h (with pCDF) or overnight (∼20 h, if without pCDF). DNA was purified using GeneJET Plasmid Miniprep Kit (Thermo Scientific, K0502). The purified DNA was loaded on an agarose gel with EtBr for visualization, then electrophoresis was performed, and DNA was imaged using the UVITEC imaging system (Cambridge). Then, the DNA (with pCDF) library was prepared using CROFT-seq method^36^ as follows. First, purified DNA (100 ng) was treated with T4 DNA polymerase (Thermo Fisher Scientific, EP0062) and then ligated to a biotinylated adapter. Then, ligation products were treated with Exonuclease I (Thermo Fisher Scientific, EN0581) to remove the unligated adapter. Ligated DNA was immobilized on streptavidin-coated MyOne C1 magnetic beads (Invitrogen, 65002), and the non-biotinylated DNA strand was removed by adding 125 mM NaOH. The remaining biotinylated DNA strand was used as a template to synthesize a complementary DNA strand using DNA oligonucleotide and T4 DNA polymerase (Thermo Fisher Scientific, EP0062). The resulting DNA was removed from magnetic beads and amplified with Phusion Plus DNA polymerase (Thermo Fisher Scientific, F630L) using indexed DNA oligonucleotides. Prepared DNA libraries were sequenced on an Illumina MiSeq system with 150 bp pair-end reads. Single-end reads were aligned to the *E. coli* strain K12 substr. BL21-AI genome (GenBank: CP047231.1) and the additional pBADHisA_XauSPARDA and pCDFDuet plasmids. The analogous bioinformatics workflow was performed as for RNA samples. Sequencing was performed in two biological replicates.

#### Analysis of in vitro activities of SPARDA

The nuclease activity of SPARDA was analyzed using synthetic oligonucleotides, which served as the guide, target, and substrate (Table S1). The SPARDA complex (250 nM final concentration) was preincubated with guide nucleic acid (as specified in figures) (500 nM) for 10 minutes at 37°C in 1× Tango buffer (33 mM Tris-acetate (pH 7.9 at 37 °C), 10 mM magnesium acetate (or any appropriate divalent cation as specified in the figures (10 mM of ions in all reactions), 66 mM potassium acetate, 0.1 mg/ml BSA (ThermoFisher, BY5). Subsequently, 750 nM target nucleic acid (as specified in Figures 2 and S3) was added, and the samples were incubated for 10 minutes. Collateral substrate oligonucleotides were added to the 100 nM final concentration (ssDNA, dsDNA, ssRNA, dsRNA, or RNA/DNA hybrid, as indicated in Figures 2 and S3). Following the addition of the substrates, either kinetic or endpoint experiments were conducted. In most experiments, the reactions were performed for 60 minutes at 37 °C unless otherwise indicated. STOP buffer containing 6 M UREA, 125 mM EDTA, and 4x TBE was added to stop the reactions. The reaction products were separated by 19% denaturing urea PAGE and visualized by SYBR GOLD staining. Nuclease activity reactions with larger substrates: supercoiled pUC19 plasmid DNA (Thermo Scientific, SD0061), λ phage DNA (Thermo Scientific, SD0011) and *E. coli* genomic DNA (obtained by the GeneJET Genomic DNA Purification Kit (Thermo Scientific, K0721) (all at 2 nM concentration) were performed with same synthetic oligonucleotides (Table S1) and reaction products were separated by 0.8% agarose gel and stained with SYBR GOLD. The gels were scanned using a Typhoon FLA 7000 scanner (GE Healthcare). Each experiment was performed in triplicates.

#### Collateral cleavage assay

Reaction mixtures with a volume of 50 μl were prepared with the following final concentrations: 1 μM WT XauSPARDA/EaeSPARDA or mutant, 0.5 μM gRNA (Table S1), 0.75 μM tDNA (Table S1) in 1× Tango buffer (33 mM Tris-acetate (pH 7.9 at 37 °C), 10 mM magnesium acetate, 66 mM potassium acetate, 0.1 mg/ml BSA (ThermoFisher, BY5), 1 mM DTT, 0.1 μM DNaseAlert (IDT, 11-04-02-04). Reactions with the RNA guide were pre-incubated for 10 min at 37 °C, then tDNA was added, and the mixture was incubated for 10 min at 37 °C. Finally, collateral cleavage reactions were initiated by adding DNaseAlert dsDNA substrate (IDT, 11-04-02-04). The assay was performed by incubating samples in a 96-well flat-bottom black polystyrene plate (Corning, 3881). The fluorescence signal (λex 533±4 nm; λem 559±4 nm) was measured in a CLARIOstar Plus microplate reader (BMG LABTECH) for 120 min at 37 °C with measurements taken every 1 or 2 min. Each experiment was performed in triplicate. Bar graphs were generated using GraphPad Prism 8 (v.10.3.1). Each dot in the figures is a mean from three independent experiments which then were mostly normalized to standard conditions of the reaction – WT protein with gRNA Mz-1800 and fully complementary tDNA Mz-1455 unless specified otherwise. Line graphs were generated with Anaconda.Navigator (v.2.6.0) in the Jupyter notebook environment (v.7.0.8).

#### Crystallization and structure determination

For crystallization, apo XauSPARDA and Se-methionine modified ΔDREN mutants were concentrated to 5 and 6.5 mg/ml, respectively, and crystals were grown in a cold room. Reservoir solutions are described in Table S2. Relatively large crystals of apo XauSPARDA were easy to grow reproducibly, but poor diffraction was observed along one of the crystal axes. Numerous tests were conducted until the crystal diffracted to 3 Å was found. Se-methionine ΔDREN crystallization was set up using a microcrystal seed solution.^90,91^ Apo EaeSPARDA was concentrated for crystallization to 4 mg/ml. All datasets were collected after a few seconds of rinsing in a cryoprotection buffer (Table S2).

Diffraction data were processed with XDS v.Jan26,2018 BUILT=20180126^92^, POINTLESS v. 1.10.26, AIMLESS v.0.7.2:27/05/18, TRUNCATE v.7.0.061 and other CCP4 v.7.1.002 tools.^93^ The structure of the Se-Methionine ΔDREN mutant was solved using the 3W-MAD protocol of Auto-Rickshaw: the EMBL-Hamburg automated crystal structure determination platform.^94^ The input diffraction data were prepared and converted for use in Auto-Rickshaw using programs of the CCP4 suite. FA values were calculated using the program SHELXC v.006/2.^95^ Based on an initial analysis of the data, the maximum resolution for substructure determination and initial phase calculation was set to 3.2 Å. All of the 19 heavy atoms requested were found using the program SHELXD v.2011/5.^96^ The correct hand for the substructure was determined using the program ABS^97^, and initial phases were calculated by SHELXE v. 2006/1.^98^ The twofold non-crystallographic symmetry (NCS) operator was found using the program RESOLVE^99^. Density modification, phase extension and NCS-averaging were performed using the program DM v.7.0.072^100^. Initial phases for EaeSPARDA and XauSPARDA structures were obtained by molecular replacement by MOLREP v.11.6.04^101^ using ΔDREN mutant structure as an initial model. All models were inspected and rebuilt in COOT v.0.9^102^. Apo XauSPARDA and EaeSPARDA complexes were refined with PHENIX 1.20_4459^103^, whereas ΔDREN mutant structure was refined with REFMAC v.5.8.0258.^104^

#### Cryo-EM sample preparation, data collection, and model refinement

For cryo-EM analysis, XauSPARDA apo dimers in the buffer containing 20 mM Tris-HCl (pH 8.0 at 25 °C), 100 mM NaCl, and 2 mM CaCl_2_ were first activated by mixing with 17 nt gRNA in the ratio 1 (monomer):1.2 (gRNA). After 10 min incubation at RT 17 nt tDNA complementary to guide, the ratio 1:1 to monomers was added. Oligomerization was started by the addition of dsDNA substrate in the ratio 1:1 to activated XauSPARDA monomers and centrifuged to remove large aggregates. Samples of 3 μl (1.96 mg/ml of protein) were applied on both sides of glow-discharged Copper R1.2/1.3 300 mesh holey carbon grid (Quantifoil) at 95% humidity and 4°C, blotted and plunge-frozen in liquid ethane using Vitrobot Mark IV (Thermo Fisher Scientific, Waltham, USA).

Cryo-EM data of the activated XauSPARDA filaments were collected at Glacios microscope (Thermo Fisher Scientific, USA) equipped with FEI Falcon III (4k x 4k) direct electron detector. Images were collected as movies consisting of 30 frames with an overall exposure of 46.33 sec using EPU v. 3 software (Thermo Fisher Scientific, USA) in the counting mode at a nominal magnification of 92000× with a defocus ranging from –1 to –2 μm. A dose rate of 0.8 e/Å^2^/sec and an overall exposure time of 46.33 sec resulted in a final dose of 30 e/Å^2^.

Cryo-EM data were processed using CryoSPARC v.4.4.1.^105^ The summary of image processing and 3D volume refinement is presented in Figure S5 and Table S2. The particles were template-picked using an *ab initio-* generated volume obtained in the preliminary dataset. We used an extraction box of 512 px. After repetitive 2D classification a subset of 1603031 filament-like particles was selected (Figures 3A, 3B, and S5) and sorted by *ab-initio* reconstruction followed by heterogenous refinement. The resulting set containing 204199 particles was refined by homogenous refinement protocol and local refinement within the mask that defined a presented model.

The cryo-EM map was sharpened using CisTEM.^106^ The crystal structure of apo XauSPARDA was used as an initial model and inserted into a cryo-EM map using CHIMERA v.1.17.1.107. The Model was refined using phenix.real_space_refine v.1.18.2-3874^107^ and rebuilt in COOT.^102^

#### Computational protein structure analysis

3D superposition of XauSPARDA monomers from Apo and Filament structures was done by using USalign.^81^ Prior to Beta Relay register shift analysis, the secondary structure elements were recalculated using DSSP,^108^ implemented in USCF ChimeraX.^109^

Protein structure modeling was performed using a Docker container of ColabFold.^80^ The default settings were used for all the modeling instances except for the “num_models” parameter, which was set to 5 when computing the truncated dimer models and the effector domain tetramer models. The computed models were not relaxed for the downstream analyses.

Clustering of structural models was carried out using the Agglomerative Clustering algorithm, implemented in Scikit-learn.^110^ For the distance metric, we used distances derived from pairwise all-versus-all TM-scores calculated with USalign via the formula: distance = 1 – TM-score. The distance threshold was set to 0.2 (corresponding to a pairwise TM-score of 0.8), while the linkage parameter was set to “single”. The clustering plots were drawn using Seaborn.^111^

Structure-based pairwise alignment of pAgo protein structures determined in different states was done using the DALI web server.^112^ The local structure differences between these pAgo states were evaluated using CAD-score.^113^ Prior to CAD-score calculation, the structure pairs were filtered to contain the intersection of their residues using Biotite.^114^

Structure-based multiple sequence alignment (MSA) of pAgo MID-PIWI chains was constructed using USalign. The MSA was derived from the top-ranked ColabFold models. The MSA was visualized using a customized version of SSDraw.^115^

Analysis of contacts and interaction surfaces was performed by VoroContacts^116^.

#### Single-molecule experiments Preparation of Modified Glass Coverslips

Cleaning and chemical modification of high-precision cover glasses (Paul Marienfeld, 0107242) were performed in Hellendahl staining jars. Glass coverslips were first rinsed in Milli-Q (MQ) water and immersed in 1% (w/v) Alcojet aqueous solution. Such staining jar was then put in ultrasonic cleaner (GT SONIC VGT-6250), and 5 min-long sonication was performed. After rinsing the glass coverslips in MQ water thrice, they were kept in 1 M NaOH for 1 h. Next, they were rinsed thrice in MQ water and left to dry on a lab table for 20 min. Finally, glass coverslips were heated in a laboratory oven for 20 min at 120°C and stored at room temperature (RT) until depleted.

To passivate and functionalize the surface of these cleaned cover glasses, they were initially treated with air plasma (∼260 mTorr, highest power setting) for 10 min in a tabletop plasma cleaner (Diener electronic, ZEPTO-W6). Immediately after this step, glass coverslips were immersed in the absolute ethanol solution containing 0.9 mM sulfobetaine silatrane, 0.1 mM 3-aminopropylsilatrane, 10% (v/v) MQ water, and 2% (v/v) glacial acetic acid. Such staining jar was then placed on a laboratory rocker-shaker and kept there for 30 min at RT. After rinsing the glass coverslips in absolute ethanol and MQ water once each, they were placed on a lab table to dry for 15 min and then transferred to a laboratory oven for 15 min-long curing at 110°C. PEGylation of silatranized glass coverslips was performed after letting them cool down for 10 min. Glass coverslips were taken out of the staining jar and laid horizontally on a holder made from pipette tip rack. PEGylation ‘sandwiches’ were formed – a single drop of 60 µL of 17 mM biotin-PEG_4_-NHS ester in 1X PBS (pH 7.4 at 25°C) was applied to the center of a cover glass and then another silatranized glass coverslip was added on top. After leaving such ‘sandwiches’ for 30 min at RT and labeling the non-PEGylated side of the glass with a marker pen, they were disassembled using tweezers, washed with MQ water, dried under a gentle N_2_ gas flow, and stored in separate 50 mL conical centrifuge tubes at −20°C for no longer than 1 month.

#### Total Internal Reflection Fluorescence (TIRF) Microscopy

Our home-built open-source single-molecule (SM) localization fluorescence microscope miEye,^117,118^ along with its dedicated Python-based software microEye, were employed to conduct all SM fluorescence microscopy experiments in this work. To maintain the desired elevated temperature of the sample during experiments, a custom sample heating system was constructed and adapted for the miEye. This system consisted of four main parts:

1. A flexible polyimide foil heater (Thorlabs, TLK-H) attached to the 3D printed vertical side wall (Clear resin, FormLabs) of the L-shaped sample holder mounted to the manual Z stage.
2. Another foil heater (Thorlabs, TLK-H) wrapped around the objective (above its correction collar) and fixed with a hook-and-loop fastener (Velcro).
3. A heater and thermoelectric cooler temperature controller (Thorlabs, TC300B) connected to both foil heating elements using two 6-pin male-to-male Hirose connector cables (Thorlabs, HR10CAB1).
4. A 3D printed housing (Clear resin, FormLabs) and PMMA lid, which can both be added to the sample holder using neodymium mounting magnets, to create a semi-hermetic container around the sample. Imaging was performed using Nikon APO TIRF 60 × 1.49 NA objective with its correction ring set for a temperature of 37 °C and cover glass thickness of 0.17 mm.

The miEye was arranged in a single-mode fiber-based excitation scheme with quad-line polychroic mirror Di03-R405/488/561/635-t3-25×36 (AHF Analysentechnik, F73-866S) installed in the microscope’s main body to direct the excitation beam towards the sample. Samples were illuminated in TIRF mode using either 488 nm or 638 nm wavelength lasers (Integrated Optics), with their power set below 1 W/cm2 to excite SYTOX Green (SG) and ATTO647N fluorophores, respectively. Additional extension of miEye’s emission pathway was done, enabling simultaneous imaging of two colors on a single detector. To split the collected fluorescence into two spectral components, a 550 nm long-pass dichroic mirror (Thorlabs, DMLP550R) was inserted in the Fourier space of the 4f configuration of the microscope. The 525/45 and 697/75 band-pass filters were utilized for isolating the fluorescence light emitted by SG and ATTO647N dyes, respectively. Both spectral channels were directed to a single industrial CMOS camera (Allied Vision Technologies, Alvium 1800 C-511m) by the knife-edge right-angle prism mirror (Thorlabs, MRAK25-P01). Upstream of this prism mirror, achromat lenses (f = 100 mm) were installed to focus fluorescence emission onto the sensor plane. In such a setting, the projected pixel size was estimated to be ∼114.17 nm in XY. Image series were acquired using 300 ms exposure time of the camera with the corresponding frame rate of 3.3 Hz.

#### DNA Flow-Stretch Assays

The prepared glass coverslips were assembled into flowcells by attaching the PEGylated side of the cover glass to the air plasma-treated clean 6-channel plastic slide (Ibidi, 80606) via a double-sided sticky tape (3 M, 9088– 200) spacer. After inserting two cut pieces of silicone tubing (Carl Roth, CH24.1) into the openings of a selected slide’s channel, shortened 1 mL pipette tip (inlet’s reservoir) was embedded into one piece, and, using PFA tubing (IDEX Health & Science, 1512L) as a joint, the other piece was connected to the peristaltic pump (Ismatec ISM831C Reglo Digital) equipped with Tygon LMT-55 tubing (Fisher Scientific, 15532181) with its other end (outlet) directed to a waste container. All TIRF microscopy experiments conducted in this study were performed at 37°C and consisted of the same initial steps described next.

The channel of an assembled flow cell was firstly filled with 200 µL of buffer A (33 mM Tris-HCl (pH 7.9 at 25°C), 66 mM KCl). Then, 100 µL of 50 nM traptavidin in buffer A were added, incubated for 10 min, and washed away twice with 200 µL of buffer A. Following an injection of 100 µL of 25 nM biotinylated DNA oligonucleotide in buffer A and subsequent 10 min incubation, the channel was again flushed twice with 200 µL of buffer A. Next, 100 µL of 35 pM Bsp120I-treated λ DNA in buffer A were loaded into the channel. After a 15 min-long incubation, one-time washing with 200 µL of buffer A and three times with buffer B (6.6 mM Tris-HCl (pH 7.9 at 25°C), 13.2 mM KCl) was performed. To visualize single-end tethered λ DNA molecules, they were stained by injecting 300 µL of 10 nM SG in buffer B. Fluorescence movies of 35 frames were recorded, during which the flow of this SG-containing buffer was started (at the rate of 1 mL/min) and then terminated after 7 frames.

Digestion of λ DNA with Bsp120I restriction enzyme was carried out by treating 1.4 nM λ DNA with 1 µL of Bsp120I in 1X FastDigest Buffer for 1 h at 37°C. To stop the reaction, EDTA in a final concentration of 10 mM was added to the reaction mixture, which was then heated at 80°C for 20 min. Between the experiments, such prepared λ DNA sample was stored at 4°C for a maximum period of 1 month. When working with solutions containing bacteriophage λ DNA, wide orifice pipette tips (Thermo Fisher, 9405020) were always used to reduce the physical shearing and fragmentation of this genomic DNA. Bsp120I-catalyzed cleavage of λ DNA was a prerequisite mainly for disrupting concatemers present in the original λ DNA stock,^119^ thus ensuring smoother Gaussian distribution of individual λ DNA molecules’ length and enabling more efficient immobilization of this DNA on the surface.

When these initial steps of the experiment were complete, 441 nM SPARDA-gRNA/tDNA complex was assembled in buffer C (6.4 mM Tris-HCl (pH 8.0 at 25°C), 13.2 mM NaCl, 2 mM CaCl_2_) in a separate tube. Equimolar concentrations of protein and respective nucleic acids were used for this complex assembly. Upon mixing SPARDA with gRNA in buffer C, pre-warming at 37°C for 5 min was performed, and this incubation step was repeated after adding tDNA to the mixture. The assembled SPARDA-gRNA/tDNA complex was then diluted in buffer B, supplemented with 10 nM SG, to the final concentration of 67 nM. In terms of salt concentration, this yielded a solution with the following final composition: 6.6 mM Tris-HCl (pH 7.9 at 25°C), 11.2 mM KCl, 1.8 mM NaCl, 0.27 mM CaCl_2_. To study the compaction of single-end immobilized λ DNA, 130 µL of such diluted SPARDA-gRNA/tDNA complex were added to the channel. Imaging was done 20 min post-injection, and the response of individual λ DNA molecules to the applied buffer flow (with a rate of 1 mL/min) was assessed by running the same solution containing this diluted complex through the channel. The same experimental conditions were used for all SPARDAs investigated at the SM level, except (the final concentration of the diluted assembled SPARDA-gRNA/tDNA complex and incubation time after its injection into the channel are indicated in the brackets): XauD41A (801 nM; 20 min), WT NbaSPARDA (67 nM; 50 min), WT EaeSPARDA (467 nM; 50 min), WT TmuSPARDA (67 nM; 50 min). WT EaeSPARDA-gRNA/tDNA complex was diluted and examined in 10 nM SG-containing buffer A. In the experiments that involved regeneration of λ DNA compacted by the activated WT XauSPARDA, decompaction of λ DNA molecules was accomplished by washing the channel a minimum of five times with buffer A and waiting for at least 30 min before proceeding with any further imaging.

Cleavage of compacted λ DNA molecules was triggered by introducing Mg^2+^ ions into the channel. For this, an injection of 130 µL of buffer D (6.6 mM Tris-HCl (pH 7.9 at 25°C), 11.2 mM KCl, 1.8 mM NaCl, 0.27 mM MgCl_2_), containing 10 nM SG and no additional SPARDA-gRNA/tDNA complex, was made. In this step, the incubation time for WT XauSPARDA and XauD41A was 15 min, whereas for the remaining XauSPARDA mutants, it was 50 min, and for homologous SPARDAs, this duration was 25 min. After incubation was complete, image series were acquired by flowing the same loaded buffer through the channel (at a rate of 1 mL/min) to stretch any remaining DNA fragments along the surface. In λ DNA cleavage experiments with WT TmuSPARDA, buffer D had an 11-fold higher MgCl_2_ concentration (3 mM), while the concentration of SG remained the same (10 nM). For WT EaeSPARDA, buffer E (28 mM Tris-HCl (pH 7.9 at 25°C), 56 mM KCl, 1.8 mM NaCl, 3 mM MgCl_2_) supplemented with 10 nM SG was used to perform λ DNA cleavage measurements. All λ DNA compaction and cleavage experiments that involved SPARDA proteins assembled in complex with gRNA/lctDNA (in the case of TmuSPARDA, gRNA/nctDNA) were carried out using the same conditions established for tDNA-including experiments.

Colocalization experiments were conducted with WT XauSPARDA. Non-labelled tDNA was mixed with tDNA-ATTO647N at a 51:1 molar ratio. To assemble the complex, 2 µM protein was mixed with 1.5 µM gRNA (final concentrations in this starting mixture are indicated) in buffer C. After incubating the sample at 37°C for 5 min, the prepared mixture of tDNAs was added (as 1/7 volume of the starting mixture) to the pre-warmed initial mixture, resulting in the final concentrations of 428 nM non-labeled tDNA and 8.4 nM tDNA-ATTO647N, and incubation at 37°C for 5 min was performed again. The assembled fluorescently labeled WT XauSPARDA-gRNA/tDNA complex was diluted in 10 nM SG-containing buffer B to the final concentration of 67 nM, 100 µL of such solution loaded into the channel, incubated for 30 min, and then dual-color imaging was performed. The remaining fragments of cleaved λ DNA were visualized 25 min post-injection of 100 µL of buffer D supplemented only with 10 nM SG. In both cases, while acquiring the image series, flow of a corresponding buffer was run through the channel (at the rate of 1 mL/min) to stretch the affected λ DNA molecules along the surface.

Experiments that involved real-time acquisition of λ DNA compaction’s process undergoing continuous buffer-flow conditions were performed using ATTO647N-labelled WT XauSPARDA-gRNA/tDNA complex. It was assembled in the same way as described above, except that the molar ratio of non-labeled tDNA and tDNA-ATTO647N was 89 to 1, and the final concentration of such tDNAs in the complex’s assembly mix was 463 nM and 5.2 nM, respectively. For immobilization of Bsp120I-digested λ DNA molecules in these experiments, the surface of the PEGylated glass coverslip was coated with streptavidin (final concentration of 50 nM in buffer A) instead of traptavidin, while all the other preparation steps remained identical to the original ones. When the channel was filled with 150 µL of 72 nM activated WT XauSPARDA diluted in 10 nM SG-containing buffer B, the peristaltic pump tubing’s outlet was immediately put into the inlet’s reservoir, thus forming a closed loop for the flow of the buffer. Fluorescence movies of 910 frames were recorded at 3.3 Hz with single-end tethered λ DNA molecules being constantly stretched by an uninterrupted buffer flow (running at the rate of 0.5 mL/min).

#### Analysis of Single-Molecule Fluorescence Microscopy Data

The acquired image series were processed using Fiji 2.16.0 software. The selected frames, which had the buffer flow necessary for stretching single end-immobilized λ DNA, launched, were averaged, and then the length of visualized λ DNA molecules was measured. At first, automated detection and marking of individual λ DNA molecules were performed in the averaged images by employing the DNA Finder tool present in the Mars plugin package dedicated to Fiji. After generating a list of regions of interest (ROIs) that were added to it for every DNA molecule identified, this list was then manually verified, discarding any included ROIs that were not corresponding to the exact location or matching the visible contour of single λ DNA molecules present in the image. Finally, any remaining undetected but acceptable DNA molecules were marked by hand, adding their respective ROIs to the final list, which was used to measure the length of such DNAs by running a custom home-written macro for Fiji. DNA molecules that were bound to the surface by their two ends overlapped with each other or appeared in any form of aggregates were not included in this analysis. The extracted length values were then transferred to the environment of Igor Pro 9.0.5.1 software, and most of the further data analysis, including the generation of the graphs, was done there.

Separate sets of measured flow-stretched λ DNA lengths collected from multiple surface positions over a single experiment were pooled together into two distinct groups – all DNA lengths obtained before injecting activated SPARDA into the flowcell’s channel and the lengths of DNA observed after adding and incubating the assembled SPARDA complex. These merged data sets were then used to generate two respective histograms showing the distribution of λ DNA molecules’ length before and after the resultant effect of SPARDA. The apparent peaks in such histograms were then fitted with a single Gaussian function to get the peak center position that represents an average length of DNA per the aforementioned condition. The extracted value of mean λ DNA fragment length before injection of SPARDA was normalized to 1, and the obtained correction factor was used to adjust all the measured DNA lengths for that single specific experiment only. Scatter dot plots of individual λ DNA molecules’ lengths were generated out of this normalized data. Index N, provided next to these graphs and below TIRF microscopy images, indicates the overall number of such data analysis-included λ DNA molecules collected from at least three different surface positions over one or, in some cases, several independent experiments.

To calculate the percentage of two distinct immobilized λ DNA fragment length states – compacted and non-compacted – present on the surface before and after adding the assembled SPARDA-gRNA/tDNA complex, an arbitrary criterion of DNA length for classifying λ DNA molecules into such two groups was introduced. The threshold value of this parameter was set manually for every different data set obtained from a separate experiment by visually selecting the largest λ DNA molecule that still highly resembles activated SPARDA-compacted DNA in one of the raw images, which were acquired after treating λ DNA with SPARDA complex and measuring its length in Fiji (as described above). All λ DNA molecules with lengths lower or equal to this established length threshold value were considered as compacted DNA, whereas the remaining ones were assigned to the category of non-compacted DNA fragments. When lctDNA or nctDNA was used instead of tDNA to assemble the SPARDA complex, the criterion for sorting λ DNA fragment length states in such data sets was kept identical to the one determined from tDNA-involving experiments with the same SPARDA protein. Having this criterion defined for every individual experiment, the quantity of compacted and non-compacted λ DNA molecules present on the surface before and after the effect of SPARDA was counted in the corresponding data sets generated by combining the data from every surface position imaged over that respective stage of the experiment. These extracted quantities were then used to calculate the percentage of compacted and non-compacted λ DNA out of the overall number of λ DNA molecules measured in all surface positions imaged per a specific condition tested. The reported percentage value is a mean of at least two independent experiments.

Quantification of SPARDA-characteristic λ DNA fragments’ cleavage activity was first performed in Fiji. Running buffer flow-lacking frames of the recorded fluorescence movies were averaged. One averaged image with the remaining immobilized λ DNA molecules visualized post-incubation of the assembled SPARDA-gRNA/(lc)tDNA complex in the flowcell’s channel filled with Ca^2+^ ions-containing buffer was selected as a reference image for the subsequent analysis of data collected during that one specific experiment. After fluorescence emission intensity-based thresholding of this image, obtained various-sized spots were counted using an in-built Analyze Particles feature. For both procedures, the parameters were chosen arbitrarily, and they were kept the same when processing all the other averaged images of that particular experiment only. The rest of the data analysis was then continued in Igor Pro 9. The average number of detected λ DNA molecules per field of view was calculated before and after SPARDA-catalyzed hydrolysis of λ DNA. The mean obtained for the ‘before cleavage’ condition (i.e. after incubation with the assembled SPARDA-gRNA/lctDNA complex in the presence of Ca^2+^ ions) was normalized to 1, and so the other mean calculated for the ‘after cleavage’ condition (i.e. after incubation with only SG and Mg^2+^-containing buffer) was adjusted accordingly. The latter was then used to generate bar charts showing the average percentage of the remaining non-cleaved λ DNA molecules, where dots indicate individual values determined for every separate surface position imaged. Here, index N reports the overall number of such data analysis-included λ DNA molecules collected from at least two different surface positions over a single experiment. For all XauSPARDA variants in which cleavage activity was studied at the SM level, N is >500. In the case of homologous SPARDAs, N is >300, except for NbaSPARDA-gRNA/tDNA (N = 68). Not all surface positions imaged before and after Mg^2+^ ions’ addition-initiated cleavage of SPARDA-treated immobilized λ DNA were the same.

### QUANTIFICATION AND STATISTICAL ANALYSIS

Statistical details for each experiment are found in the figure legend and the accompanying method details.

#### RNA and DNA sequences correlation analysis

The correlation between total RNA and SPARDA-associated small RNA sequence abundance and total RNA and DNA cleavage site sequence abundance was tested for each comparison by calculating the Spearman correlation coefficient.

## SUPPLEMENTARY INFORMATION

### Supplementary figures

**Figure S1.**
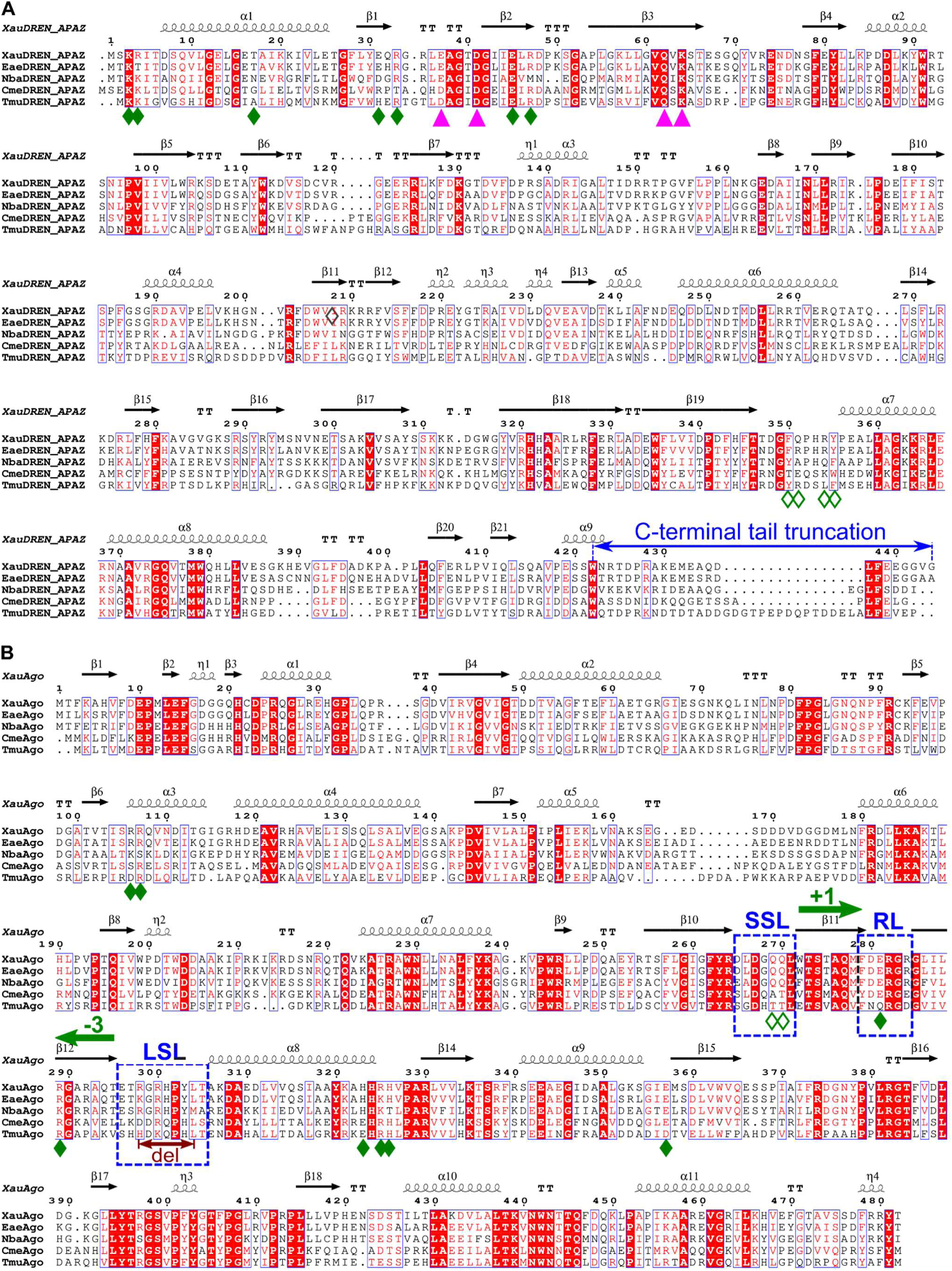
Multiple sequence alignments of experimentally characterized SPARDA systems. (**A**) Alignment of DREN-APAZ sequences. Residues important for catalytic activity and for the formation of either Apo form or active filament are indicated below sequence alignment. Active site residues within the PD-(D/E)XK domain are indicated with pink triangles. Residues involved in tetramerization of an Apo form and an active filament are labeled with empty and filled green diamonds, respectively. The truncated region of the C-terminal tail is indicated above the alignment. (**B**) Sequence alignment of MID-PIWI proteins. Residues involved in tetramerization of an Apo form and an active filament are labeled with empty and filled green diamonds, respectively. The direction and the extent of register shifts within “beta-relay” upon Xau SPARDA activation are indicated correspondingly with green arrows and numbers. The positions of small sensor loop (SLS), large sensor loop (LSL) and response loop (RL) are indicated with blue dashed rectangles. The deletion (del) within LSL is indicated below the alignment. Xau, *Xanthobacter autotrophicus Py2*; Eae, *Enhydrobacter aerosaccus*; Nba, *Novosphingopyxis baekryungensis*; Cme, *Cupriavidus metallidurans*; Tmu, *Thermocrispum municipale*. The figure was prepared using ESPript 3 (PMID: 24753421). Related to Figure 1, 2 and 4

**Figure S2.**
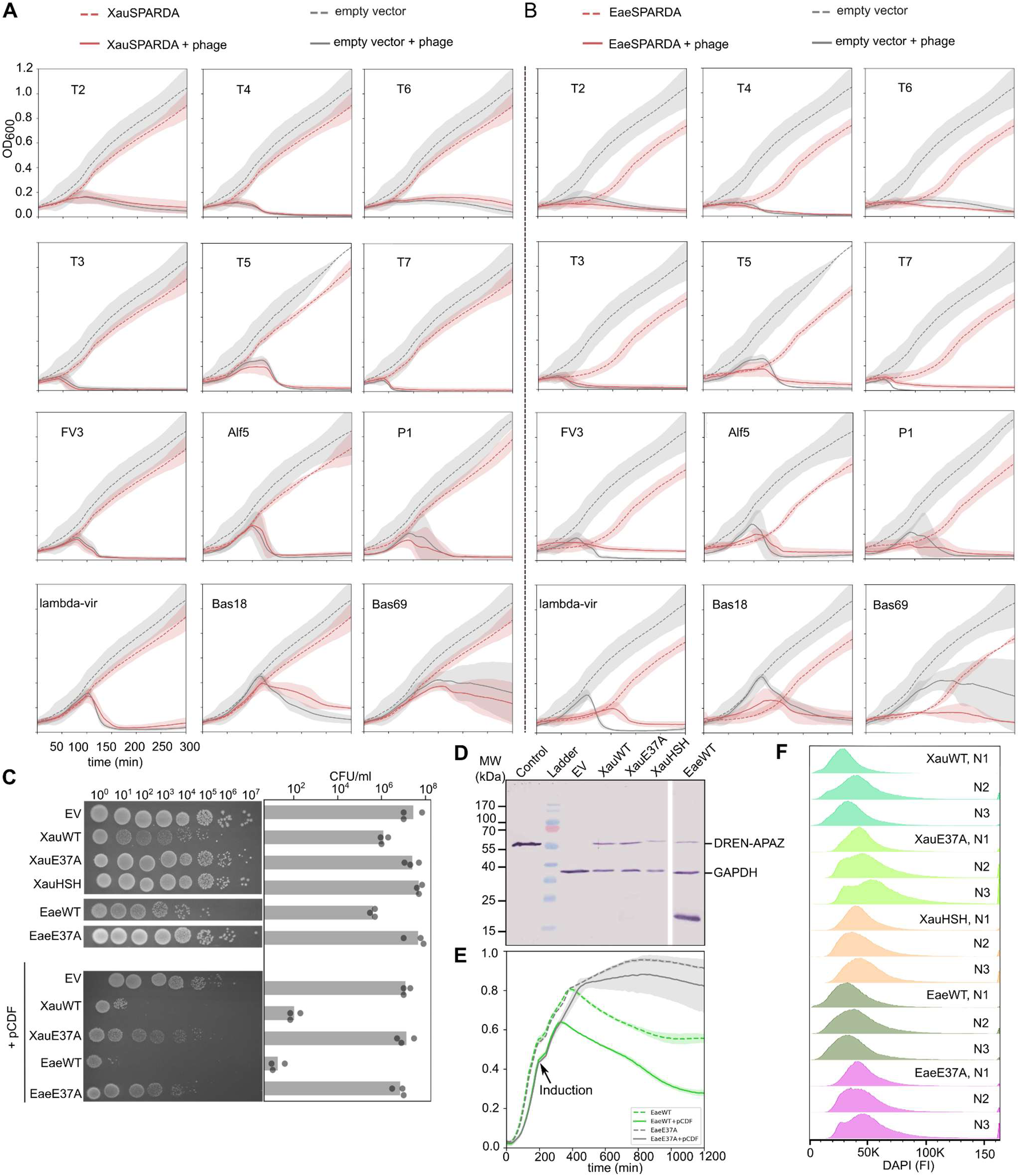
SPARDA is an anti-plasmid defense system. *E. coli* cultures expressing XauSPARDA **(A)** or EaeSPARDA **(B)** or harboring an empty vector were grown in the presence of various bacteriophages (bacteriophage type indicated within each graph), and the OD600 nm was monitored over time. The average of three biological replicates is shown; shadings indicate standard deviations. **(C)** Cytotoxic effect of SPARDA system on *E. coli* cells expressing XauSPARDA (XauWT) or EaeSPARDA (EaeWT) system, or harboring an empty vector (EV) in the absence or the presence of second plasmid pCDF (indicated +pCDF). Left: serial dilutions from 1 to 10^7^ on agar plates, right: colony forming units (CFU) were counted, and the average of three biological replicates is shown in bars. **(D)** Anti-His Tag Western blot analysis of *E. coli* BL21-AI lysates expressing Xau and EaeSPARDA for 4 hours or harboring empty vector (EV). Control - purified XauSPARDA protein, where the visible part is DREN-APAZ (indicated on the right); GAPDH - loading control. **(E)** Growth curves of *E. coli* cultures expressing EaeSPARDA with and without pCDF plasmid. Data are shown as mean ± SD across three biological replicates. **(F)** DAPI signal of *E. coli* cells expressing SPARDA systems in the presence of pCDF plasmid. The DAPI signal intensity (FI) of stained cells was measured by flow cytometry. The three biological replicates (N1-3) are shown in profiles of DAPI mode. Related to Figure 1.

**Figure S3.**
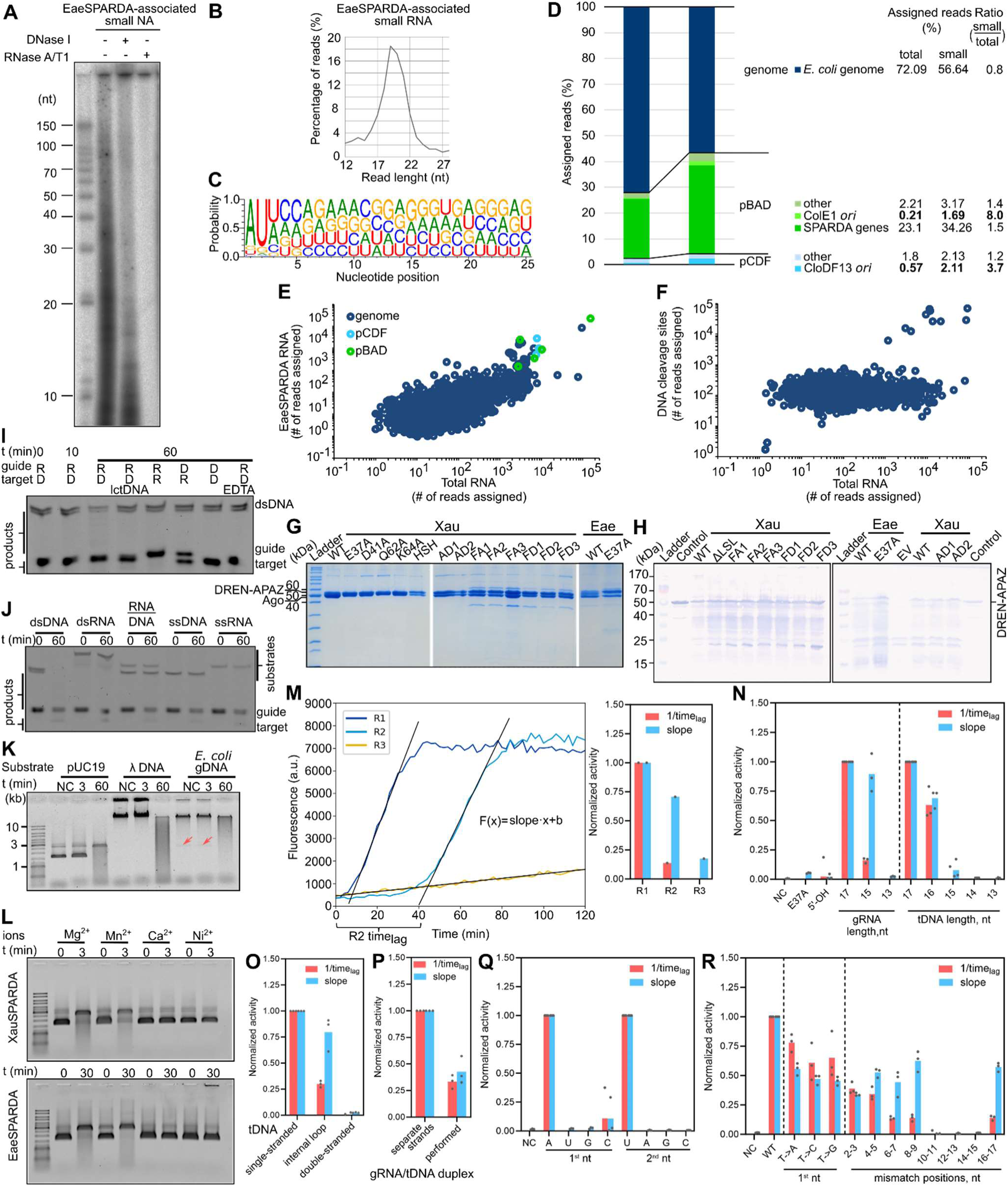
Biochemical characterization of Xau and EaeSPARDA. **(A)** Identification of EaeSPARDA-bound nucleic acids. Nucleic acids copurified with EaeSPARDA were radiolabeled, treated with DNase I or RNase A/T1, and resolved on a denaturing polyacrylamide gel. **(B)** Length distribution of small RNA copurified with EaeSPARDA as determined by sequencing. **(C)** Nucleotide bias of small RNAs copurified with EaeSPARDA. **(D)** Percentages of total RNA (*E. coli* expressing EaeSPARDA) and small RNA (copurified with EaeSPARDA) reads that align to specific genomic or plasmid sequences. **(E)** Correlation between EaeSPARDA-associated small RNA and *E. coli*-extracted total RNA sequences. **(F)** XauSPARDA-associated DNA cleavage site sequences and *E. coli*-extracted total RNA sequences do not correlate. **(G)** SDS-PAGE of purified Xau and EaeSPARDA proteins used in this study. **(H)** Anti-Strep-tag Western blot analysis of *E. coli* BL21-AI lysates expressing Xau and EaeSPARDA for 4 hours or harboring empty vector (EV). Control - purified XauSPARDA protein, where the visible part is DREN-APAZ (indicated on the left). **(I)** Analysis of the specificity of EaeSPARDA for gRNA and tDNA with a dsDNA substrate. **(J)** Collateral cleavage activity of EaeSPARDA with various short NA substrates. **(K)** Collateral cleavage activity of EaeSPARDA with various long dsDNA substrates. NC (negative control) - reaction without protein. The red arrow indicates the pBAD plasmid. **(L)** Collateral cleavage activity of Xau and EaeSPARDA in the presence of various divalent metal ions using pUC19 plasmid DNA as a substrate. **(M)** Calculation of the SPARDA collateral cleavage activity parameters (slope and 1/time_lag_) by measuring the fluorescent signal change of the substrate during its hydrolysis reaction. (Left) The equation of the line (F(x)= slope*x+b) was fitted to the experimental points where the largest signal change was observed. The slope of the resulting line activity indicates the cleavage rate (slope). The X-intercept of the resulting line corresponds to the lag time (time_lag_), and the parameter 1/time_lag_ reflects how fast the cleavage begins after the reaction. Calculated values were plotted as bar charts (right). **(N)** Cleavage activity of EaeSPARDA with gRNAs and tDNAs of various lengths. 5′-OH depicts dephosphorylated gRNA of 17nt length. NC (negative control) - reaction without protein. E37A is active center mutant EaeE37A. **(O)** Collateral cleavage activity of XauSPARDA with various tDNA forms. **(P)** Effect of different gRNA-tDNA duplex forms on cleavage activity of XauSPARDA. **(Q)** Effects of substitutions of the first or second 5’-nucleotide in gRNA on the cleavage activity of EaeSPARDA**. (R)** Effect of single or double mismatches within tDNA on EaeSPARDA activation. Related to Figures 1 and 2.

**Figure S4.**
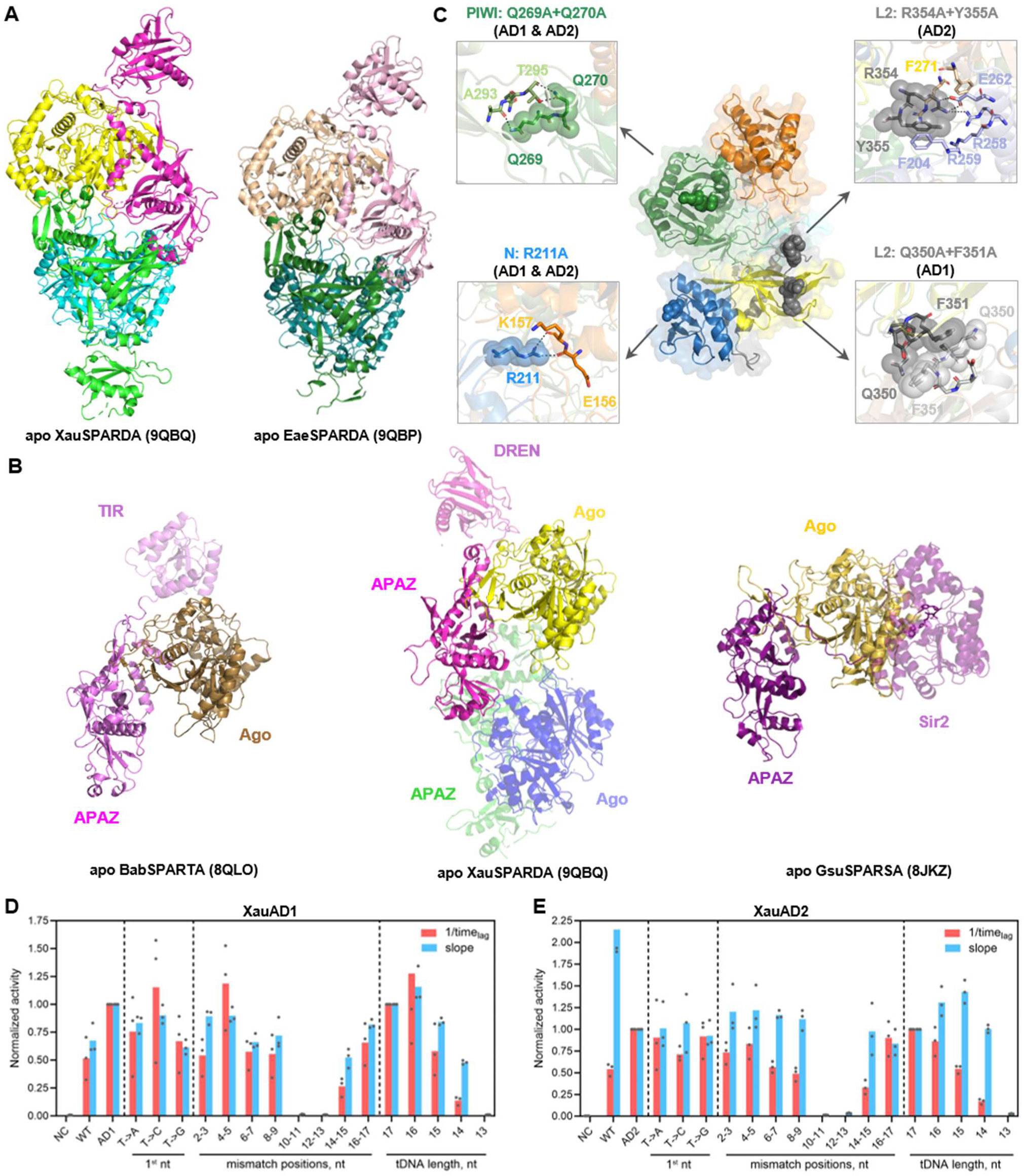
Apo Xau and EaeSPARDA are dimers. **(A)** Crystal structures of apo Xau and EaeSPARDA dimers. Subunits are in different colors. **(B)** Comparison of apo XauSPARDA (center) with apo BabSPARTA (left) and apo GsuSPARSA (right). Binary monomers are overlaid by secondary structure elements of Ago proteins. The other monomer of XauSPARDA (blue-green) is transparent. Effector domains are also transparent. Ago subunits are colored in yellow tints, and APAZ proteins are colored in magenta tints. **(C)** Mutations of the apo dimerization surface of XauSPARDA. Mutated residues are shown as spheres, and close views are shown on side panels. Domains are colored as in Figure 1A. Cleavage activity of XauSPARDA apo dimerization mutants XauAD1 **(D)** and XauAD2 **(E)**. WT indicates WT XauSPARDA. Related to Figure 2.

**Figure S5.**
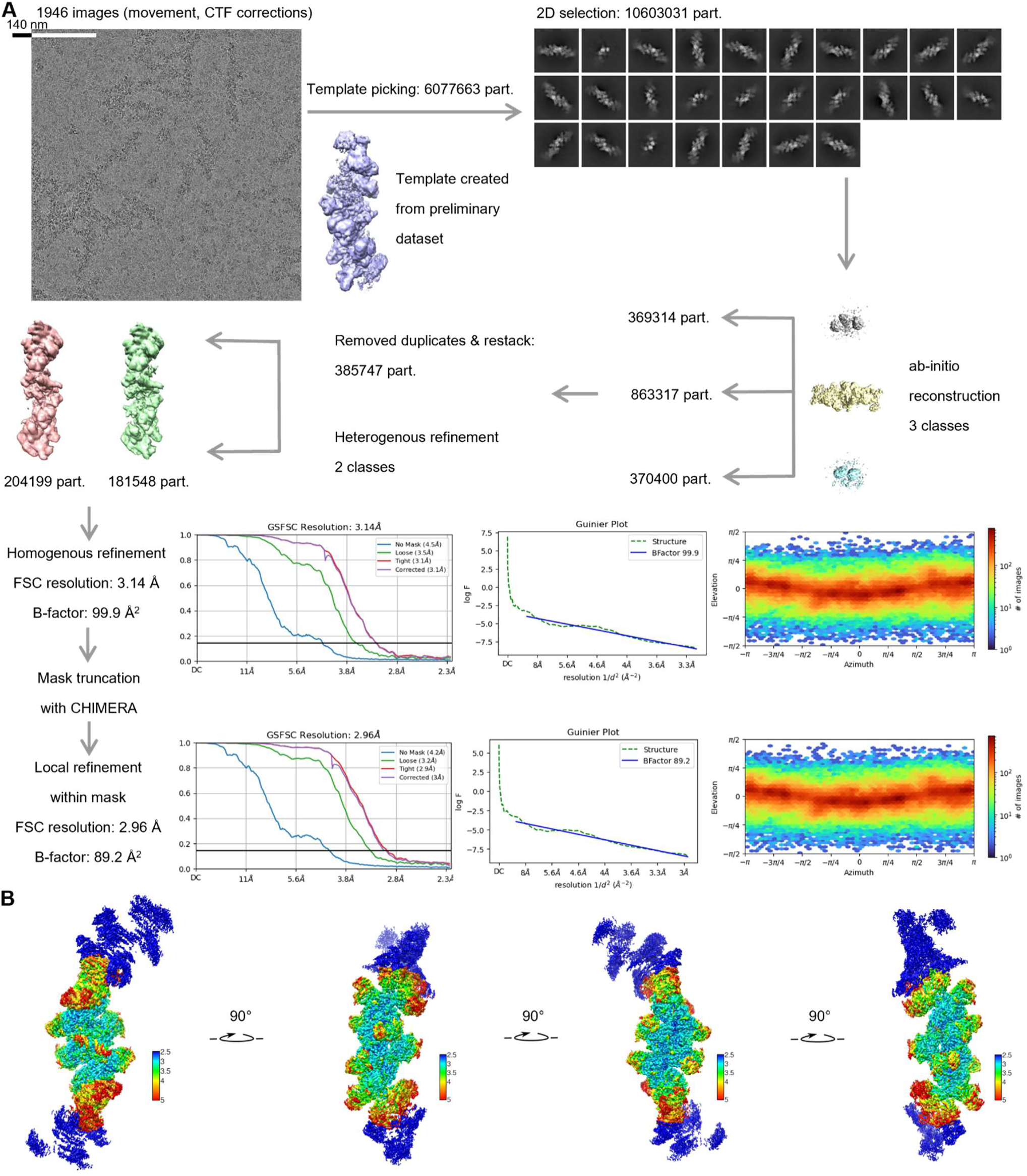
Cryo-EM structure of the activated XauSPARDA filament (PDB ID: 9QCC and EMD-53008). **(A)** Cryo-EM structure solution workflow. **(B)** Local resolution map of the volume refined within the mask. Related to Figure 3.

**Figure S6.**
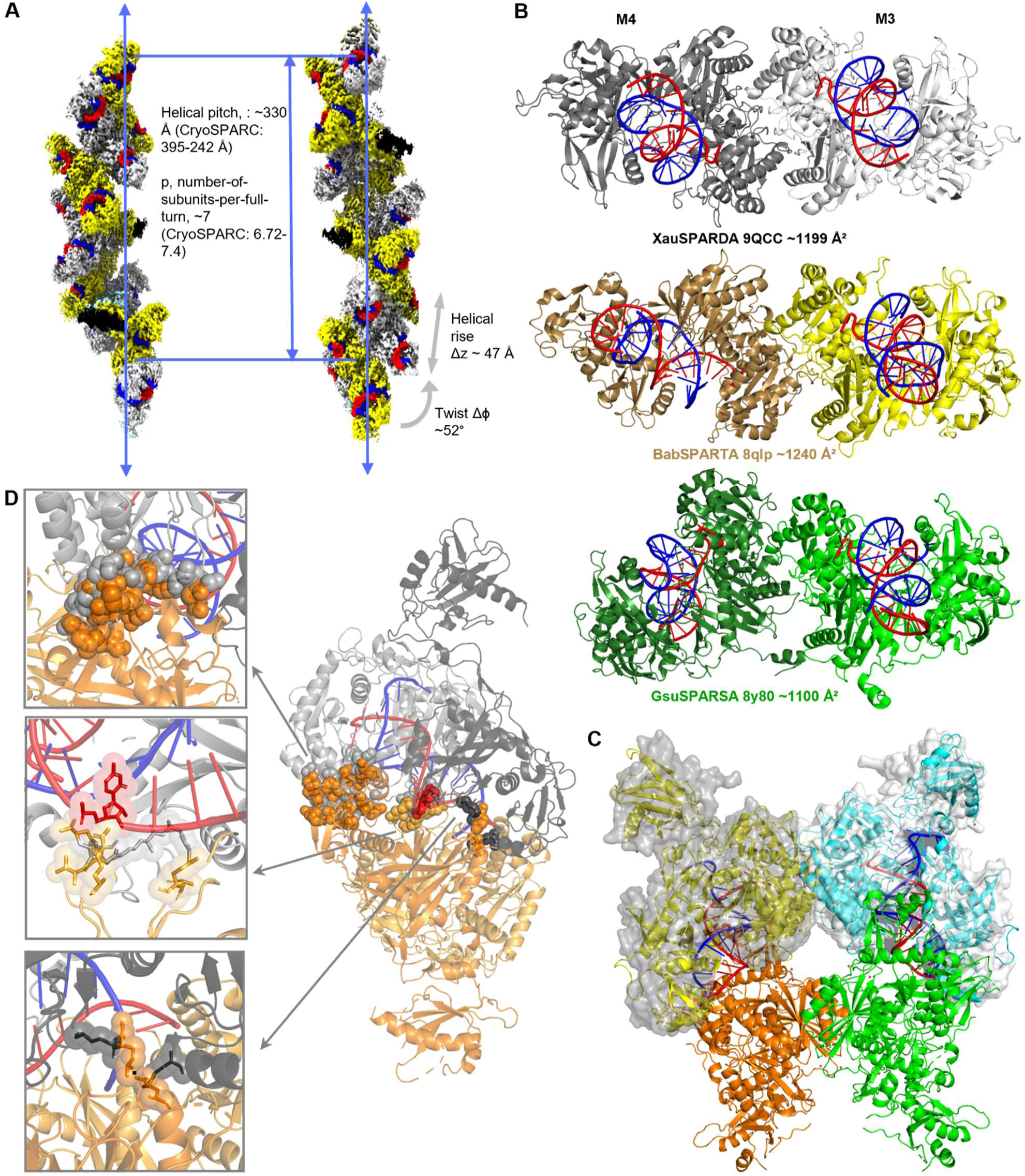
Activated XauSPARDA. **(A)** Helical structure of XauSPARDA filament. Elongated filament constructed from 3 translated models. Approximate parameters of the helix are estimated by direct measurements of the model and calculated using CryoSPARC. **(B)** Comparison of dimerization modes of activated oligomeric short pAgo. Dimers of BabSPARTA (yellow, PDB ID: 8qlp) and GsuSPARSA (green, PDB ID: 8y80) are superimposed by one of the monomers with M3 on the central step of the activated XauSPARDA complex (this study, PDB ID: 9QCC). The dimerization surfaces were calculated by VoroContacts. **(C)** Formation of the activated XauSPARDA complex requires disruption of the apo dimer. Two apo dimers (yellow-orange and cyan-green) are superimposed (by residues Ago:150-450) on dimers of the central step of XauSPARDA (shown as transparent grey surfaces). Ago subunits of apo complexes clash (orange and green) under the step. **(D)** Guide/target binding disrupts XauSPARDA apo dimer. “Chimeric” complex is constructed from the guide/target-bound XauSPARDA monomer, M3 of central step (Ago: grey, DREN-APAZ: dark grey) and apo-monomer (DREN-APAZ: light orange, Ago: orange). Three main clashing surfaces are shown as spheres and enlarged in inserts. The guide RNA strand is red, and the DNA target is blue. Related to Figure 4.

**Figure S7.**
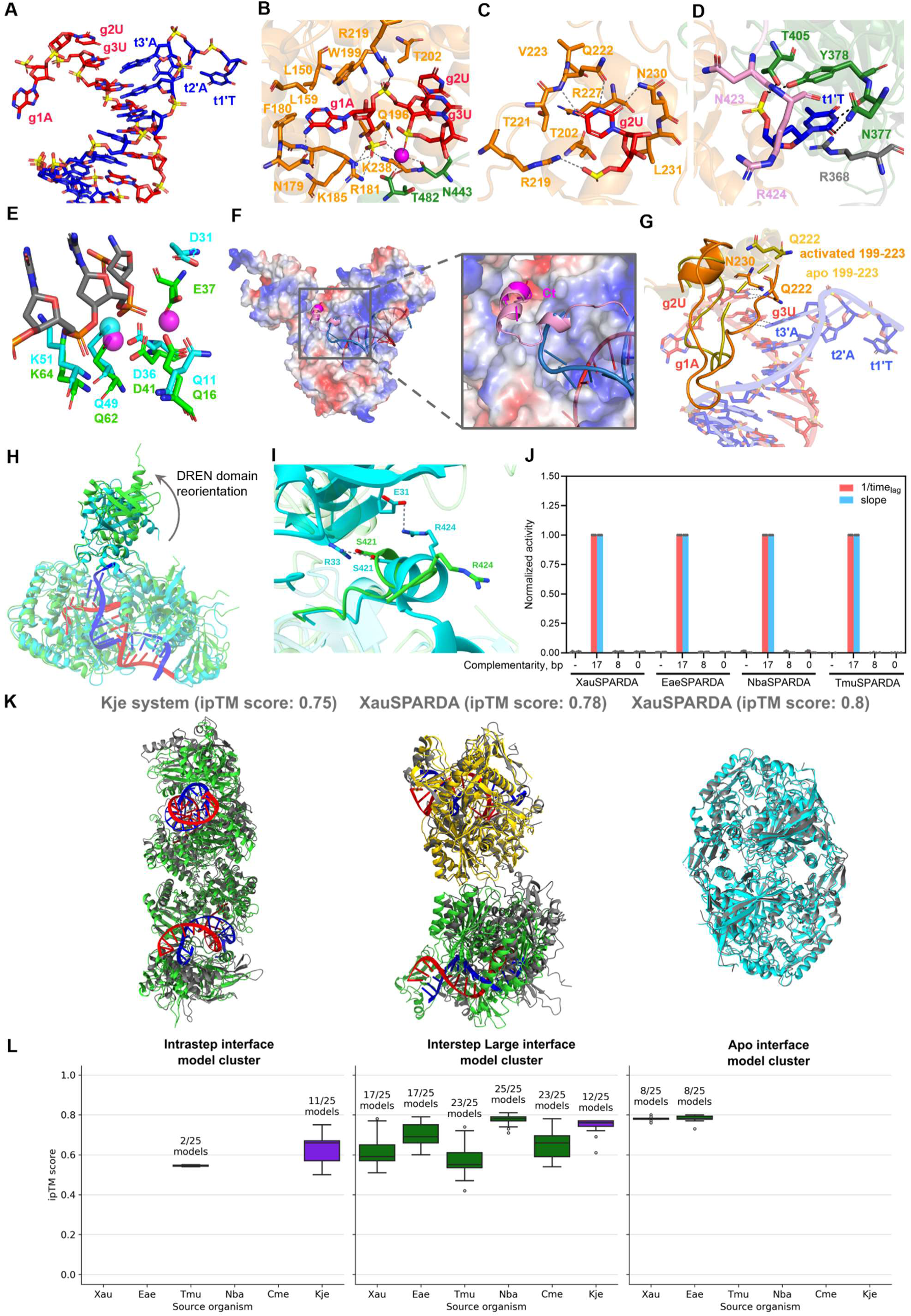
Activated XauSPARDA and homologous systems. **(A)** Guide/target heteroduplex bound in XauSPARDA. gRNA is colored red, tDNA - blue. **(B)** Binding of gRNA 5’-end. Protein residues are colored according to domains. Ca^2+^ is shown as a magenta sphere. H-bonds are shown by dashed lines. **(C)** Contacts of g2U base. **(D)** Binding of t1’T nucleotide. **(E)** Active center residues of the active DREN domain (green, A-DREN) are superimposed with homing endonuclease I-Ssp6803I (cyan, PDB ID: 2ost). dsDNA substrate is grey. **(F)** Ct from apo XauSPARDA would clash with the first base of tDNA. Apo XauSPARDA Ct is colored pink as in the domain scheme and is shown along with the electrostatic surface of activated XauSPARDA. A few residues of activated XauSPARDA Ct (414-424) are colored magenta. **(G)** Loop Ago:199-223 is disordered in apo XauSPARDA (yellow) upon gRNA/tDNA binding (orange) interacts with t3’-t7’ backbone. Q222 at the beginning of the loop disrupts the first two bp of gRNA/tDNA heteroduplex and keeps g2U base within the duplex, replacing the t2’ base. **(H)** Overlay of XauSPARDA monomers from Apo form (turquoise) and filament (green). The DREN domain, whose reorganization is the most pronounced, is highlighted. **(I)** Dynamics of interactions between the DREN domain and the Ct tail. XauSPARDA’s Apo form monomer is colored turquoise, while a monomer from the filament is colored green. **(J)** Effect of gRNA and tDNA complementarity on different SPARDA proteins. The length of the gRNA and tDNA was 17 nt. Numbers indicate the complementarity (bp) of gRNA and tDNA from the 5’ end of the RNA guide. In the case of the “0” reaction, a 17 nt poly(dT) oligonucleotide was used as tDNA. The “−” reaction represents a control lacking tDNA. **(K)** Structural overlay of the best-scored representatives (best-scored XauSPARDA model or best-scored model of another system if there are no XauSPARDA models) from the TDM clusters onto the corresponding interfaces of XauSPARDA. The reference XauSPARDA filament central step monomer is colored yellow, the neighboring step monomers are colored green, the Apo form monomers are colored turquoise, and the AlphaFold models are colored grey. **(L)** A boxplot of TDM populations of proteins from different sources in the model clusters. The labels above the boxes indicate the number of models (out of 25 models produced in each case) present in each cluster. The box colors indicate the phylogeny of different systems. Green represents the SPARDA systems, while purple represents the SPARMA system from *K. jejudonensis*. Related to Figures 4, 6, and 7.

## REFERENCES

1. Ozata, D.M., Gainetdinov, I., Zoch, A., O’Carroll, D., and Zamore, P.D. (2019). PIWI-interacting RNAs: small RNAs with big functions. Nat Rev Genet 20, 89–108. 10.1038/s41576-018-0073-3.

2. Park, M.S., Sim, G., Kehling, A.C., and Nakanishi, K. (2020). Human Argonaute2 and Argonaute3 are catalytically activated by different lengths of guide RNA. Proc. Natl. Acad. Sci. U.S.A. 117, 28576–28578. 10.1073/pnas.2015026117.

3. Koopal, B., Potocnik, A., Mutte, S.K., Aparicio-Maldonado, C., Lindhoud, S., Vervoort, J.J.M., Brouns, S.J.J., and Swarts, D.C. (2022). Short prokaryotic Argonaute systems trigger cell death upon detection of invading DNA. Cell 185, 1471–1486.e19. 10.1016/j.cell.2022.03.012.

4. Swarts, D.C., Jore, M.M., Westra, E.R., Zhu, Y., Janssen, J.H., Snijders, A.P., Wang, Y., Patel, D.J., Berenguer, J., Brouns, S.J.J., et al. (2014). DNA-guided DNA interference by a prokaryotic Argonaute. Nature 507, 258–261. 10.1038/nature12971.

5. Olovnikov, I., Chan, K., Sachidanandam, R., Newman, D.K., and Aravin, A.A. (2013). Bacterial Argonaute Samples the Transcriptome to Identify Foreign DNA. Molecular Cell 51, 594–605. 10.1016/j.molcel.2013.08.014.

6. Wang, Y., Sheng, G., Juranek, S., Tuschl, T., and Patel, D.J. (2008). Structure of the guide-strand-containing argonaute silencing complex. Nature 456, 209–213. 10.1038/nature07315.

7. Wang, Y., Juranek, S., Li, H., Sheng, G., Wardle, G.S., Tuschl, T., and Patel, D.J. (2009). Nucleation, propagation and cleavage of target RNAs in Ago silencing complexes. Nature 461, 754–761. 10.1038/nature08434.

8. Wang, Y., Juranek, S., Li, H., Sheng, G., Tuschl, T., and Patel, D.J. (2008). Structure of an argonaute silencing complex with a seed-containing guide DNA and target RNA duplex. Nature 456, 921–926. 10.1038/nature07666.

9. Nakanishi, K., Weinberg, D.E., Bartel, D.P., and Patel, D.J. (2012). Structure of yeast Argonaute with guide RNA. Nature 486, 368–374. 10.1038/nature11211.

10. Schirle, N.T., and MacRae, I.J. (2012). The Crystal Structure of Human Argonaute2. Science 336, 1037– 1040. 10.1126/science.1221551.

11. Wilson, R.C., and Doudna, J.A. (2013). Molecular Mechanisms of RNA Interference. Annu. Rev. Biophys. 42, 217–239. 10.1146/annurev-biophys-083012-130404.

12. Vagin, V.V., Sigova, A., Li, C., Seitz, H., Gvozdev, V., and Zamore, P.D. (2006). A Distinct Small RNA Pathway Silences Selfish Genetic Elements in the Germline. Science 313, 320–324. 10.1126/science.1129333.

13. Ryazansky, S., Kulbachinskiy, A., and Aravin, A.A. (2018). The Expanded Universe of Prokaryotic Argonaute Proteins. mBio 9, e01935–18. 10.1128/mBio.01935-18.

14. Prostova, M., Kanevskaya, A., Panteleev, V., Lisitskaya, L., Perfilova Tugaeva, K.V., Sluchanko, N.N., Esyunina, D., and Kulbachinskiy, A. (2024). DNA-targeting short Argonautes complex with effector proteins for collateral nuclease activity and bacterial population immunity. Nat Microbiol 9, 1368–1381. 10.1038/s41564-024-01654-5.

15. Zaremba, M., Dakineviciene, D., Golovinas, E., Zagorskaitė, E., Stankunas, E., Lopatina, A., Sorek, R., Manakova, E., Ruksenaite, A., Silanskas, A., et al. (2022). Short prokaryotic Argonautes provide defence against incoming mobile genetic elements through NAD+ depletion. Nat Microbiol 7, 1857–1869. 10.1038/s41564-022-01239-0.

16. Kuzmenko, A., Oguienko, A., Esyunina, D., Yudin, D., Petrova, M., Kudinova, A., Maslova, O., Ninova, M., Ryazansky, S., Leach, D., et al. (2020). DNA targeting and interference by a bacterial Argonaute nuclease. Nature 587, 632–637. 10.1038/s41586-020-2605-1.

17. Jolly, S.M., Gainetdinov, I., Jouravleva, K., Zhang, H., Strittmatter, L., Bailey, S.M., Hendricks, G.M., Dhabaria, A., Ueberheide, B., and Zamore, P.D. (2020). Thermus thermophilus Argonaute Functions in the Completion of DNA Replication. Cell 182, 1545–1559.e18. 10.1016/j.cell.2020.07.036.

18. Fu, L., Xie, C., Jin, Z., Tu, Z., Han, L., Jin, M., Xiang, Y., and Zhang, A. (2019). The prokaryotic Argonaute proteins enhance homology sequence-directed recombination in bacteria. Nucleic Acids Research 47, 3568– 3579. 10.1093/nar/gkz040.

19. Lee, K.Z., Mechikoff, M.A., Kikla, A., Liu, A., Pandolfi, P., Fitzgerald, K., Gimble, F.S., and Solomon, K.V. (2021). NgAgo possesses guided DNA nicking activity. Nucleic Acids Research 49, 9926–9937. 10.1093/nar/gkab757.

20. Song, X., Lei, S., Liu, S., Liu, Y., Fu, P., Zeng, Z., Yang, K., Chen, Y., Li, M., She, Q., et al. (2023). Catalytically inactive long prokaryotic Argonaute systems employ distinct effectors to confer immunity via abortive infection. Nat Commun 14, 6970. 10.1038/s41467-023-42793-3.

21. Manakova, E., Golovinas, E., Pocevičiūtė, R., Sasnauskas, G., Silanskas, A., Rutkauskas, D., Jankunec, M., Zagorskaitė, E., Jurgelaitis, E., Grybauskas, A., et al. (2024). The missing part: the *Archaeoglobus fulgidus* Argonaute forms a functional heterodimer with an N-L1-L2 domain protein. Nucleic Acids Research 52, 2530–2545. 10.1093/nar/gkad1241.

22. Makarova, K.S., Wolf, Y.I., Van Der Oost, J., and Koonin, E.V. (2009). Prokaryotic homologs of Argonaute proteins are predicted to function as key components of a novel system of defense against mobile genetic elements. Biol Direct 4, 29. 10.1186/1745-6150-4-29.

23. Finocchio, G., Koopal, B., Potocnik, A., Heijstek, C., Westphal, A.H., Jinek, M., and Swarts, D.C. (2024). Target DNA-dependent activation mechanism of the prokaryotic immune system SPARTA. Nucleic Acids Research 52, 2012–2029. 10.1093/nar/gkad1248.

24. Shen, Z., Yang, X.-Y., Xia, S., Huang, W., Taylor, D.J., Nakanishi, K., and Fu, T.-M. (2023). Oligomerization-mediated activation of a short prokaryotic Argonaute. Nature 621, 154–161. 10.1038/s41586-023-06456-z.

25. Wang, X., Li, X., Yu, G., Zhang, L., Zhang, C., Wang, Y., Liao, F., Wen, Y., Yin, H., Liu, X., et al. (2023). Structural insights into mechanisms of Argonaute protein-associated NADase activation in bacterial immunity. Cell Res 33, 699–711. 10.1038/s41422-023-00839-7.

26. Ni, D., Lu, X., Stahlberg, H., and Ekundayo, B. (2023). Activation mechanism of a short argonaute-TIR prokaryotic immune system. Sci. Adv. 9, eadh9002. 10.1126/sciadv.adh9002.

27. Guo, M., Zhu, Y., Lin, Z., Yang, D., Zhang, A., Guo, C., and Huang, Z. (2023). Cryo-EM structure of the ssDNA-activated SPARTA complex. Cell Res 33, 731–734. 10.1038/s41422-023-00850-y.

28. Kottur, J., Malik, R., and Aggarwal, A.K. (2024). Nucleic acid mediated activation of a short prokaryotic Argonaute immune system. Nat Commun 15, 4852. 10.1038/s41467-024-49271-4.

29. Gao, X., Shang, K., Zhu, K., Wang, L., Mu, Z., Fu, X., Yu, X., Qin, B., Zhu, H., Ding, W., et al. (2024). Nucleic-acid-triggered NADase activation of a short prokaryotic Argonaute. Nature 625, 822–831. 10.1038/s41586-023-06665-6.

30. Zhang, J.-T., Wei, X.-Y., Cui, N., Tian, R., and Jia, N. (2024). Target ssDNA activates the NADase activity of prokaryotic SPARTA immune system. Nat Chem Biol 20, 503–511. 10.1038/s41589-023-01479-z.

31. Guo, L., Huang, P., Li, Z., Shin, Y.-C., Yan, P., Lu, M., Chen, M., and Xiao, Y. (2024). Auto-inhibition and activation of a short Argonaute-associated TIR-APAZ defense system. Nat Chem Biol 20, 512–520. 10.1038/s41589-023-01478-0.

32. Cui, N., Zhang, J.-T., Li, Z., Wei, X.-Y., Wang, J., and Jia, N. (2024). Tetramerization-dependent activation of the Sir2-associated short prokaryotic Argonaute immune system. Nat Commun 15, 8610. 10.1038/s41467-024-52910-5.

33. Zhen, X., Xu, X., Ye, L., Xie, S., Huang, Z., Yang, S., Wang, Y., Li, J., Long, F., and Ouyang, S. (2024). Structural basis of antiphage immunity generated by a prokaryotic Argonaute-associated SPARSA system. Nat Commun 15, 450. 10.1038/s41467-023-44660-7.

34. Sun, D., Zhu, K., Wang, L., Mu, Z., Wu, K., Hua, L., Qin, B., Gao, X., Wang, Y., and Cui, S. (2024). Nucleic acid-induced NADase activation of a short Sir2-associated prokaryotic Argonaute system. Cell Reports 43, 114391. 10.1016/j.celrep.2024.114391.

35. Lu, X., Xiao, J., Wang, L., Zhu, B., and Huang, F. (2024). The nuclease-associated short prokaryotic Argonaute system nonspecifically degrades DNA upon activation by target recognition. Nucleic Acids Research 52, 844–855. 10.1093/nar/gkad1145.

36. Toliusis, P., Grybauskas, A., Sinkunas, T., Karvelis, T., Sasnauskas, G., and Zaremba, M. (2025). Off-target detection of CRISPR-Cas9 nuclease in vitro with CROFT-Seq. bioRxiv. 10.1101/2025.02.17.638614.

37. Tolia, N.H., and Joshua-Tor, L. (2006). Strategies for protein coexpression in Escherichia coli. Nat Methods 3, 55–64. 10.1038/nmeth0106-55.

38. Kim, S.-Y., Jung, Y., and Lim, D. (2020). Argonaute system of Kordia jejudonensis is a heterodimeric nucleic acid-guided nuclease. Biochemical and Biophysical Research Communications 525, 755–758. 10.1016/j.bbrc.2020.02.145.

39. Chao, Y., Li, L., Girodat, D., Förstner, K.U., Said, N., Corcoran, C., Śmiga, M., Papenfort, K., Reinhardt, R., Wieden, H.-J., et al. (2017). In Vivo Cleavage Map Illuminates the Central Role of RNase E in Coding and Non-coding RNA Pathways. Molecular Cell 65, 39–51. 10.1016/j.molcel.2016.11.002.

40. Mackie, G.A. (2013). RNase E: at the interface of bacterial RNA processing and decay. Nat Rev Microbiol 11, 45–57. 10.1038/nrmicro2930.

41. Hoyos, M., Huber, M., Förstner, K.U., and Papenfort, K. (2020). Gene autoregulation by 3’ UTR-derived bacterial small RNAs. eLife 9, e58836. 10.7554/eLife.58836.

42. McDowall, K.J., Lin-Chao, S., and Cohen, S.N. (1994). A+U content rather than a particular nucleotide order determines the specificity of RNase E cleavage. J Biol Chem 269, 10790–10796.

43. Ma, J.-B., Yuan, Y.-R., Meister, G., Pei, Y., Tuschl, T., and Patel, D.J. (2005). Structural basis for 5′-end-specific recognition of guide RNA by the A. fulgidus Piwi protein. Nature 434, 666–670. 10.1038/nature03514.

44. Miyoshi, T., Ito, K., Murakami, R., and Uchiumi, T. (2016). Structural basis for the recognition of guide RNA and target DNA heteroduplex by Argonaute. Nat Commun 7, 11846. 10.1038/ncomms11846.

45. Willkomm, S., Oellig, C.A., Zander, A., Restle, T., Keegan, R., Grohmann, D., and Schneider, S. (2017). Structural and mechanistic insights into an archaeal DNA-guided Argonaute protein. Nat Microbiol 2, 17035. 10.1038/nmicrobiol.2017.35.

46. Zhao, L., Bonocora, R.P., Shub, D.A., and Stoddard, B.L. (2007). The restriction fold turns to the dark side: a bacterial homing endonuclease with a PD-(D/E)-XK motif. EMBO J 26, 2432–2442. 10.1038/sj.emboj.7601672.

47. Dai, Z., Chen, Y., Guan, Z., Chen, X., Tan, K., Yang, K., Yan, X., Liu, Y., Gong, Z., Han, W., et al. (2025). Structural and mechanistic insights into the activation of a short prokaryotic argonaute system from archaeon Sulfolobus islandicus. Nucleic Acids Res 53, gkaf059. 10.1093/nar/gkaf059.

48. Song, J.-J., Smith, S.K., Hannon, G.J., and Joshua-Tor, L. (2004). Crystal Structure of Argonaute and Its Implications for RISC Slicer Activity. Science 305, 1434–1437. 10.1126/science.1102514.

49. Wang, L., Chen, W., Zhang, C., Xie, X., Huang, F., Chen, M., Mao, W., Yu, N., Wei, Q., Ma, L., et al. (2024). Molecular mechanism for target recognition, dimerization, and activation of Pyrococcus furiosus Argonaute. Molecular Cell 84, 675–686.e4. 10.1016/j.molcel.2024.01.004.

50. Sheng, G., Zhao, H., Wang, J., Rao, Y., Tian, W., Swarts, D.C., Van Der Oost, J., Patel, D.J., and Wang, Y. (2014). Structure-based cleavage mechanism of *Thermus thermophilus* Argonaute DNA guide strand-mediated DNA target cleavage. Proc. Natl. Acad. Sci. U.S.A. 111, 652–657. 10.1073/pnas.1321032111.

51. Tao, X., Ding, H., Wu, S., Wang, F., Xu, H., Li, J., Zhai, C., Li, S., Chen, K., Wu, S., et al. (2024). Structural and mechanistic insights into a mesophilic prokaryotic Argonaute. Nucleic Acids Res 52, 11895–11910. 10.1093/nar/gkae820.

52. Doxzen, K.W., and Doudna, J.A. (2017). DNA recognition by an RNA-guided bacterial Argonaute. PLoS ONE 12, e0177097. 10.1371/journal.pone.0177097.

53. Kaya, E., Doxzen, K.W., Knoll, K.R., Wilson, R.C., Strutt, S.C., Kranzusch, P.J., and Doudna, J.A. (2016). A bacterial Argonaute with noncanonical guide RNA specificity. Proc. Natl. Acad. Sci. U.S.A. 113, 4057–4062. 10.1073/pnas.1524385113.

54. Saldaño, T., Escobedo, N., Marchetti, J., Zea, D.J., Mac Donagh, J., Velez Rueda, A.J., Gonik, E., García Melani, A., Novomisky Nechcoff, J., Salas, M.N., et al. (2022). Impact of protein conformational diversity on AlphaFold predictions. Bioinformatics 38, 2742–2748. 10.1093/bioinformatics/btac202.

55. Georjon, H., and Bernheim, A. (2023). The highly diverse antiphage defence systems of bacteria. Nat Rev Microbiol 21, 686–700. 10.1038/s41579-023-00934-x.

56. Rousset, F., and Sorek, R. (2023). The evolutionary success of regulated cell death in bacterial immunity. Current Opinion in Microbiology 74, 102312. 10.1016/j.mib.2023.102312.

57. Vassylyev, D.G., Vassylyeva, M.N., Perederina, A., Tahirov, T.H., and Artsimovitch, I. (2007). Structural basis for transcription elongation by bacterial RNA polymerase. Nature 448, 157–162. 10.1038/nature05932.

58. Masukata, H., Dasgupta, S., and Tomizawa, J. (1987). Transcriptional activation of ColE1 DNA synthesis by displacement of the nontranscribed strand. Cell 51, 1123–1130. 10.1016/0092-8674(87)90598-8.

59. Nakanishi, K. (2024). When Argonaute takes out the ribonuclease sword. J Biol Chem 300, 105499. 10.1016/j.jbc.2023.105499.

60. Smalakyte, D., Ruksenaite, A., Sasnauskas, G., Tamulaitiene, G., and Tamulaitis, G. (2024). Filament formation activates protease and ring nuclease activities of CRISPR Lon-SAVED. Molecular Cell 84, 4239–4255.e8. 10.1016/j.molcel.2024.09.002.

61. Tamulaitiene, G., Sabonis, D., Sasnauskas, G., Ruksenaite, A., Silanskas, A., Avraham, C., Ofir, G., Sorek, R., Zaremba, M., and Siksnys, V. (2024). Activation of Thoeris antiviral system via SIR2 effector filament assembly. Nature 627, 431–436. 10.1038/s41586-024-07092-x.

62. Hogrel, G., Guild, A., Graham, S., Rickman, H., Grüschow, S., Bertrand, Q., Spagnolo, L., and White, M.F. (2023). Author Correction: Cyclic nucleotide-induced helical structure activates a TIR immune effector. Nature 614, E15–E15. 10.1038/s41586-023-05701-9.

63. Morehouse, B.R., Yip, M.C.J., Keszei, A.F.A., McNamara-Bordewick, N.K., Shao, S., and Kranzusch, P.J. (2022). Cryo-EM structure of an active bacterial TIR–STING filament complex. Nature 608, 803–807. 10.1038/s41586-022-04999-1.

64. Tal, N., Morehouse, B.R., Millman, A., Stokar-Avihail, A., Avraham, C., Fedorenko, T., Yirmiya, E., Herbst, E., Brandis, A., Mehlman, T., et al. (2021). Cyclic CMP and cyclic UMP mediate bacterial immunity against phages. Cell 184, 5728–5739.e16. 10.1016/j.cell.2021.09.031.

65. Lowey, B., Whiteley, A.T., Keszei, A.F.A., Morehouse, B.R., Mathews, I.T., Antine, S.P., Cabrera, V.J., Kashin, D., Niemann, P., Jain, M., et al. (2020). CBASS Immunity Uses CARF-Related Effectors to Sense 3′–5′- and 2′–5′-Linked Cyclic Oligonucleotide Signals and Protect Bacteria from Phage Infection. Cell 182, 38–49.e17. 10.1016/j.cell.2020.05.019.

66. Polley, S., Lyumkis, D., and Horton, N.C. (2019). Mechanism of Filamentation-Induced Allosteric Activation of the SgrAI Endonuclease. Structure 27, 1497–1507.e3. 10.1016/j.str.2019.08.001.

67. Payne, L., Jackson, S., and Pinilla-Redondo, R. (2024). Supramolecular assemblies in bacterial immunity: an emerging paradigm. Trends in Microbiology 32, 828–831. 10.1016/j.tim.2024.06.003.

68. Vajjhala, P.R., Ve, T., Bentham, A., Stacey, K.J., and Kobe, B. (2017). The molecular mechanisms of signaling by cooperative assembly formation in innate immunity pathways. Molecular Immunology 86, 23– 37. 10.1016/j.molimm.2017.02.012.

69. Kagan, J.C., Magupalli, V.G., and Wu, H. (2014). SMOCs: supramolecular organizing centres that control innate immunity. Nat Rev Immunol 14, 821–826. 10.1038/nri3757.

70. Blattner, F.R., Plunkett, G., Bloch, C.A., Perna, N.T., Burland, V., Riley, M., Collado-Vides, J., Glasner, J.D., Rode, C.K., Mayhew, G.F., et al. (1997). The Complete Genome Sequence of *Escherichia coli* K-12. Science 277, 1453–1462. 10.1126/science.277.5331.1453.

71. Bhawsinghka, N., Glenn, K.F., and Schaaper, R.M. (2020). Complete Genome Sequence of Escherichia coli BL21-AI. Microbiol Resour Announc 9, e00009–20. 10.1128/MRA.00009-20.

72. Grant, S.G., Jessee, J., Bloom, F.R., and Hanahan, D. (1990). Differential plasmid rescue from transgenic mouse DNAs into Escherichia coli methylation-restriction mutants. Proc. Natl. Acad. Sci. U.S.A. 87, 4645– 4649. 10.1073/pnas.87.12.4645.

73. Truncaite, L., Šimoliūnas, E., Zajančkauskaite, A., Kaliniene, L., Mankevičiūte, R., Staniulis, J., Klausa, V., and Meškys, R. (2012). Bacteriophage vB_EcoM_FV3: a new member of “rV5-like viruses.” Arch Virol 157, 2431–2435. 10.1007/s00705-012-1449-x.

74. Alijošius, L., Šimoliūnas, E., Kaliniene, L., Meškys, R., and Truncaitė, L. (2017). Complete Genome Sequence of Escherichia coli Phage vB_EcoM_Alf5. Genome Announc 5, e00315–17. 10.1128/genomeA.00315-17.

75. Maffei, E., Shaidullina, A., Burkolter, M., Heyer, Y., Estermann, F., Druelle, V., Sauer, P., Willi, L., Michaelis, S., Hilbi, H., et al. (2021). Systematic exploration of Escherichia coli phage–host interactions with the BASEL phage collection. PLoS Biol 19, e3001424. 10.1371/journal.pbio.3001424.

76. Huang, C.-J., and Zheng, Y.-Y. (2019). Controlled Silanization Using Functional Silatrane for Thin and Homogeneous Antifouling Coatings. Langmuir 35, 1662–1671. 10.1021/acs.langmuir.8b01981.

77. Lyubchenko, Y.L., and Shlyakhtenko, L.S. (2009). AFM for analysis of structure and dynamics of DNA and protein–DNA complexes. Methods 47, 206–213. 10.1016/j.ymeth.2008.09.002.

78. Kupcinskaite, E., Tutkus, M., Kopūstas, A., Ašmontas, S., Jankunec, M., Zaremba, M., Tamulaitiene, G., and Sinkunas, T. (2022). Disarming of type I-F CRISPR-Cas surveillance complex by anti-CRISPR proteins AcrIF6 and AcrIF9. Sci Rep 12, 15548. 10.1038/s41598-022-19797-y.

79. Kopu̅stas, A., Ivanovaitė, Š., Rakickas, T., Pocevičiu̅tė, E., Paksaitė, J., Karvelis, T., Zaremba, M., Manakova, E., and Tutkus, M. (2021). Oriented Soft DNA Curtains for Single-Molecule Imaging. Langmuir 37, 3428–3437. 10.1021/acs.langmuir.1c00066.

80. Mirdita, M., Schütze, K., Moriwaki, Y., Heo, L., Ovchinnikov, S., and Steinegger, M. (2022). ColabFold: making protein folding accessible to all. Nat Methods 19, 679–682. 10.1038/s41592-022-01488-1.

81. Zhang, C., Shine, M., Pyle, A.M., and Zhang, Y. (2022). US-align: universal structure alignments of proteins, nucleic acids, and macromolecular complexes. Nat Methods 19, 1109–1115. 10.1038/s41592-022-01585-1.

82. Schubert, M., Lindgreen, S., and Orlando, L. (2016). AdapterRemoval v2: rapid adapter trimming, identification, and read merging. BMC Res Notes 9, 88. 10.1186/s13104-016-1900-2.

83. Quinlan, A.R. (2014). BEDTools: The Swiss-Army Tool for Genome Feature Analysis. CP in Bioinformatics 47. 10.1002/0471250953.bi1112s47.

84. Li, H., and Durbin, R. (2009). Fast and accurate short read alignment with Burrows–Wheeler transform. Bioinformatics 25, 1754–1760. 10.1093/bioinformatics/btp324.

85. Wang, L., Wang, S., and Li, W. (2012). RSeQC: quality control of RNA-seq experiments. Bioinformatics 28, 2184–2185. 10.1093/bioinformatics/bts356.

86. Danecek, P., Bonfield, J.K., Liddle, J., Marshall, J., Ohan, V., Pollard, M.O., Whitwham, A., Keane, T., McCarthy, S.A., Davies, R.M., et al. (2021). Twelve years of SAMtools and BCFtools. GigaScience 10, giab008. 10.1093/gigascience/giab008.

87. Crooks, G.E., Hon, G., Chandonia, J.-M., and Brenner, S.E. (2004). WebLogo: A Sequence Logo Generator: Figure 1. Genome Res. 14, 1188–1190. 10.1101/gr.849004.

88. Schindelin, J., Arganda-Carreras, I., Frise, E., Kaynig, V., Longair, M., Pietzsch, T., Preibisch, S., Rueden, C., Saalfeld, S., Schmid, B., et al. (2012). Fiji: an open-source platform for biological-image analysis. Nat Methods 9, 676–682. 10.1038/nmeth.2019.

89. Huisjes, N.M., Retzer, T.M., Scherr, M.J., Agarwal, R., Rajappa, L., Safaric, B., Minnen, A., and Duderstadt, K.E. (2022). Mars, a molecule archive suite for reproducible analysis and reporting of single-molecule properties from bioimages. eLife 11, e75899. 10.7554/eLife.75899.

90. D’Arcy, A., Villard, F., and Marsh, M. (2007). An automated microseed matrix-screening method for protein crystallization. Acta Crystallogr D Biol Crystallogr 63, 550–554. 10.1107/S0907444907007652.

91. Ireton, G.C., and Stoddard, B.L. (2004). Microseed matrix screening to improve crystals of yeast cytosine deaminase. Acta Crystallogr D Biol Crystallogr 60, 601–605. 10.1107/S0907444903029664.

92. Kabsch, W. (2010). XDS. Acta Crystallogr D Biol Crystallogr 66, 125–132. 10.1107/S0907444909047337.

93. Agirre, J., Atanasova, M., Bagdonas, H., Ballard, C.B., Baslé, A., Beilsten-Edmands, J., Borges, R.J., Brown, D.G., Burgos-Mármol, J.J., Berrisford, J.M., et al. (2023). The CCP4 suite: integrative software for macromolecular crystallography. Acta Crystallogr D Struct Biol 79, 449–461. 10.1107/S2059798323003595.

94. Panjikar, S., Parthasarathy, V., Lamzin, V.S., Weiss, M.S., and Tucker, P.A. (2005). Auto-rickshaw: an automated crystal structure determination platform as an efficient tool for the validation of an X-ray diffraction experiment. Acta Crystallogr D Biol Crystallogr 61, 449–457. 10.1107/S0907444905001307.

95. Rossmann, M.G., and Arnold, E. eds. (2006). International Tables for Crystallography: Crystallography of biological macromolecules 1st ed. (International Union of Crystallography) 10.1107/97809553602060000106.

96. Schneider, T.R., and Sheldrick, G.M. (2002). Substructure solution with SHELXD. Acta Crystallogr D Biol Crystallogr 58, 1772–1779. 10.1107/s0907444902011678.

97. Hao, Q. (2004). *ABS* : a program to determine absolute configuration and evaluate anomalous scatterer substructure. J Appl Crystallogr 37, 498–499. 10.1107/S0021889804008696.

98. Sheldrick, G.M. (2002). Macromolecular phasing with SHELXE. Zeitschrift für Kristallographie - Crystalline Materials 217, 644–650. 10.1524/zkri.217.12.644.20662.

99. Terwilliger, T.C. (2000). Maximum-likelihood density modification. Acta Crystallogr D Biol Crystallogr 56, 965–972. 10.1107/s0907444900005072.

100. Cowtan, K., and Main, P. (1998). Miscellaneous algorithms for density modification. Acta Crystallogr D Biol Crystallogr 54, 487–493. 10.1107/s0907444997011980.

101. Vagin, A., and Teplyakov, A. (2010). Molecular replacement with MOLREP. Acta Crystallogr D Biol Crystallogr 66, 22–25. 10.1107/S0907444909042589.

102. Emsley, P., Lohkamp, B., Scott, W.G., and Cowtan, K. (2010). Features and development of Coot. Acta Crystallogr D Biol Crystallogr 66, 486–501. 10.1107/S0907444910007493.

103. Liebschner, D., Afonine, P.V., Baker, M.L., Bunkóczi, G., Chen, V.B., Croll, T.I., Hintze, B., Hung, L.W., Jain, S., McCoy, A.J., et al. (2019). Macromolecular structure determination using X-rays, neutrons and electrons: recent developments in Phenix. Acta Crystallogr D Struct Biol 75, 861–877. 10.1107/S2059798319011471.

104. Murshudov, G.N., Vagin, A.A., and Dodson, E.J. (1997). Refinement of macromolecular structures by the maximum-likelihood method. Acta Crystallogr D Biol Crystallogr 53, 240–255. 10.1107/S0907444996012255.

105. Punjani, A., Rubinstein, J.L., Fleet, D.J., and Brubaker, M.A. (2017). cryoSPARC: algorithms for rapid unsupervised cryo-EM structure determination. Nat Methods 14, 290–296. 10.1038/nmeth.4169.

106. Grant, T., Rohou, A., and Grigorieff, N. (2018). cisTEM, user-friendly software for single-particle image processing. Elife 7, e35383. 10.7554/eLife.35383.

107. Afonine, P.V., Poon, B.K., Read, R.J., Sobolev, O.V., Terwilliger, T.C., Urzhumtsev, A., and Adams, P.D. (2018). Real-space refinement in PHENIX for cryo-EM and crystallography. Acta Crystallogr D Struct Biol 74, 531–544. 10.1107/S2059798318006551.

108. Kabsch, W., and Sander, C. (1983). Dictionary of protein secondary structure: Pattern recognition of hydrogen-bonded and geometrical features. Biopolymers 22, 2577–2637. 10.1002/bip.360221211.

109. Meng, E.C., Goddard, T.D., Pettersen, E.F., Couch, G.S., Pearson, Z.J., Morris, J.H., and Ferrin, T.E. (2023). UCSF C_HIMERA_X : Tools for structure building and analysis. Protein Science 32, e4792. 10.1002/pro.4792.

110. Pedregosa, F., Varoquaux, G., Gramfort, A., Michel, V., Thirion, B., Grisel, O., Blondel, M., Prettenhofer, P., Weiss, R., Dubourg, V., et al. (2011). Scikit-learn: Machine Learning in Python. Journal of Machine Learning Research 12, 2825–2830.

111. Waskom, M. (2021). seaborn: statistical data visualization. JOSS 6, 3021. 10.21105/joss.03021.

112. Holm, L., Laiho, A., Törönen, P., and Salgado, M. (2023). DALI shines a light on remote homologs: One hundred discoveries. Protein Science 32, e4519. 10.1002/pro.4519.

113. Olechnovič, K., Kulberkytė, E., and Venclovas, Č. (2013). CAD-score: A new contact area difference-based function for evaluation of protein structural models. Proteins 81, 149–162. 10.1002/prot.24172.

114. Kunzmann, P., Müller, T.D., Greil, M., Krumbach, J.H., Anter, J.M., Bauer, D., Islam, F., and Hamacher, K. (2023). Biotite: new tools for a versatile Python bioinformatics library. BMC Bioinformatics 24, 236. 10.1186/s12859-023-05345-6.

115. Chen, E.A., and Porter, L.L. (2023). SSD_RAW_ : S_OFTWARE_ for generating comparative protein secondary structure diagrams. Protein Science 32, e4836. 10.1002/pro.4836.

116. Olechnovič, K., and Venclovas, Č. (2021). VoroContacts: a tool for the analysis of interatomic contacts in macromolecular structures. Bioinformatics 37, 4873–4875. 10.1093/bioinformatics/btab448.

117. Eisenstein, M. (2021). Microscopy made to order. Nat Methods 18, 1277–1281. 10.1038/s41592-021-01313-1.

118. Alsamsam, M.N., Kopūstas, A., Jurevičiūtė, M., and Tutkus, M. (2022). The miEye: Bench-top super-resolution microscope with cost-effective equipment. HardwareX 12, e00368. 10.1016/j.ohx.2022.e00368.

119. Shin, E., Kim, W., Lee, S., Bae, J., Kim, S., Ko, W., Seo, H.S., Lim, S., Lee, H.S., and Jo, K. (2019). Truncated TALE-FP as DNA Staining Dye in a High-salt Buffer. Sci Rep 9, 17197. 10.1038/s41598-019-53722-0.

